# *Scn1a*-GFP transgenic mouse revealed Nav1.1 expression in neocortical pyramidal tract projection neurons

**DOI:** 10.1101/2021.03.31.437794

**Authors:** Tetsushi Yamagata, Ikuo Ogiwara, Tetsuya Tatsukawa, Toshimitsu Suzuki, Yuka Otsuka, Nao Imaeda, Emi Mazaki, Ikuyo Inoue, Natsuko Tokonami, Yurina Hibi, Shigeyoshi Itohara, Kazuhiro Yamakawa

## Abstract

Expressions of voltage-gated sodium channels Nav1.1 and Nav1.2, encoded by *SCN1A* and *SCN2A* genes respectively, have been reported to be mutually exclusive in most brain regions. In juvenile and adult neocortex, Nav1.1 is predominantly expressed in inhibitory neurons while Nav1.2 is in excitatory neurons. Although a distinct subpopulation of layer V (L5) neocortical excitatory neurons were also reported to express Nav1.1, their nature has been uncharacterized. In hippocampus, Nav1.1 has been proposed to be expressed only in inhibitory neurons. By using newly-generated transgenic mouse lines expressing *Scn1a* promoter-driven green fluorescent protein (GFP), here we confirm the mutually-exclusive expressions of Nav1.1 and Nav1.2 and the absence of Nav1.1 in hippocampal excitatory neurons. We also show that Nav1.1 is expressed in inhibitory and a subpopulation of excitatory neurons not only in L5 but all layers of neocortex. By using neocortical excitatory projection neuron markers including FEZF2 for L5 pyramidal tract (PT) and TBR1 for layer VI (L6) cortico-thalamic (CT) projection neurons, we further show that most L5 PT neurons and a minor subpopulation of layer II/III (L2/3) cortico-cortical (CC) neurons express Nav1.1 while the majority of L6 CT, L5/6 cortico-striatal (CS) and L2/3 CC neurons express Nav1.2. These observations now contribute to the elucidation of pathological neural circuits for diseases such as epilepsies and neurodevelopmental disorders caused by *SCN1A* and *SCN2A* mutations.

## Introduction

Voltage-gated sodium channels (VGSCs) play crucial roles in the generation and propagation of action potentials, contributing to excitability and information processing (Catterall, 2012). They consist of one main pore-forming alpha- and one or two subsidiary beta-subunits that regulate kinetics or subcellular trafficking of the alpha subunits. Human has nine alpha (Nav1.1∼Nav1.9) and four beta (beta-1∼beta-4) subunits. Among alphas, Nav1.1, Nav1.2, Nav1.3 and Nav1.6, encoded by *SCN1A*, *SCN2A*, *SCN3A* and *SCN8A*, respectively, are expressed in central nervous system. *SCN3A* is mainly expressed embryonically (Brysch et al., 1991), and *SCN1A*, *SCN2A* and *SCN8A* are major alphas after birth. Although these three genes show mutations in a wide spectrum of neurological diseases such as epilepsy, autism spectrum disorder (ASD) and intellectual disability, two of those, *SCN1A* and *SCN2A,* are major ones (reviewed in Yamakawa et al., 2016; Meisler et al., 2021). To understand the circuit basis of these diseases, it’s indispensable to know the detailed distributions of these molecules in the brain.

We previously reported that expressions of Nav1.1 and Nav1.2 seem to be mutually-exclusive in many brain regions (Yamagata et al., 2017). In adult neocortex and hippocampus, Nav1.1 is dominantly expressed in medial ganglionic eminence-derived parvalbumin-positive (PV-IN) and somatostatin-positive (SST-IN) inhibitory neurons (Ogiwara et al., 2007; Lorincz and Nusser, 2008; Ogiwara et al., 2013; Li et al., 2014; Tai et al., 2014; Tian et al., 2014; Yamagata et al, 2017). In the neocortex, some amount of Nav1.1 is also expressed in a distinct subset of layer V (L5) excitatory neurons (Ogiwara et al., 2013), but their natures were unknown. In the hippocampus, Nav1.1 seems to be expressed in inhibitory but not in excitatory neurons (Ogiwara et al., 2007; Ogiwara et al., 2013). In contrast, a major amount of Nav1.2 (∼95%) is expressed in excitatory neurons including the most of neocortical and all of hippocampal ones, and a minor amount is expressed in caudal ganglionic eminence-derived inhibitory neurons such as vasoactive intestinal polypeptide (VIP)-positive ones (Lorincz and Nusser 2010; Yamagata et al., 2017; Ogiwara et al., 2018). However, a recent study reported that a subpopulation (more than half) of VIP-positive inhibitory neurons is Nav1.1-positive (Goff and Goldberg, 2019).

VGSCs are mainly localized at axons and therefore it is not always easy to identify their origins, soma. To overcome this, here in this study we generated bacterial artificial chromosome (BAC) transgenic mouse lines that express GFP under the control of *Scn1a* promoters, and we carefully investigated the GFP/Nav1.1 distribution in mouse brain. Our analysis confirmed that expressions of Nav1.1 and Nav1.2 are mutually-exclusive and that in neocortex Nav1.1 is expressed in both inhibitory and excitatory neurons while in hippocampus only in inhibitory but totally absent in excitatory neurons. Furthermore by using a transcription factor FEZF2 (FEZ family zinc finger protein 2 transcriptional factor), also referred to as Fezl, Fez1, Zfp312 and Fez, as a marker for L5 PT neurons (Inoue et al., 2004; Chen et al., 2005; Chen et al., 2008; Molyneaux et al., 2005; Lodato et al., 2014 ; Matho et al., 2021), and a transcription factor TBR1 which suppresses FEZF2 expression and therefore does not overlap with FEZF2 (Han et al., 2011; McKenna et al., 2011, Matho et al., 2021), we found that most of L5 FEZF2-positive neurons are GFP-positive while L5/6 TBR1-positive neurons are largely GFP-negative and Nav1.2-positive. These results proposed that Nav1.1 is expressed in L5 PT while Nav1.2 in L5/6 non-PT neurons such as L5/6 CS and L6 CT projection neurons. A majority of L2/3 excitatory neurons express Nav1.2 but a minor subpopulation are GFP-positive, suggesting that most of CC projection neurons express Nav1.2 but the distinct minor population express Nav1.1. These results refine the expression loci of Nav1.1 and Nav1.2 in the brain and should contribute to the understanding of circuit mechanisms for diseases caused by *SCN1A* and *SCN2A* mutations.

## Results

### Generation and verification of *Scn1a-*GFP transgenic mouse lines

*Scn1a*-GFP founder mice were generated from C57BL/6J zygotes microinjected with a modified *Scn1a*-GFP BAC construct harboring all, upstream and downstream, *Scn1a*-promoters (Nakayama et al., 2010) (Figure 1A) (see Materials and Methods for details). Western blot analysis (Figure 1B) and immunohistochemistry (Supplementary figure S1) showed robust GFP expression and mostly normal expression levels of Nav1.1 in *Scn1a*-GFP mouse lines #184 and #233. Both lines showed a similar distribution of chromogenic GFP immunosignals across the entire brain (Figures 2A-H), and a similar distribution was also obtained in fluorescence detection of GFP (Figures 2I-L and Supplementary figure S2). In neocortex (Figures 2B, F, J and Supplementary figures S2B, F), GFP-positive cells were distributed throughout all cortical layers. In hippocampus (Figures 2C, G, K and Supplementary figures S2C, G), cells with intense GFP signals, which are assumed to be PV-IN and SST-IN (Ogiwara et al, 2007; Tai et al., 2014) (see also Figure 8), were scattered in stratum oriens, pyramidale, radiatum, lucidum and lacunosum-moleculare of the CA (cornu ammonis) fields, hilus and molecular layer of dentate gyrus. Of note, somata of dentate granule cells were apparently GFP-negative. CA1∼3 pyramidal cells were twined around with fibrous GFP immunosignals. We previously reported that the fibrous Nav1.1-signals clinging to somata of hippocampal CA1∼3 pyramidal cells were disappeared by conditional elimination of Nav1.1 in PV-INs but not in excitatory neurons, and therefore concluded that these Nav1.1-immunopositive fibers are axon terminals of PV-INs (Ogiwara et al., 2013). As such, GFP signals are fibrous but do not form cell shapes in the CA pyramidal cell layer (Figures 2C, G, K and Supplementary figures S2C, G), and therefore these CA pyramidal cells themselves are assumed to be GFP-negative. These observations further confirmed our previous proposal that hippocampal excitatory neurons are negative for Nav1.1 (Ogiwara et al., 2007; Ogiwara et al., 2013). In cerebellum (Figures 2D, H, L and Supplementary figures S2D, H), GFP signals appeared in Purkinje, basket, and deep cerebellar nuclei cells, again consistent to the previous reports (Ogiwara et al., 2007; Ogiwara et al., 2013). In the following analyses, we used the line #233 which shows stronger GFP signals than #184.

**Figure 1.**
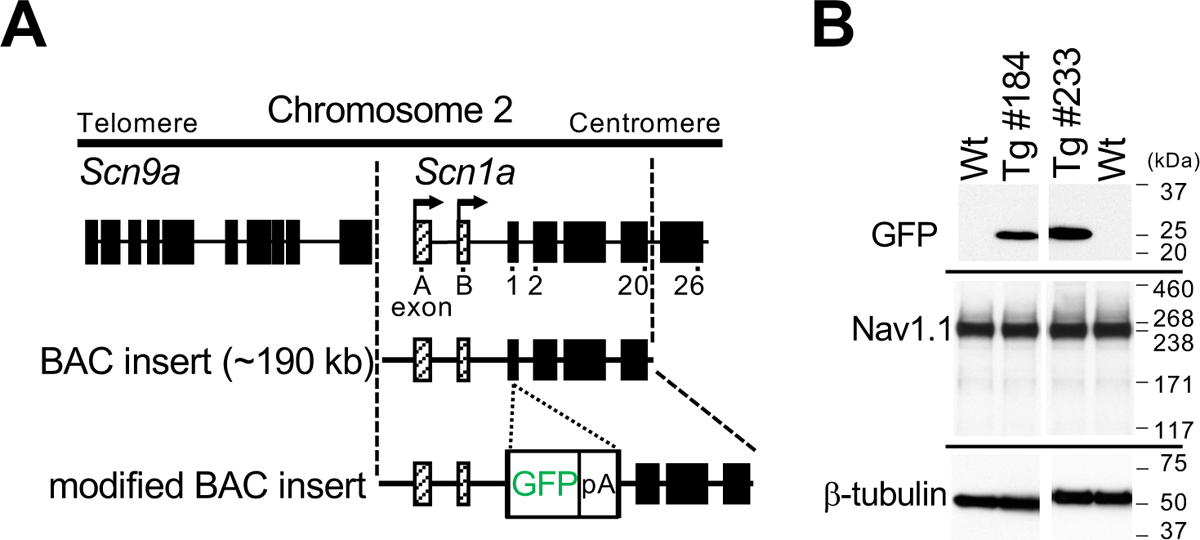
Generation of *Scn1a*-GFP mice. (**A**) Schematic representation of the modified BAC construct containing the *Scn1a*-GFP transgene. A GFP reporter cassette consisting of GFP cDNA and a polyadenylation signal was inserted at the ATG initiation codon in the coding exon 1 of *Scn1a*. Filled and hatched boxes indicate the coding and non-coding exons of *Scn9a* and *Scn1a*. Arrows indicate the start sites and orientation of transcription of *Scn1a*. (**B**) Western blot analysis for *Scn1a*-GFP and endogenous Nav1.1. The whole cytosolic fractions from 5W *Scn1a*-GFP brains (lines #184 and #233) were probed with anti-GFP and their membrane fractions were probed with anti-Nav1.1 antibodies. β-Tubulin was used as an internal control. pA, polyadenylation signal; Tg, hemizygous *Scn1a*-GFP transgenic mice; Wt, wild-type littermates.

**Figure 2.**
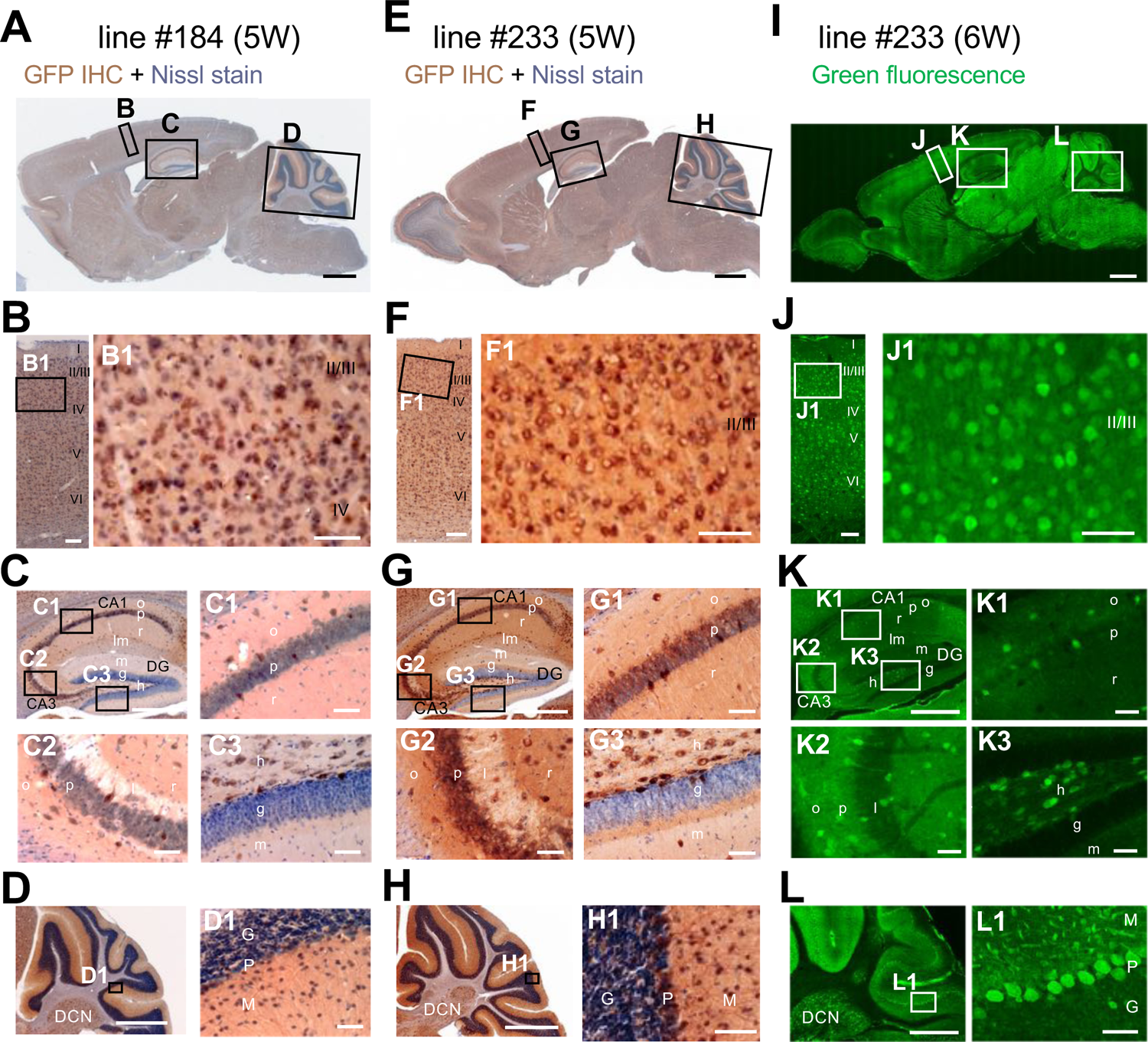
Distributions of GFP signals in brains are similar among *Scn1a*-GFP mouse lines. Chromogenic immunostaining of GFP (brown) with Nissl counterstaining (violet) of lines #184 and #233 (A-H) and GFP fluorescence images of line #233 (I-L) on parasagittal sections from 5–6W *Scn1a*-GFP brains. Boxed areas in (A, E, I, B, F, J, C, G, K, D, H, L) are magnified in (B-D, F-H, J-L, B1, F1, J1, C1-3, G1-3, K1-3, D1, H1, L1). The two lines (lines #184 and #233) showed a similar distribution pattern of GFP-expressing cells across all brain regions (A-H), but the signals in the line #233 are more intense than the line #184. In neocortex (B, F, J), GFP-expressing cells were scattered throughout the entire region. In the hippocampus (C, G, K), GFP-positive inhibitory neurons were sparsely distributed (see also Figure 8), while excitatory neurons in stratum pyramidale and stratum granulosum are GFP-negative. In cerebellum (D, H, L), Purkinje, basket, and deep cerebellar nuclei cells were GFP-positive. IHC, immunohistochemistry; CA, cornu ammonis; DG, dentate gyrus; o, stratum oriens; p, stratum pyramidale; r, stratum radiatum; lm, stratum lacunosum-moleculare; l, stratum lucidum; m, stratum moleculare; g, stratum granulosum; h, hilus; DCN, deep cerebellar nuclei; M, molecular layer; P, Purkinje cell layer; G, granular cell layer. Scale bars; 1 mm (A, E, I), 500 µm (C, D, G, H, K, L), 100 µm (B, F, J), 50 µm (B1, C1-3, D1, F1, G1-3, H1, J1, K1-3, L1).

Quantification of Nav1.1 signals in western blot analyses of brain lysates from the *Scn1a*-GFP mice and their wild-type littermates (N = 5 animals per each genotype) showed no difference between genotypes, while that of GFP somehow deviated among individual *Scn1a*-GFP mice (Supplementary figure S3). Fluorescence imaging of the *Scn1a*-GFP sagittal brain sections at postnatal day 15 (P15), 4-week-old (4W) and 8W showed that GFP-signals continue to be intense in caudal region such as thalamus, mid brain, and brainstem (Figure 3), which is well consistent with our previous report of Nav1.1 protein and *Scn1a* mRNA distributions in wild-type mouse brain (Ogiwara et al, 2007).

**Figure 3.**
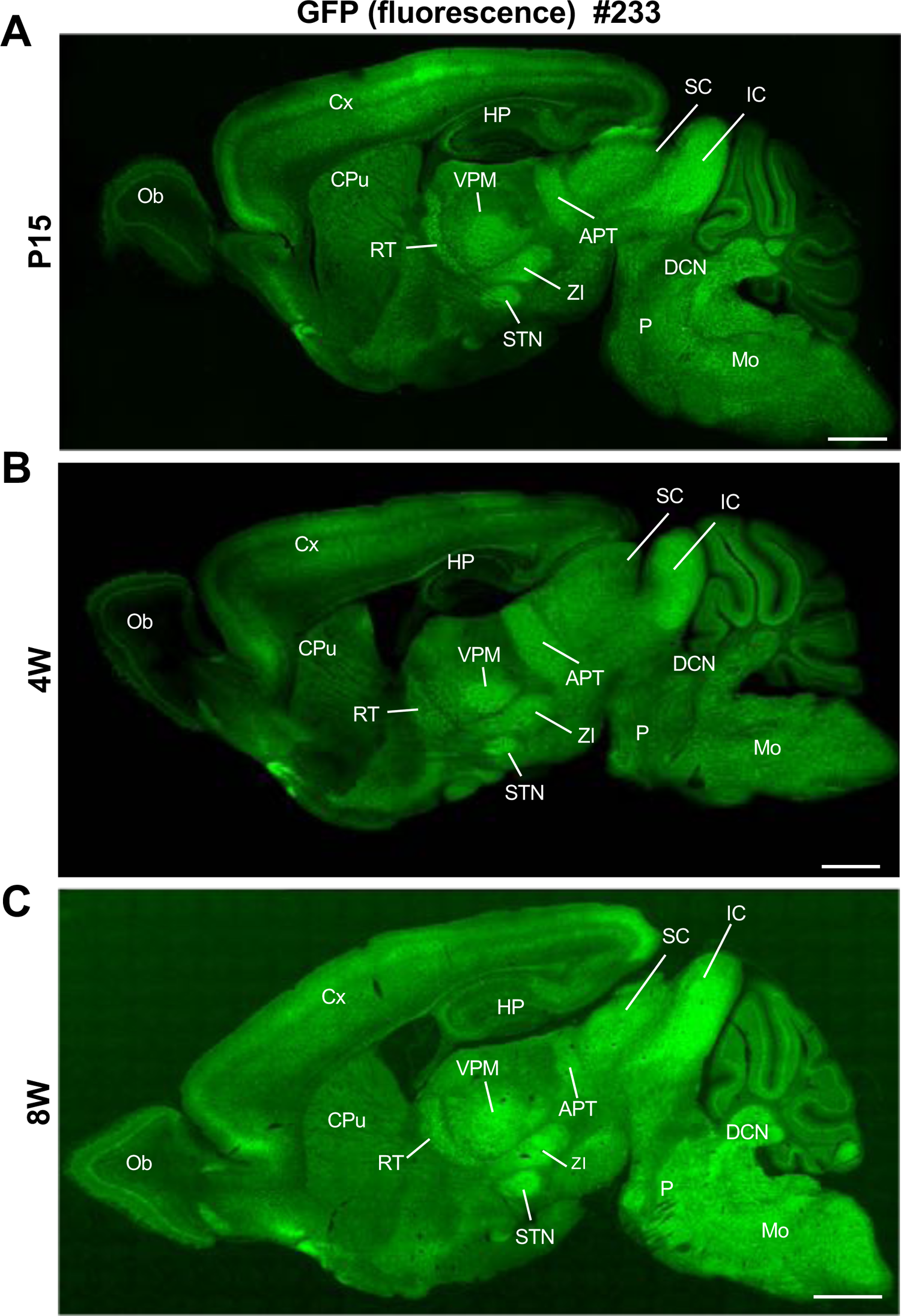
Distribution of GFP signals in *Scn1a*-GFP mouse brain are largely maintained through development. Fluorescent images of parasagittal sections from P15 (**A**), 4W (**B**) and 8W (**C**) *Scn1a*-GFP mouse brains (line #233). GFP signals were observed in multiple brain regions. APT, anterior pretectal nucleus; CPu, caudate putamen; Cx, cerebral cortex; DCN, deep cerebellar nuclei; HP, hippocampus; IC, inferior colliculus; Mo, medulla oblongata; Ob, olfactory bulb; P, pons; RT, reticular thalamic nucleus; SC, superior colliculus; STN, subthalamic nucleus; VPM, ventral posteromedial thalamic nucleus; ZI, zona incerta. Scale bars; 1 mm.

### Nav1.1 is expressed in both excitatory and inhibitory neurons in neocortex but only in inhibitory neurons in hippocampus

In the neocortex of *Scn1a*-GFP mouse, a large number of cells with GFP-positive somata (GFP-positive cells) were broadly distributed across all cortical layers (Figure 3 and Supplementary figure S4). Intensities of GFP signals in primary somatosensory cortex (S1) at L2/3 are much higher than other areas such as primary motor cortex (M1) (Supplementary figure S4), however the cell population (density) of GFP-positive cells did not differ in these areas indicating that GFP-signals for GFP-positive cells are stronger in S1 at L2/3. Although GFP signals are strong in PV-INs (see Figure 8), cell density of PV-INs is not specifically high at S1 area and therefore most cells with strong GFP signals in S1 at L2/3 may not be PV-INs but excitatory neurons.

In order to know the ratio of GFP-positive cells among all neurons, we further performed immunohistochemical staining using NeuN-antibody on *Scn1a*-GFP mouse at P15 and cells were counted at M1 and S1 (Supplementary figure S5 and Supplementary table S1). The NeuN staining showed that GFP-positive cells occupy 30% (L2/3), 32% (L5) and 22% (L6) of NeuN- or GFP-positive cells at P15 (Supplementary figure S5B and Supplementary table S1). However, we noticed that sparsely-distributed cells with intense GFP-signals, which are assumed to be PV-INs (see Figure 8), were often NeuN-negative (Supplementary figure S5A - arrowheads), reminiscent of a previous report that NeuN expression is absent in cerebellar inhibitory neurons such as Golgi, basket and satellite cells in cerebellum (Weyer et al., 2003). Therefore, NeuN-positive cells do not represent all neurons in neocortex as well. NeuN/GFP-double negative neurons could even exist and therefore above figure (Supplementary figure S5B) may deviate from the real ratios of GFP-positive cells among all neurons.

Next, we performed triple-immunostaining of Nav1.1, GFP and ankyrinG on brains of *Scn1a*-GFP mouse at P15. In the neocortex (Figure 4), axon initial segments (AISs) of cells with Nav1.1-positive somata were always Nav1.1-positive but somata of cells with Nav1.1-positive AISs were occasionally Nav1.1-negative (Figures 4A-C). Cell counting revealed that 17% (L2/3), 21% (L5) and 8% (L6) of neurons (cells with ankyrinG-positive AISs) were GFP-positive (Figure 4D-left panel and Supplementary table S2). Of note, all cells with Nav1.1-positive AISs or somata were GFP-positive, but AISs or somata for only half of GFP-positive cells were Nav1.1-positive (Figure 4D and Supplementary tables S3, S4), possibly due to undetectably low levels of Nav1.1 immunosignals in a subpopulation of GFP-positive cells. The above ratios of GFP-positive cells among neurons (cells with ankyrinG-positive AISs) obtained in the triple-immunostaining of Nav1.1, GFP and ankyrinG are rather discordant to those obtained in the later experiment of triple-immunostaining of Nav1.2, GFP and ankyrinG, 23% (L2/3), 30% (L5) and 21% (L6) (see Figure 11). Therefore, we additionally performed double-immunostaining of GFP and ankyrinG on brains of *Scn1a*-GFP mouse at P15, and the ratios of of GFP-positive cells among neurons were 30% (L2/3), 26% (L5) and 9% (L6) (Supplementary figure S6 and Supplementary table S5). Averaged ratios of GFP-positive cells among neurons of Figures 4D, 11B and Supplementary figure S6B are 23% (L2/3), 26% (L5) and 13% (L6) (Supplementary figure S7 and Supplementary table S6), which are actually significantly lower than those obtained in the NeuN-staining (Supplementary figure S5 and Supplementary table S1).

**Figure 4.**
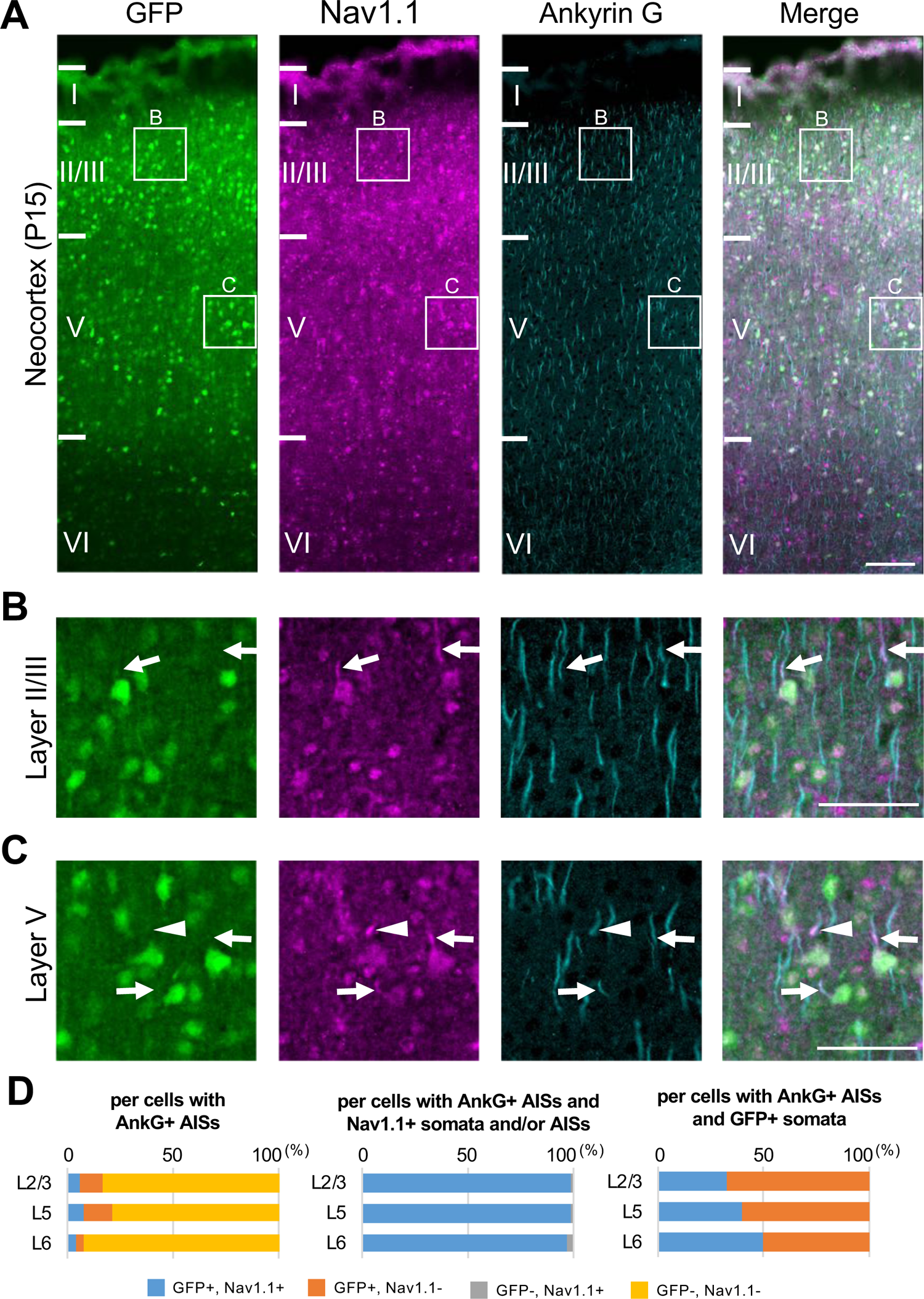
All cells with Nav1.1-positive AISs are GFP-positive but only half or less population of AISs for GFP-positive cells are Nav1.1-positive in *Scn1a*-GFP mouse neocortex. (**A**) Triple immunofluorescent staining of parasagittal section from P15 *Scn1a*-GFP mouse brain (line #233) by mouse anti-GFP (green), rabbit anti-Nav1.1 (magenta), and goat anti-ankyrinG (cyan) antibodies. Regions at primary motor cortex are shown. (**B**, **C**) Magnified images outlined in (A) are shown in (B) and (C). Arrows indicate AISs of cells with GFP-positive somata in which both somata and AISs are positive for Nav1.1. Arrowheads indicate AISs of cells with GFP-positive somata in which AISs but not somata are positive for Nav1.1. All images are oriented from pial surface (top) to callosal (bottom). m (A), 50 μm (B and C). (**D**) Cell counting of three *Scn1a*-GFP mice. Bar graphs indicating the percentage of cells with GFP- and Nav1.1-positive/negative somata and AISs per cells with ankyrinG-positive AISs (left panel), the percentage of cells with GFP-positive/negative somata per cells with ankyrinG-positive AISs and Nav1.1-positive somata and/or AISs (middle panel), and the percentage of cells with Nav1.1-positive/negative somata and/or AISs per cells with ankyrinG-positive AISs and GFP-positive somata (right panel) in L2/3, L5, and L6 (see also Supplementary tables S2-S4). Only cells with ankyrinG-positive AISs were counted. Nav1.1 immunosignals were occasionally observed in somata, but in such cases Nav1.1 signals were always observed in their AISs if visible by ankyrinG staining. Note that 99% (L2/3), 99% (L5) and 97% (L6) of cells with Nav1.1-positive AISs have GFP-positive somata (middle panel), but only half or less of cells with GFP-positive somata have Nav1.1-positive AISs (right panel). L2/3, L5: neocortical layer II/III and V. AnkG, ankyrinG; +, positive; -, negative.

In contrast to the neocortex where only half of GFP-positive cells were Nav1.1-positive, in the hippocampus all GFP-positive cells were Nav1.1-positive and all Nav1.1-positive cells were GFP-positive (Figure 5). Actually, most of excitatory neurons such as CA1∼3 pyramidal cells and dentate granule cells were GFP-negative. As described above (Figures 2C, G), fibrous GFP and Nav1.1 signals twining around CA1∼3 pyramidal cells’ somata which are assumed to be axon terminals of PV-INs were again observed (Figures 5A, B, D). Cell counting in the hippocampal CA1 region showed that 98% of cells with GFP-positive somata were Nav1.1-positive at their AISs and 100% of cells with Nav1.1-positive AISs were GFP-positive (Figure 5F and Supplementary tables S7, S8).

**Figure 5.**
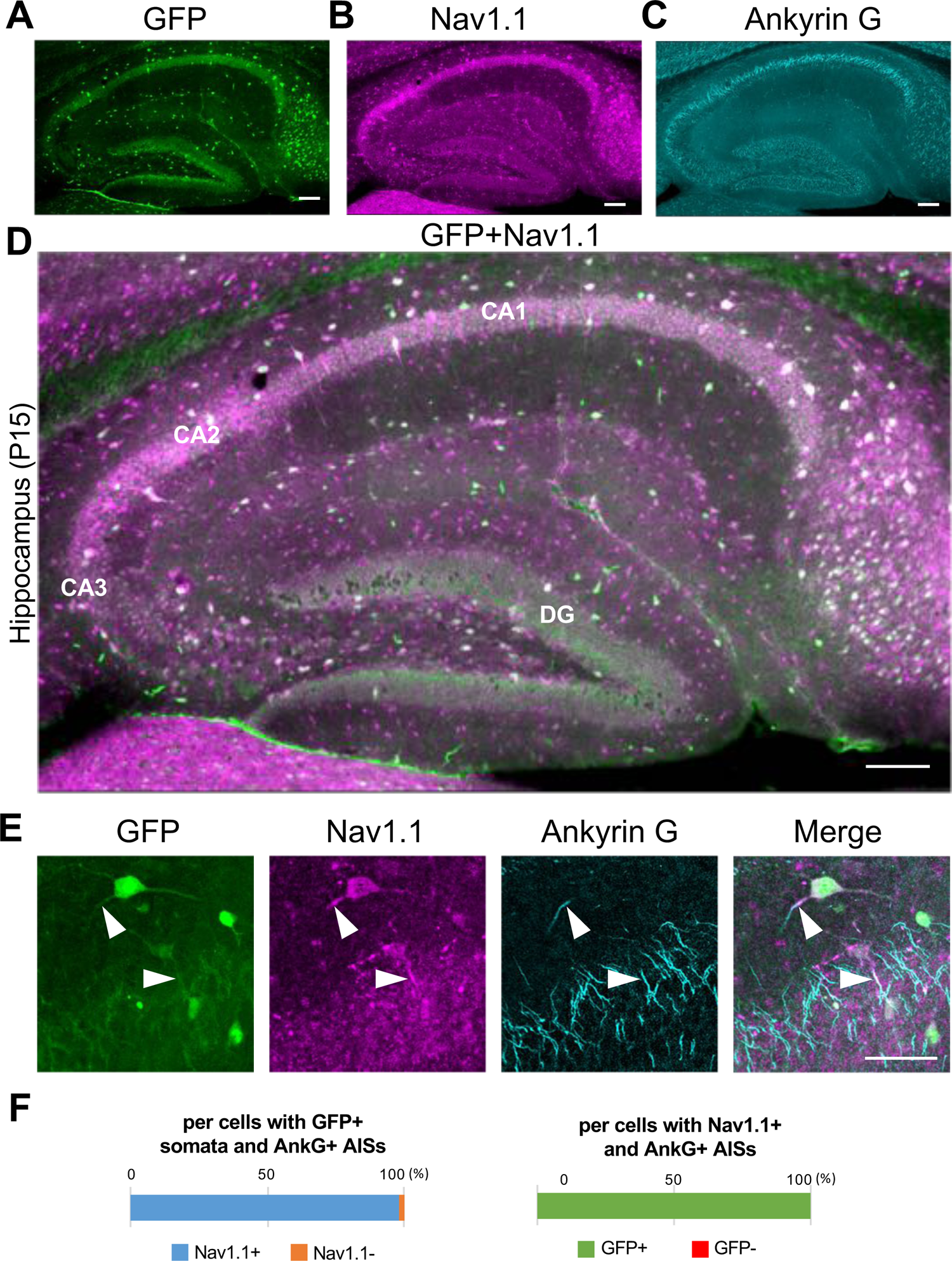
All cells with Nav1.1-positive AISs are GFP-positive and all AISs of GFP-positive cells are Nav1.1-positive in *Scn1a*-GFP mouse hippocampus. (**A**-**D**) Triple immunofluorescent staining of parasagittal section from P15 *Scn1a*-GFP mouse brain (line #233) by mouse anti-GFP (green), rabbit anti-Nav1.1 (magenta), and goat anti-ankyrinG (cyan) antibodies. Regions at hippocampus were shown. Note that GFP and Nav1.1 immunosignals mostly overlap at somata. CA1, cornu ammonis 1; CA2, cornu ammonis 2; CA3, cornu ammonis 3; DG, dentate gyrus. Images are oriented from pial surface (top) to callosal (bottom). Scale bars; 100 μm. (**E**) Magnified images for co-expression of GFP and Nav1.1 in cells at CA1 region. Arrowheads indicate Nav1.1-positive AISs GFP-expression cells. Scale bar; 50 μm. (**F**) Bar graphs indicate the percentage of cells in hippocampal CA1 region with Nav1.1-positive/negative AISs per cells with GFP-positive somata and ankyrinG-positive AISs (left panel), and the percentage of cells with GFP-positive/negative somata per cells with Nav1.1/ankyrinG-double positive AISs (right panel) (see also Supplementary tables S7, S8). Only cells with ankyrinG-positive AISs were counted. GFP/Nav1.1-double negative cells, most of which are pyramidal cells, were not counted because of the accumulated nature of their ankyrinG-positive AISs. AnkG, ankyrinG; +, positive; -, negative.

Double *in situ* hybridization of *Scn1a* and GFP mRNAs showed that these signals well overlap in both neocortex and hippocampus of *Scn1a*-GFP mice (Figure 6), further supporting that the GFP signals well represent endogenous *Scn1a*/Nav1.1 expression. Again, in neocortex *Scn1a* and GFP mRNAs seem to be expressed in a number of neurons including some of excitatory pyramidal cells, while in hippocampus they are absent in excitatory neurons such as CA1∼3 pyramidal cells and dentate granule cells. All of these distributions of *Scn1a* and GFP mRNAs in *Scn1a*-GFP transgenic mouse brain are consistent to our previous report of regional distributions of *Scn1a* mRNA in wild-type mouse (Ogiwara et al, 2007).

**Figure 6.**
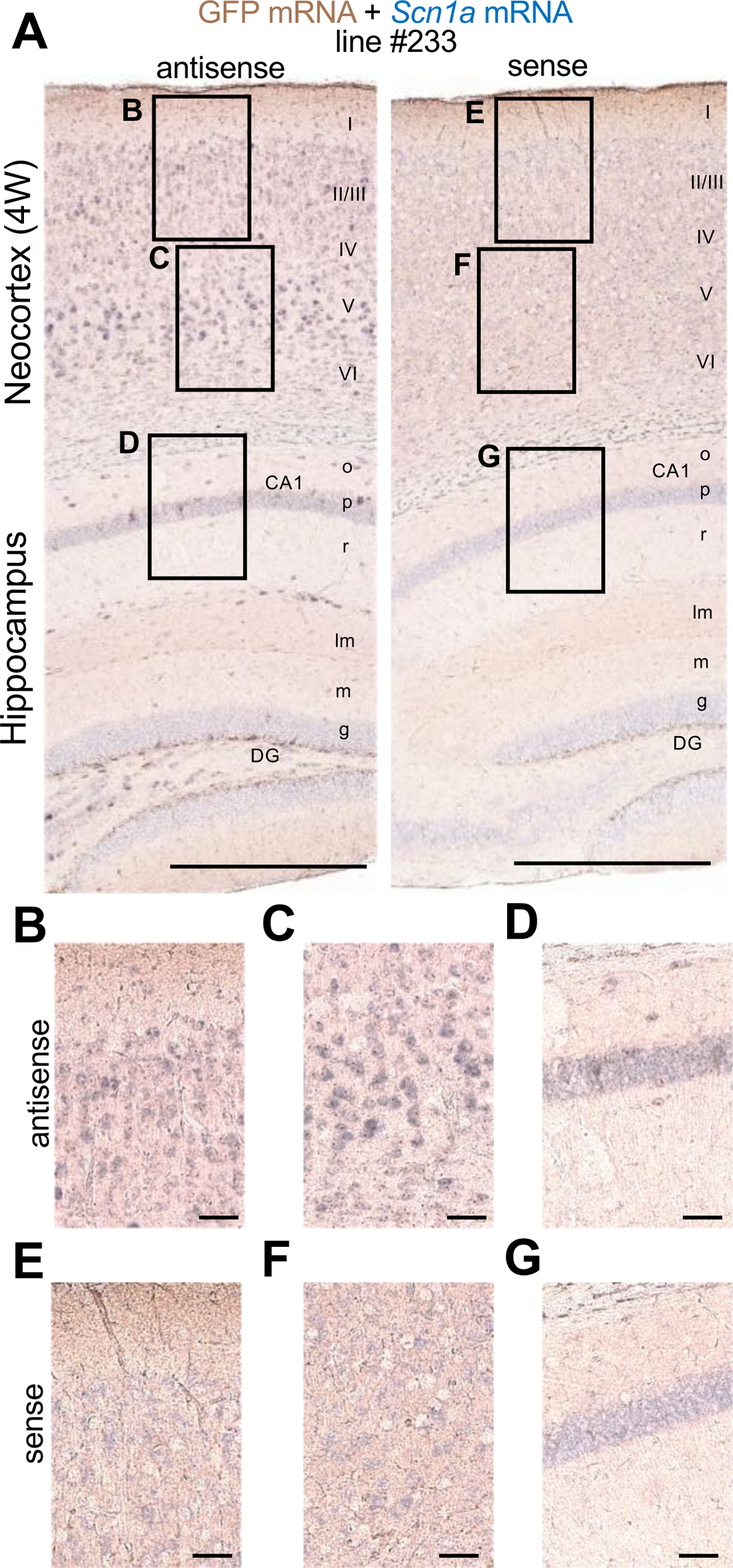
GFP and *Scn1a* mRNAs expressions goodly overlap in *Scn1a*-GFP mouse brain. Double *in situ* hybridization for *Scn1a*-GFP transgene mRNA and endogenous *Scn1a* mRNA on parasagittal sections from 4W *Scn1a*-GFP brains (line #233). (**A**) Sections were hybridized with antisense (left) and sense (right) RNA probes for GFP transgene (brown) and endogenous *Scn1a* (blue) mRNA species and chromogenically stained. Magnified images outlined in (A) are shown in (**B-D**) for antisense probes, and (**E-G**) for sense probes. o, stratum oriens; p, stratum pyramidale; r, stratum radiatum; lm, stratum lacunosum-moleculare; m, stratum moleculare; g, stratum granulosum, CA1, cornu ammonis 1; DG, dentate gyrus. Scale bars; 500 µm (A), 50 µm (B-G).

To investigate the ratio of inhibitory neurons in GFP-positive cells, we generated and examined *Scn1a*-GFP and vesicular GABA transporter (*Vgat*)-Cre (Ogiwara et al., 2013) double transgenic mice in which *Vgat*-Cre is expressed in all GABAergic inhibitory neurons and visualized by floxed tdTomato transgene (Figure 7). In the neocortex at 4W, 23% (L2/3), 28% (L5) and 27% (L6) of GFP-positive cells were Tomato-positive inhibitory neurons and 73% (L2/3), 77% (L5) and 83% (L6) of Tomato-positive cells were GFP-positive (Figure 7C and Supplementary tables S9, S10). These results suggest that a significant subpopulation of neocortical excitatory neurons also express Nav1.1. Our previous observation that Nav1.1 is expressed in callosal axons of neocortical excitatory neurons (Ogiwara et al., 2013) supports that a subpopulation of L2/3 CC neurons express Nav1.1. Unlike in neocortex, in the hippocampus most of GFP-positive cells were Tomato-positive, 98% (CA1) and 94% (DG), and majorities of Tomato-positive GABAergic neurons are GFP-positive, 93% (CA1) and 77% (DG). These results further confirmed that in hippocampus Nav1.1 is expressed in inhibitory neurons but not in excitatory neurons. Although somata of pyramidal cells in CA2/3 region are weakly GFP-positive in this and some other experiments (Figure 7B, Supplementary figure S2G), those were GFP-negative in other experiments (Figures 2K, 5A, D) and therefore the Nav1.1 expression in CA2/3 pyramidal cells would be minimal if any.

**Figure 7.**
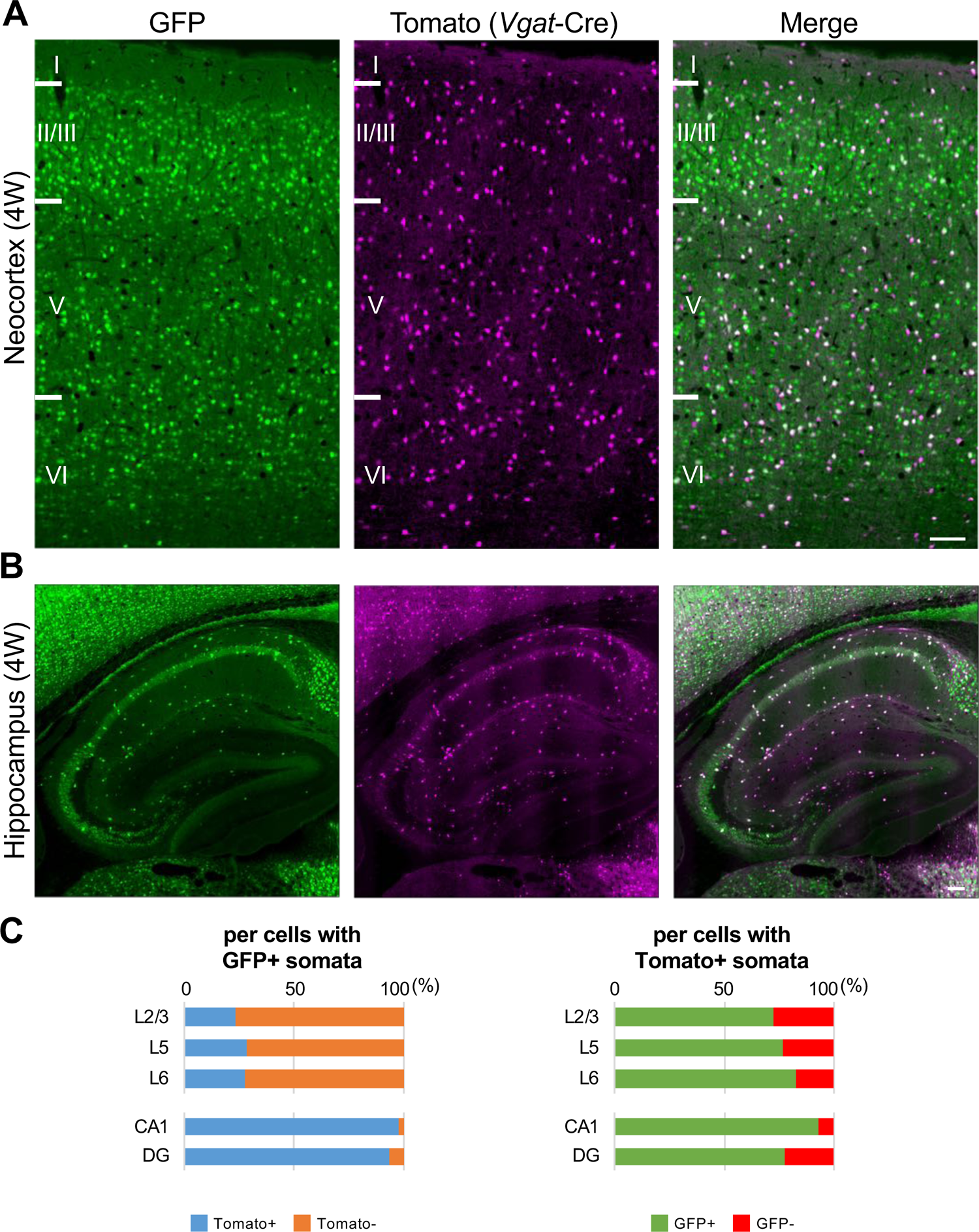
One third of GFP-positive cells in neocortex are inhibitory neurons, but most of GFP-positive cells in hippocampus are inhibitory neurons. (**A, B**) GFP (green) and Tomato (magenta) fluorescent images of parasagittal sections from 4W *Scn1a*-GFP/*Vgat*-cre/*Rosa26*-tdTomato mouse. Regions at primary motor cortex (A) and hippocampus (B) are shown. Scale bar; 100 μm. (**C**) Bar graphs indicate the percentage of cells with Tomato-positive/negative somata per cells with GFP-positive somata (left panel) (see also Supplementary table S9) and the percentage of cells with GFP-positive/negative somata per cells with Tomato-positive somata (right panel) (see also Supplementary table S10) in L2/3, L5, L6, CA1 and DG. Cells in primary motor cortex and hippocampus of *Scn1a*-GFP mouse at 4W were counted. L2/3, L5, L6, CA1 and DG: neocortical layer II/III, V, VI, cornu ammonis 1, dentate gyrus. +, positive; -, negative.

We further performed immunohistochemical staining of PV and SST in neocortex and hippocampus of *Scn1a*-GFP mice at 4W (Figure 8). PV and SST do not co-express in cells and do not overlap. PV-INs and SST-INs were both GFP-positive, and especially GFP signals in PV-INs were intense (Figure 8A). Cell counting revealed that 21% (L2/3), 37% (L5), 37% (L6), 58% (CA1), 42% (CA2/3), and 41% (DG) of GFP-positive cells were PV or SST-positive depending on regions in neocortex and hippocampus (Figure 8B and Supplementary tables S11-S13). All PV-INs were GFP-positive (Figure 8B - middle), and most of SST-INs were GFP-positive (Figure 8B - right). Comparison of these results with those of *Vgat*-Cre mouse (Figure 7) suggests that GFP-positive GABAergic neurons in neocortex are mostly PV- or SST-positive, while in hippocampus a half of those are PV/SST-negative GABAergic neurons. Higher ratios of PV- or SST-positive cells (Figure 8B) compared with those of *Vgat*-Cre-positive cells (Figure 7C) among GFP-positive cells would be explained by that we counted PV-positive cells even if their PV-immunosignals are moderate and a significant subpopulation of such cells are known to be excitatory neurons (Jinno et al., 2004; Tanahira et al., 2009; Matho et al., 2021). Quantitative analysis of GFP signal intensity and area size of cells revealed that GFP signal intensities in PV-positive cells were significantly higher than those in PV-negative cells and GFP signal intensities in SST-positive cells were lower than those in PV-positive cells but similar to PV/SST-double negative cells (Figure 9 and Supplementary tables S14-S16). These results indicate that Nav1.1 expression level in PV-INs is significantly higher than those in excitatory neurons and PV-negative GABAergic neurons including SST-INs.

**Figure 8.**
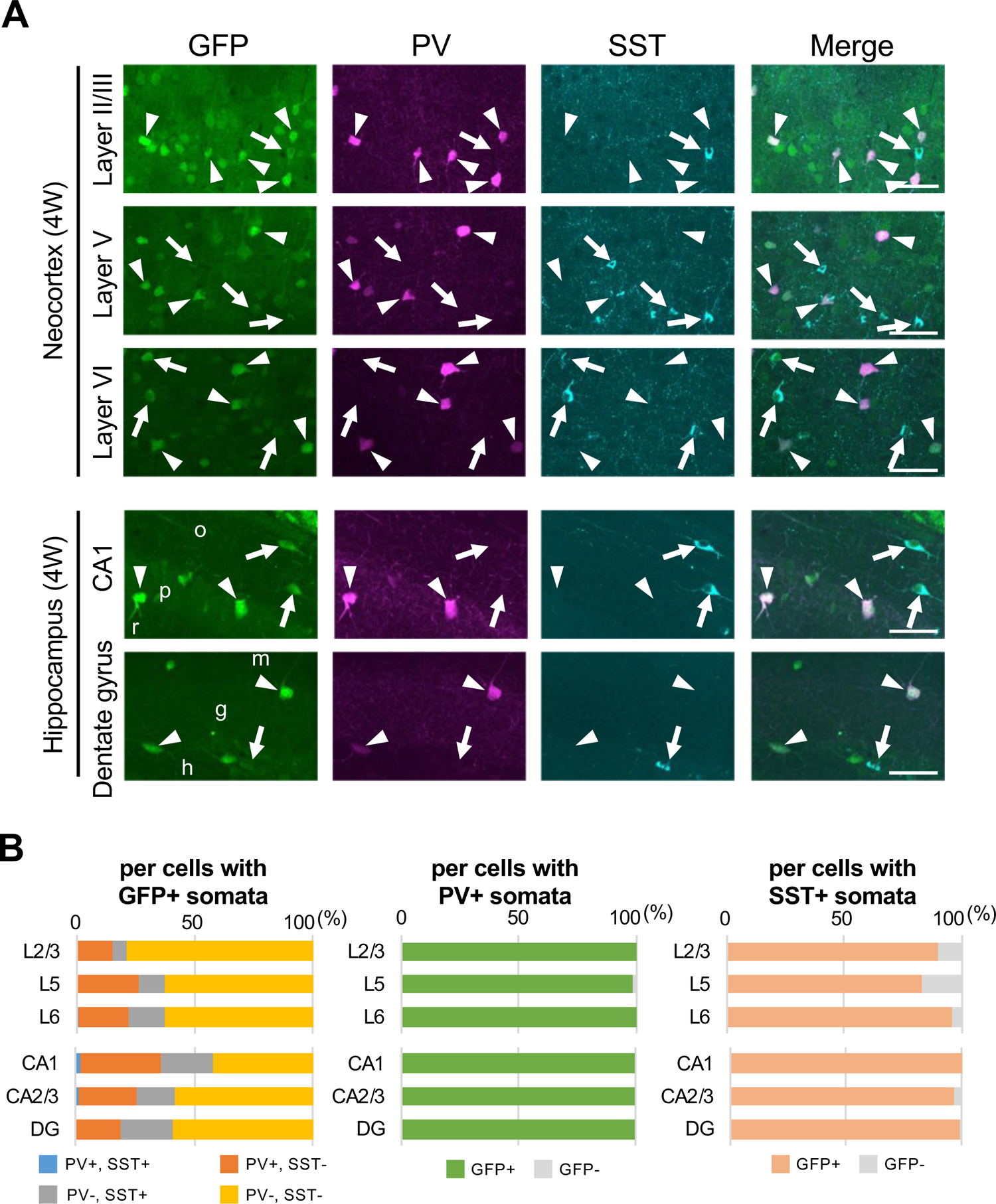
Parvalbumin or somatostatin-positive inhibitory neurons are GFP-positive in *Scn1a*-GFP mouse neocortex and hippocampus. (**A**) Triple immunofluorescent staining of parasagittal sections from 4W *Scn1a*-GFP mouse (line #233) by mouse anti-GFP (green), rabbit anti-parvalbumin (PV) (magenta) and goat anti-somatostatin (SST) (Cyan) antibodies. Regions at neocortex and hippocampus are shown. Merged images were shown in the right columns. Arrows indicate SST/GFP-double positive cells. Arrowheads indicate PV/GFP-double positive. o, stratum oriens; p, stratum pyramidale; r, stratum radiatum; h, hilus; g, stratum granulosum; m, stratum moleculare. All images are oriented from pial surface (top) to callosal (bottom). Scale bars; 50 μm. (**B**) Bar graphs indicate the percentage of cells with PV- and SST-positive/negative somata per cells with GFP-positive somata (left panel) (see also Supplementary table S11), the percentage of cells with GFP-positive/negative somata per cells with PV-positive somata (middle panel) (see also Supplementary table S12), and the percentage of cells with GFP-positive/negative somata per cells with SST-positive somata (right panel) (see also Supplementary table S13) in L2/3, L5, L6, CA1, CA2/3 and DG. Cells in neocortex and hippocampus of *Scn1a*-GFP mouse at 4W were counted. L2/3, L5, L6, CA1, CA2/3 and DG: neocortical layer II/III, V, VI, cornu ammonis 1, 2 plus 3, dentate gyrus. +, positive; -, negative.

**Figure 9.**
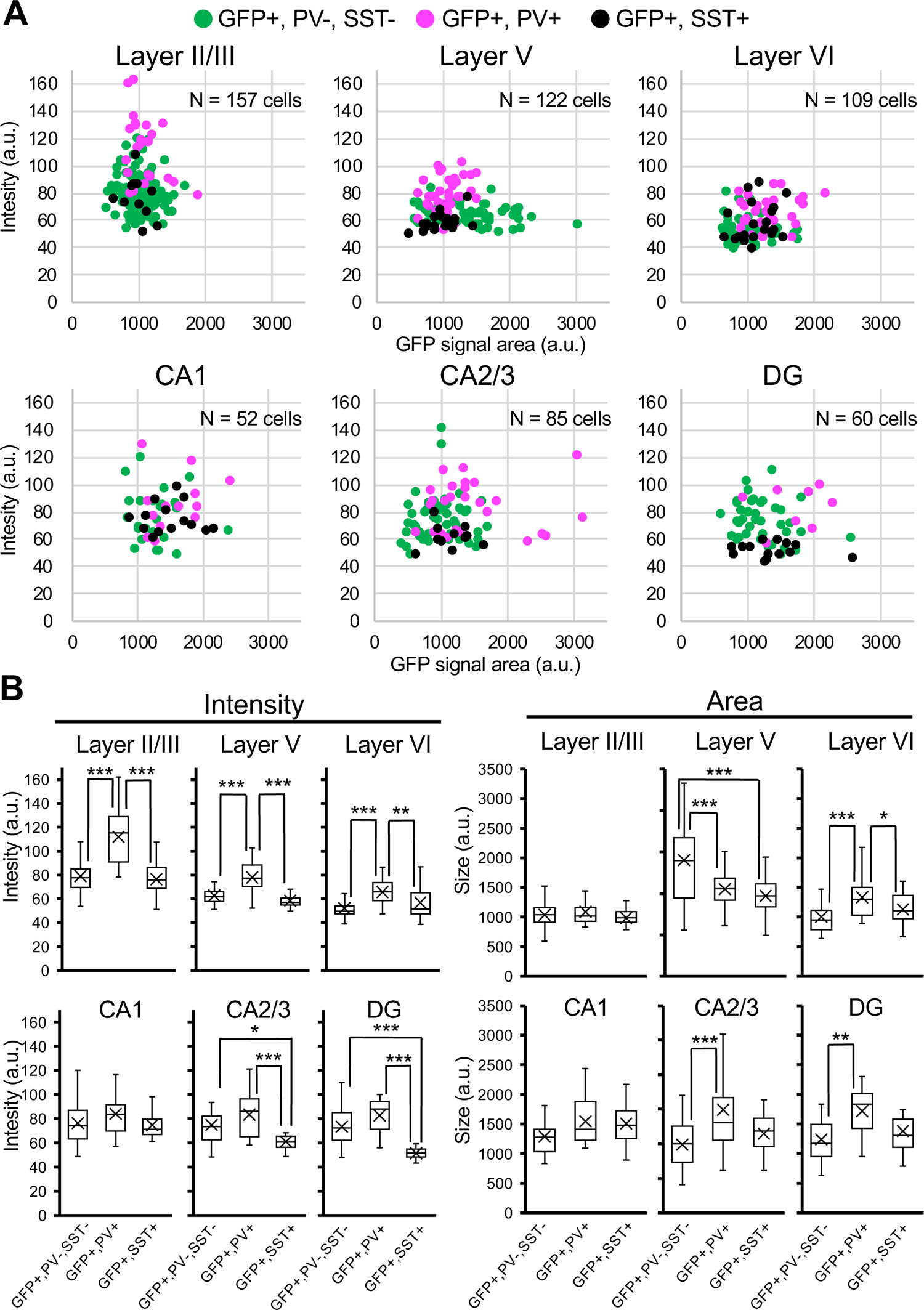
GFP-signals in parvalbumin-positive inhibitory neurons are higher than PV-negative/GFP-positive cells in *Scn1a*-GFP mouse neocortex. (**A**) Scatter plots of intensities and area sizes of GFP immunosignals in GFP-positive cells with PV or SST-positive or negative somata. Cells at primary motor cortex (upper panels) and hippocampus (lower panels) in parasagittal sections from 4W *Scn1a*-GFP mouse (line #233) were analyzed. PV-positive (magenta circles) or SST-positive (black circles) and negative (green circles) cells in neocortical L2/3, L5, and L6 or hippocampal CA1, CA2/3, and DG are plotted (see also Supplementary table S14). (**B**) Box-plots represent values for the intensity and area size in each cell type (see also Supplementary tables S15, S16). Cross marks indicate mean values in each cell type. Statistical significance was assessed using one-way ANOVA followed by Tukey–Kramer post-hoc multiple comparison test. *; p < 0.05, **; p < 0.01, ***; p < 0.001. Note that GFP signal intensities of PV/GFP-double positive cells were significantly higher than that of SST/GFP-double positive cells and PV/SST-negative/GFP-positive cells (all layers), while GFP signal intensities of SST/GFP-double positive cells were similar to PV/SST-negative/GFP-positive cells in neocortex. In hippocampus, GFP signal intensities of SST/GFP-double positive cells were significantly lower than that of SST-negative/GFP-positive cells at CA2/3 and DG. CA1, CA2/3 and DG: cornu ammonis 1, 2 plus 3, dentate gyrus. a.u., arbitrary unit; +, positive; -, negative.

### Nav1.1 and Nav1.2 expressions are mutually exclusive in mouse brain

We previously reported that expressions of Nav1.1 and Nav1.2 seem to be mutually-exclusive in multiple brain regions including neocortex, hippocampal CA1, dentate gyrus, striatum, globus pallidus, and cerebellum in wild-type mice (Yamagata et al., 2017). To further confirm it, here we performed triple immunostaining for Nav1.1, Nav1.2 and ankyrinG, and counted Nav1.1- or Nav1.2-immunopositive AISs in the neocortex of *Scn1a*-GFP mice (Figure 10). The staining again showed that Nav1.1 and Nav1.2 expressions are mutually exclusive in brain regions including neocortex (Figure 10A). The counting revealed that 5% (L2/3), 6% (L5) and 3% (L6) of AISs at P15 were Nav1.1-positive and 78% (L2/3), 69% (L5) and 69% (L6) of AISs at P15 were Nav1.2-positive in the neocortex (Figure 10B and Supplementary table S17). Of note, less than 0.5% of AISs are Nav1.1/Nav1.2-double positive confirming that Nav1.1 and Nav1.2 do not co-express. These results are consistent with our previous study (Yamagata et al., 2017) and further support that expressions of Nav1.1 and Nav1.2 are mutually exclusive in mouse neocortex at least at immunohistochemical level.

**Figure 10.**
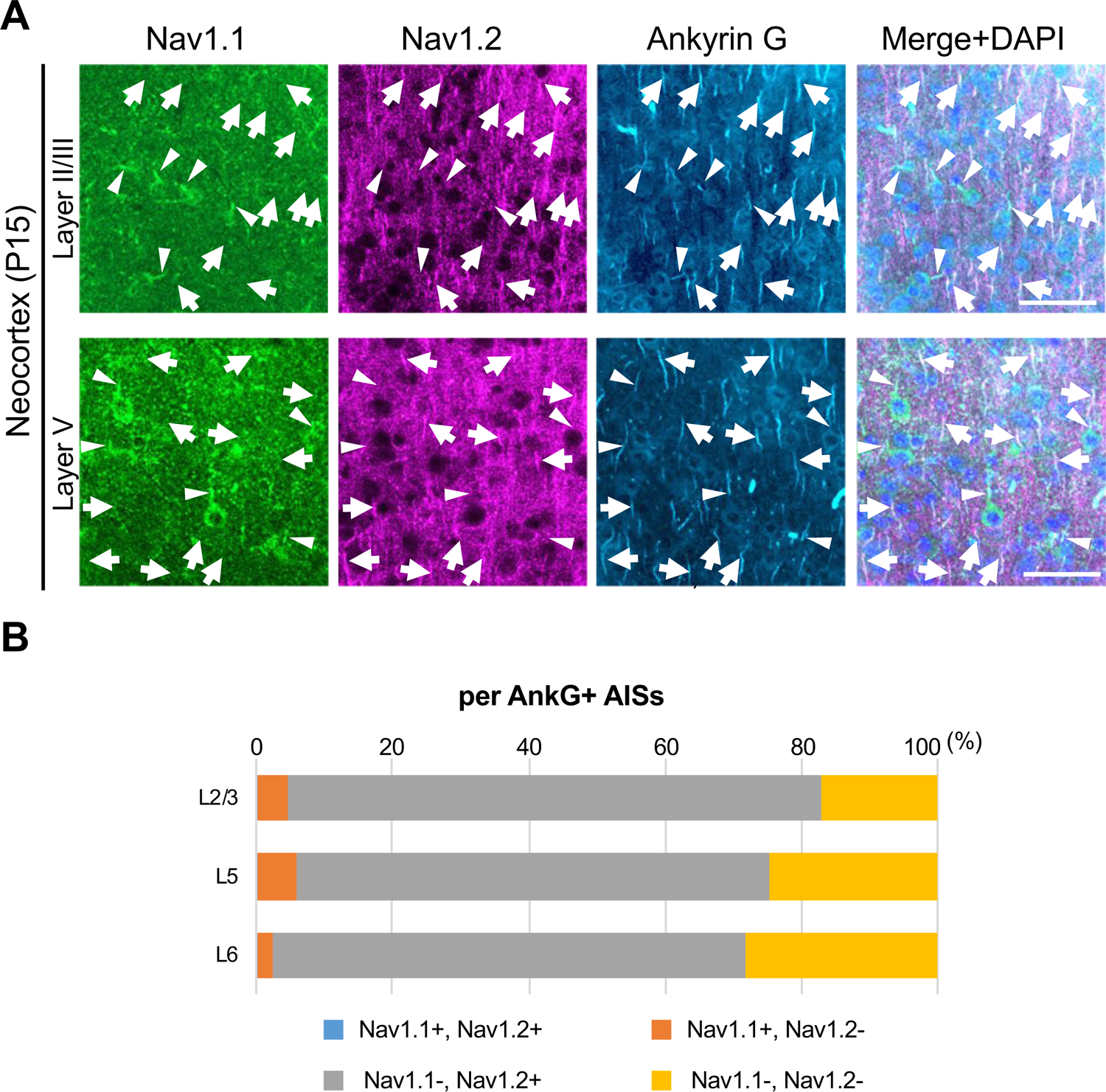
Nav1.1 and Nav1.2 are mutually exclusive at AISs in mouse brain. (**A**) Triple immunofluorescent staining on parasagittal sections from *Scn1a*-GFP mouse at P15 by rabbit anti-Nav1.1 (green), goat anti-Nav1.2 (magenta), and mouse anti-ankyrinG (cyan) antibodies. Merged images are shown in the right panels. Arrows indicate Nav1.2-positive AISs. Arrowheads indicate Nav1.1-positive AISs. Note that there are no Nav1.1/Nav1.2-double positive AISs. Images are oriented from pial surface (top) to callosal (bottom). Scale bars; 50 μm. (**B**) Bar graphs indicating the percentage of Nav1.1 and Nav1.2-positive/negative AISs per AISs detected by ankyrinG-staining in L2/3, L5, and L6 of *Scn1a*-GFP mice. Note that Nav1.1/Nav1.2-double positive AISs were less than 0.5% of all AISs in these layers (see Supplementary table S17). L2/3, L5, L6: neocortical layer II/III, V, VI. AnkG, ankyrinG; +, positive; -, negative.

As mentioned, GFP signals in *Scn1a*-GFP mouse can represent even moderate or low Nav1.1 expressions which cannot be detected by immunohistochemical staining, so some of GFP-positive cells may still express Nav1.2. To investigate whether and if so how much of GFP-positive cells have Nav1.2-positive AISs in *Scn1a*-GFP mouse neocortex, we performed triple immunohistochemical staining for Nav1.2, GFP and ankyrinG (Figure 11 and Supplementary tables S18-S20). The staining showed that AISs of GFP-positive cells are largely negative for Nav1.2, and cells with Nav1.2-positive AISs are mostly GFP-negative (Figure 11A). Cell counting revealed that 88% (L2/3), 90% (L5) and 95% (L6) of cells with Nav1.2-positive AISs at P15 were GFP-negative (Figure 11B-middle panel and Supplementary table S19), and 69% (L2/3), 83% (L5), and 86% (L6) of AISs of GFP-positive cells at P15 were Nav1.2-negative (Figure 11B-right panel and Supplementary table S20). These results indicate that the co-expression of GFP and Nav1.2 would be minimal if any.

### Neocortical pyramidal tract and a subpopulation of cortico-cortical projection neurons express Nav1.1

Neocortical excitatory neurons can be divided into functionally distinct subpopulations, a majority of those are pyramidal cells which have axons of long-range projections such as L2/3 CC, L5 PT, L5/6 CS and L6 CT projection neurons (Shepherd, 2013). Although PT neurons also project their axon collaterals to ipsilateral striatum, CS neurons project bilaterally to ipsi- and contralateral striata. A transcription factor FEZF2 is expressed in L5 PT neurons, forms their axonal projections and defines their targets (Inoue et al., 2004; Chen et al., 2005; Chen et al., 2008; Lodato et al., 2014). Most of PT neurons are FEZF2-positive (Molyneaux et al., 2005; Matho et al., 2021). We previously reported that a subpopulation of neocortical L5 pyramidal neurons is Nav1.1-positive (Ogiwara et al., 2013), but their natures were unclear. To investigate those, here we performed immunohistochemical staining of FEZF2 and GFP on *Scn1a*-GFP mouse brains (Figures 12 and 13, Supplementary tables S21-S23). In L5 where a major population of FEZF2-positive cells locate (Figure 12A), a majority of FEZF2-positive neurons were GFP-positive (83% and 96% of FEZF2-positive neurons were GFP-positive at P15 and 4W, respectively) (Figure 12B - middle panels, Supplementary table S22). In L2/3, FEZF2-positive cells were scarce (Figure 12A). In L6, a certain number of FEZF2-posiive cells exist but overlaps of FEZF2 and GFP signals are much less compared to those in L5 (Figure 12 and Supplementary tables S21-S23). Quantitative analyses revealed that FEZF2/GFP-double positive cells in L5 showed significantly lower GFP immunosignal intensities and larger signal areas (soma sizes) compared to those of FEZF2-negative/GFP-positive cells (Figure 13 and Supplementary table S24), indicating that L5 PT neurons showed lower Nav1.1 expression compared to other neurons such as PV-Ins (see also Figure 9). Soma sizes of GFP-positive cells in L6 were overall smaller than those of FEZF2/GFP-double positive cells in L5, and there was no statistically significant difference in size between the FEZF2-positive/negative subpopulations. However, FEZF2/GFP-double positive cells still showed lower intensity of GFP signals compared to FEZF2-negative/GFP-positive cells (Figure 13 and Supplementary table S24).

**Figure 11.**
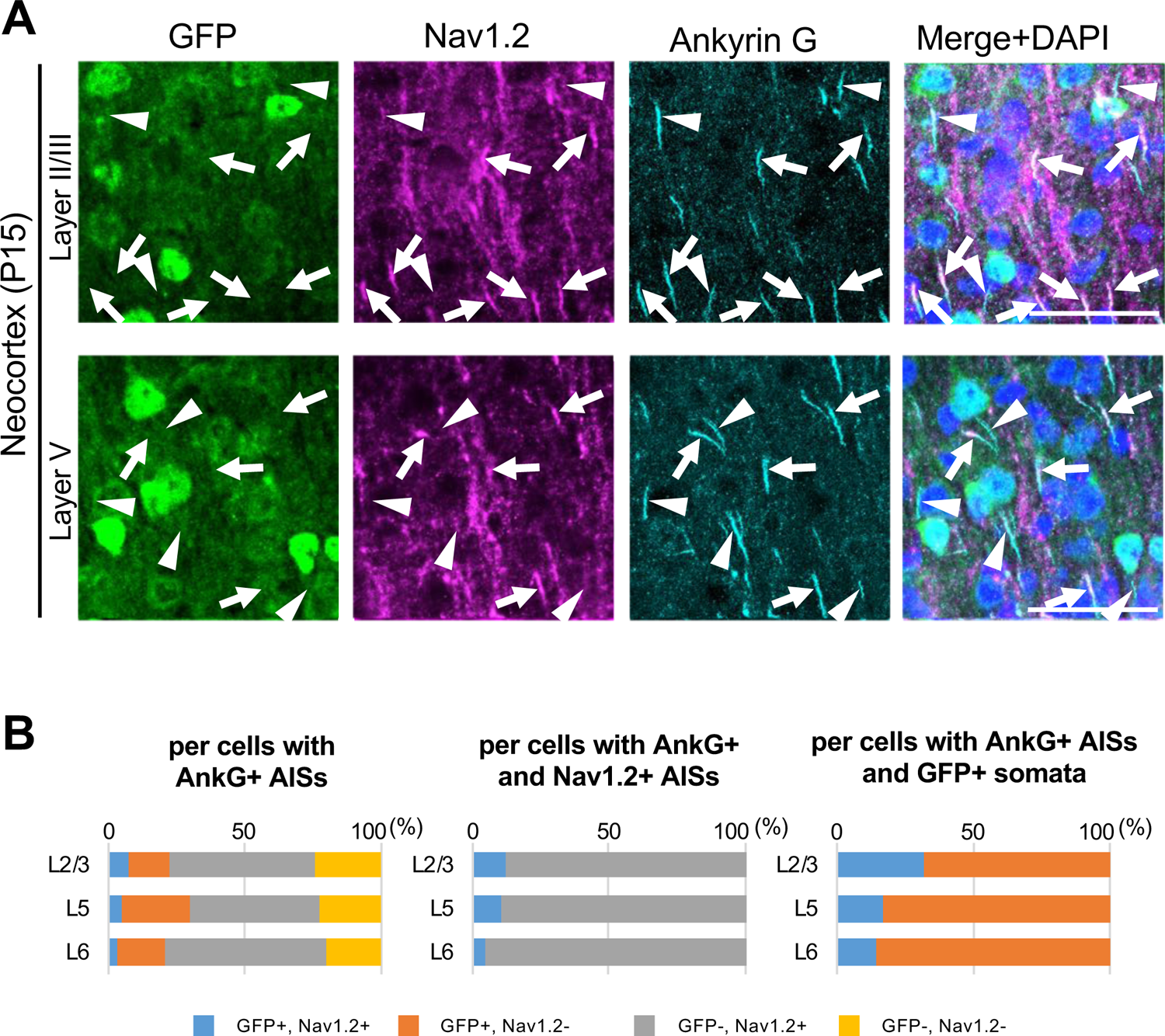
Cells with Nav1.2-positive AISs are mostly GFP-negative in *Scn1a*-GFP mouse neocortex. (**A**) Triple immunofluorescent staining of parasagittal sections from P15 *Scn1a*-GFP mouse brain (line #233) by mouse anti-GFP (green), goat anti-Nav1.2 (magenta) and rabbit anti-ankyrinG (cyan) antibodies. Merged images of the signals are shown in the right panels. Arrows indicate Nav1.2-positive AISs of cells with GFP-negative somata. Arrowheads indicate Nav1.2-negative AISs of cells with GFP-positive somata. All images are oriented from pial surface (top) to callosal (bottom). Scale bars; 50μm. (**B**) Bar graphs indicating the percentage of cells with GFP- and Nav1.2-positive/negative somata and AISs per cells with ankyrinG-positive AISs (left panel) (see also Supplementary table S18), the percentage of cells with GFP-positive/negative somata per cells with ankyrinG/Nav1.2-double positive AISs (middle panel) (see also Supplementary table S19), and the percentage of cells with Nav1.2-positive/negative AISs per cells with ankyrinG-positive AISs and GFP-positive somata (right panel) (see also Supplementary table S20) in L2/3, L5, and L6. Note that 88% (L2/3), 90% (L5) and 95% (L6) of cells with Nav1.2-positive AISs have GFP-negative somata (middle panel), and 68% (L2/3), 83% (L5) and 86% (L6) of cells with GFP-positive somata have Nav1.2-negative AISs (right panel). L2/3, L5, L6: neocortical layer II/III, V, VI. AnkG, ankyrinG; +, positive; -, negative.

**Figure 12.**
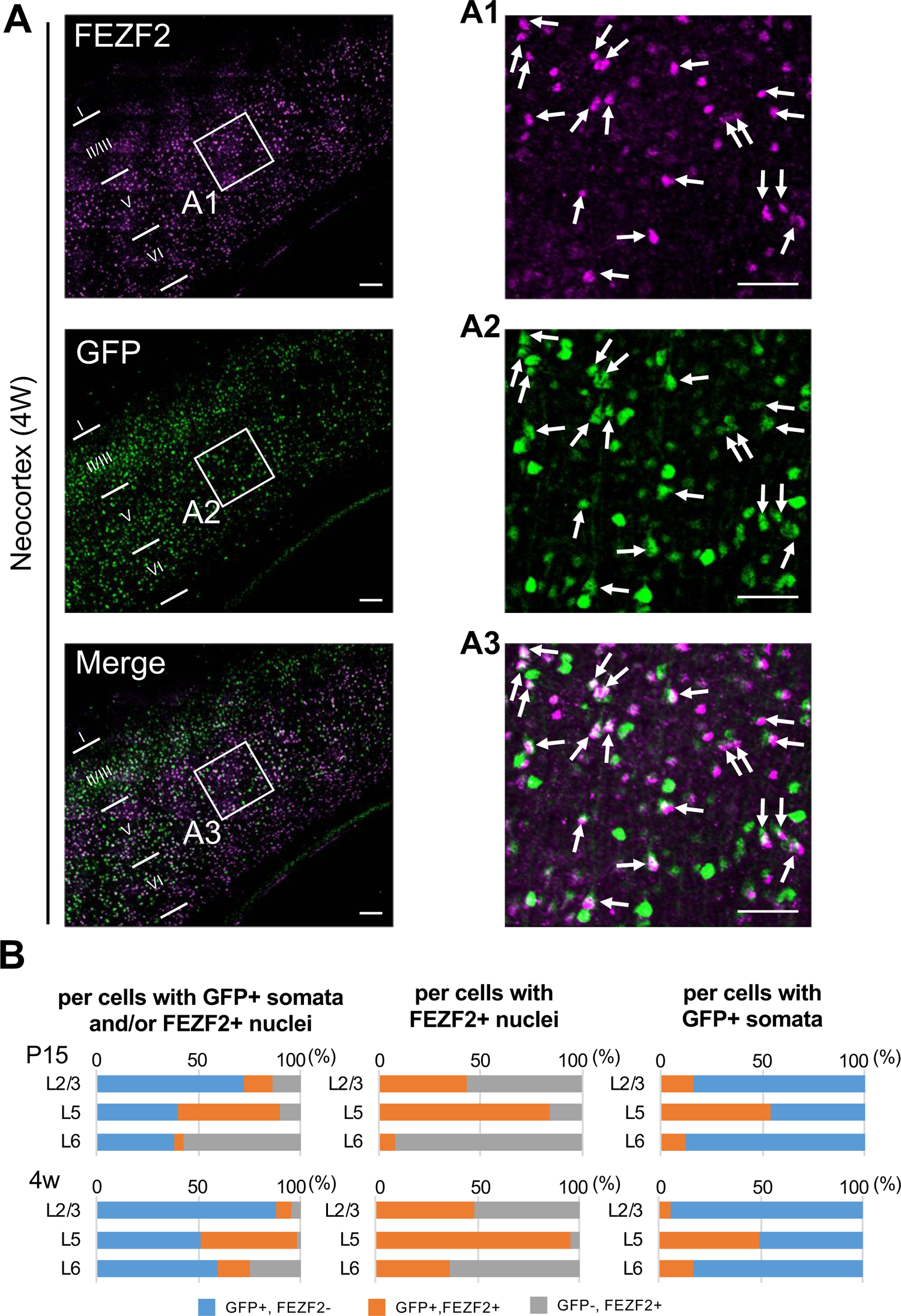
Cells positive for FEZF2 are mostly GFP-positive at L5 of *Scn1a*-GFP mouse neocortex. (**A**) Double immunostaining of FEZF2 and GFP in neocortex of 4W *Scn1a*-GFP mouse (line #233) by rabbit anti-FEZF2 (magenta) and mouse anti-GFP (green) antibodies. Arrows indicate FEZF2/GFP-double positive cells. Magnified images outlined in (A) are shown in (A1-A3). Note that FEZF2 signals mostly overlap with GFP signals in L5. Many of the remained GFP-positive/FEZF2-negative cells have intense GFP signals and are assumed to be inhibitory neurons (see Figure 8). Scale bars; 100 μm (A), 50 μm (A1-A3). (**B**) Bar graphs indicating the percentage of cells with FEZF2- and GFP-positive/negative nuclei and somata per cells with GFP-positive somata and/or FEZF2-positive nuclei (left panels) (see also Supplementary table S21), the percentage of cells with GFP-positive/negative somata per cells with FEZF2-positive nuclei (middle panels) (see also Supplementary table S22), and the percentage of cells with FEZF2-positive/negative nuclei per cells with GFP-positive somata (right panels) (see also Supplementary table S23) in L2/3, L5, and L6. Cells at primary motor cortex of *Scn1a*-GFP mouse at P15 and 4W were counted. Note that 83% (P15) and 96% (4W) of cells with FEZF2-positive cells are GFP-positive in L5 (middle panels), but a half of cells with GFP-positive cells are FEZF2-positive in L5 (right panel). L2/3, L5, L6: neocortical layer II/III, V, VI. +, positive; -, negative.

**Figure 13.**
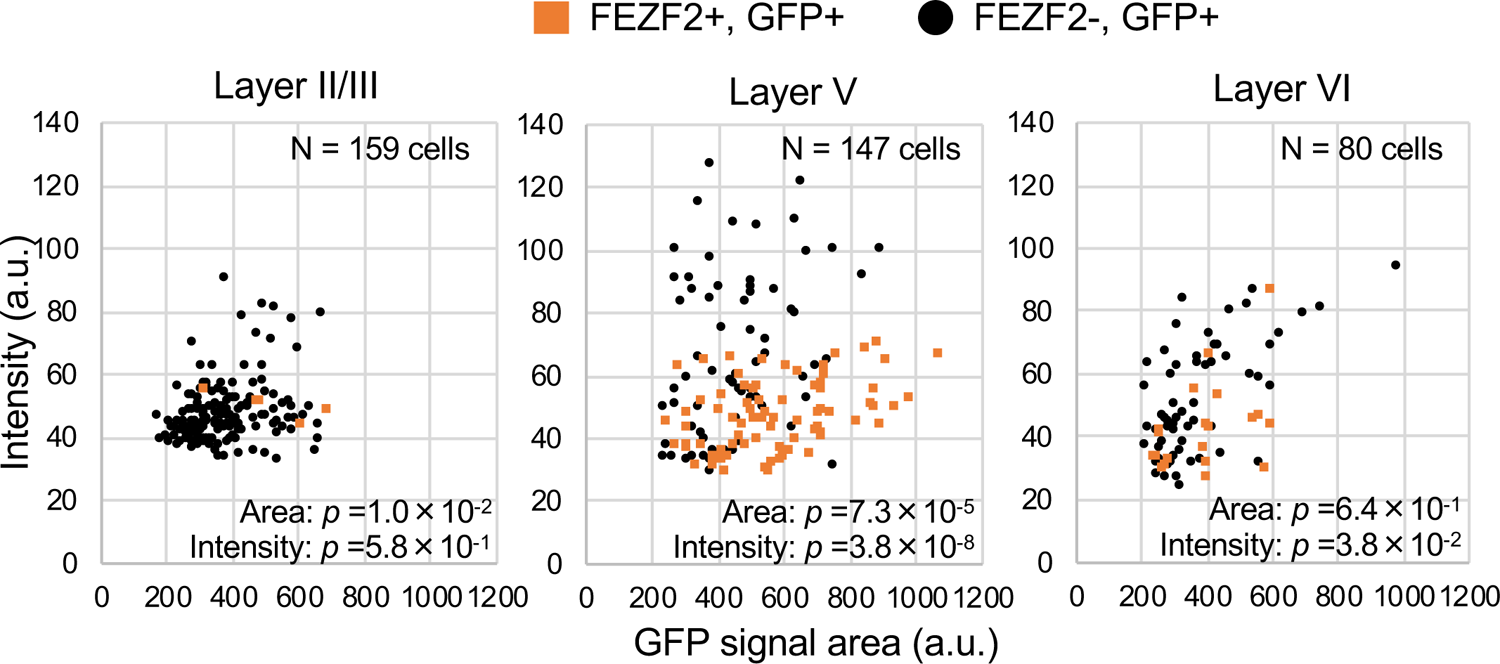
FEZF2-positive cells have lower GFP signal intensities in *Scn1a*-GFP mouse neocortex. Scatter plots of intensities and area sizes of GFP immunosignals in GFP-positive cells with FEZF2-positive/negative nuclei. Cells at primary motor cortex in parasagittal sections from 4W *Scn1a*-GFP mouse (line #233) were analyzed. FEZF2-positive (orange squares) and negative (black circles) cells in neocortical L2/3, L5, and L6 are plotted (see also Supplementary table S24). Note that GFP signal intensities of FEZF2/GFP-double positive cells were significantly lower than that of FEZF2-negative/GFP-positive cells (L5, L6), and signal area size of FEZF2/GFP-double positive cells were significantly larger than that of FEZF2-negative/GFP-positive cells (L2/3, L5). Statistical significance was assessed using t-test. a.u., arbitrary unit; +, positive; -, negative.

We further performed triple immunostaining of FEZF2, Nav1.1 and ankyrinG on *Scn1a*-GFP mice at P15, and found that 11% of FEZF2-positive cells have Nav1.1-positive AIS in L5 of *Scn1a*-GFP mouse neocortex (Supplementary figure S8 and Supplementary tables S25-S27). The low ratios of FEZF2/Nav1.1-double positive cells are most possibly due to immunohistochemically-undetectable low levels of Nav1.1 expression in these excitatory neurons.

We also performed triple immunostaining of FEZF2, Nav1.2 and ankyrinG on *Scn1a*-GFP mice at P15 (Supplementary figure S9 and Supplementary tables S28-S31). The staining showed that 20% of neurons (cells with ankyrinG-positive AISs) in L5 are FEZF2-positive and a half of L5 FEZF2-positive cells have Nav1.2-positive AISs. Together with the observation that most of FEZF2-positive cells are GFP-positive (Figure 12), these results indicate that a subpopulation of FEZF2-positive PT neurons may express both Nav1.1 and Nav1.2.

We additionally performed triple immunostaining of FEZF2, GFP, and Nav1.2 on *Scn1a*-GFP mice at P15 (Supplementary figure S10 and Supplementary table S32), showing that in L5 74% of FEZF2-positive cells are GFP-positive but a majority of their AISs are Nav1.2-negative. The ratios of Nav1.2-positive cells among FEZF2-positive cells obtained in the triple-immunostaining of FEZF2, GFP and Nav1.2 (Supplementary figure S10) are 27% (L5) and 40% (L6). These results further support the above notion that a subpopulation of FEZF2-positive PT neurons may express both Nav1.1 and Nav1.2. Although further studies such retrograde tracking analyses are required to confirm and figure out the detailed circuits, all these results propose that the majority of L5 PT neurons express Nav1.1.

### The majority of cortico-thalamic, cortico-striatal and cortico-cortical projection neurons express Nav1.2

TBR1 (T-box brain 1 transcription factor) is a negative regulator of FEZF2 and therefore not expressed in the PT neurons, all of which are known to be FEZF2-positive (Chen et al., 2008; Han et al., 2011; McKenna et al., 2011). TBR1 is predominantly expressed in L6 CT neurons and subpopulations of L2/3 and L5 non-PT excitatory neurons instead (Han et al., 2011; McKenna et al., 2011; Matho et al., 2021). To further elucidate the distributions of GFP (Nav1.1)-expressing neurons in neocortex, we performed immunohistochemical staining of TBR1 on *Scn1a*-GFP mice (Figure 14 and Supplementary tables S33-S35) and quantitated GFP signal intensities and area sizes of cells (Figure 15 and Supplementary table S36). TBR1-positive cells were predominant in neocortical L6, some in L5 and a few in L2/3. In L5, contrary to the high ratios of GFP-positive cells among FEZF2-positive cells (83% at P15, and 96% at 4W) (Figure 12B), the ratios of GFP-positive cells among TBR1-positive cells are quite low (11% at P15, and 5% at 4W) (Figure 14B-middle panels and Supplementary table S34). Soma sizes of TBR1/GFP-double positive cells were smaller than those of TBR1-negative/GFP-positive cells (Figure 15-middle panel and Supplementary table S36). In L6 where a major population of TBR1-positive neurons locate, the ratios of GFP-positive cells among TBR1-positive cells are still low (15% at P15, and 26% at 4W) (Figure 14B-middle panels and Supplementary table S34). Soma sizes of TBR1/GFP-double positive cells were also smaller than those of TBR1-negative/GFP-positive cells, and TBR1/GFP-double positive cells showed lower intensity of GFP immunosignals compared to TBR1-negative/GFP-positive cells (Figure 15-right panel and Supplementary table S36). We additionally performed triple immunostaining for TBR1, Nav1.1, and ankyrinG on *Scn1a*-GFP mice (Supplementary figure S11 and Supplementary tables S37-S39). Notably, the ratios of Nav1.1-positive cells among TBR1-positive cells are 0% in all layers (Supplementary figure S11B-right panel and Supplementary table S39). These results indicate that the major population of TBR1-positive cells do not express Nav1.1.

**Figure 14.**
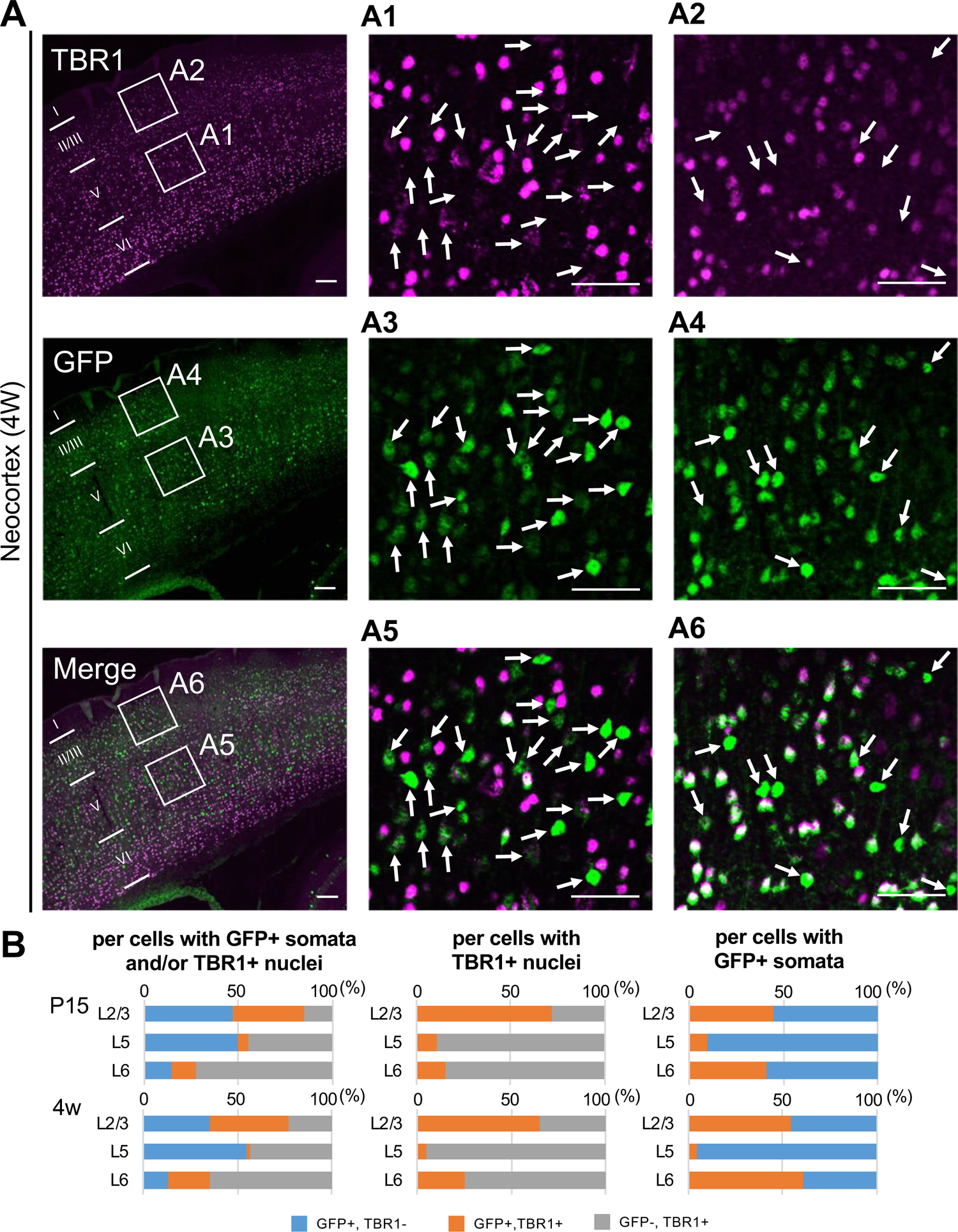
GFP-positive cells were mostly negative for TBR1 at L5 of *Scn1a*-GFP mouse neocortex. (**A**) Double immunostaining of TBR1 and GFP in neocortex of 4W *Scn1a*-GFP mouse (line #233) detected by mouse rabbit anti-TBR1 (magenta) and anti-GFP (green) antibodies. Arrows indicate TBR1-negative/GFP-positive cells. Magnified images outlined in (A) are shown in (A1-A6). Note that at L5 GFP-positive cells were mostly TBR1-negative but at L2/3 more than half of GFP-positive cells were TBR1-positive. Scale bars; 100 μm (A), 50 μm (A1-A6). (**B**) Bar graphs indicating the percentage of cells with TBR1- and GFP-positive/negative nuclei and somata per GFP-positive cells and/or TBR1-positive nuclei (left panels) (see also Supplementary table S33), the percentage of cells with GFP-positive/negative somata per cells with TBR1-positive nuclei (middle panels) (see also Supplementary table S34), and the percentage of cells with TBR1-positive/negative nuclei per cells with GFP-positive somata (right panels) (see also Supplementary table S35) in L2/3, L5, and L6. Cells in primary motor cortex of *Scn1a*-GFP mouse at P15 and 4W were counted. Note that 86% (P15) and 95% (4W) of cells with TBR1-positive cells are GFP-negative in L5 (middle panels), and 90% (P15) and 96% (4W) of cells with GFP-positive cells are TBR1-negative in L5 (right panel). L2/3, L5, L6: neocortical layer II/III, V, VI. +, positive; -, negative.

**Figure 15.**
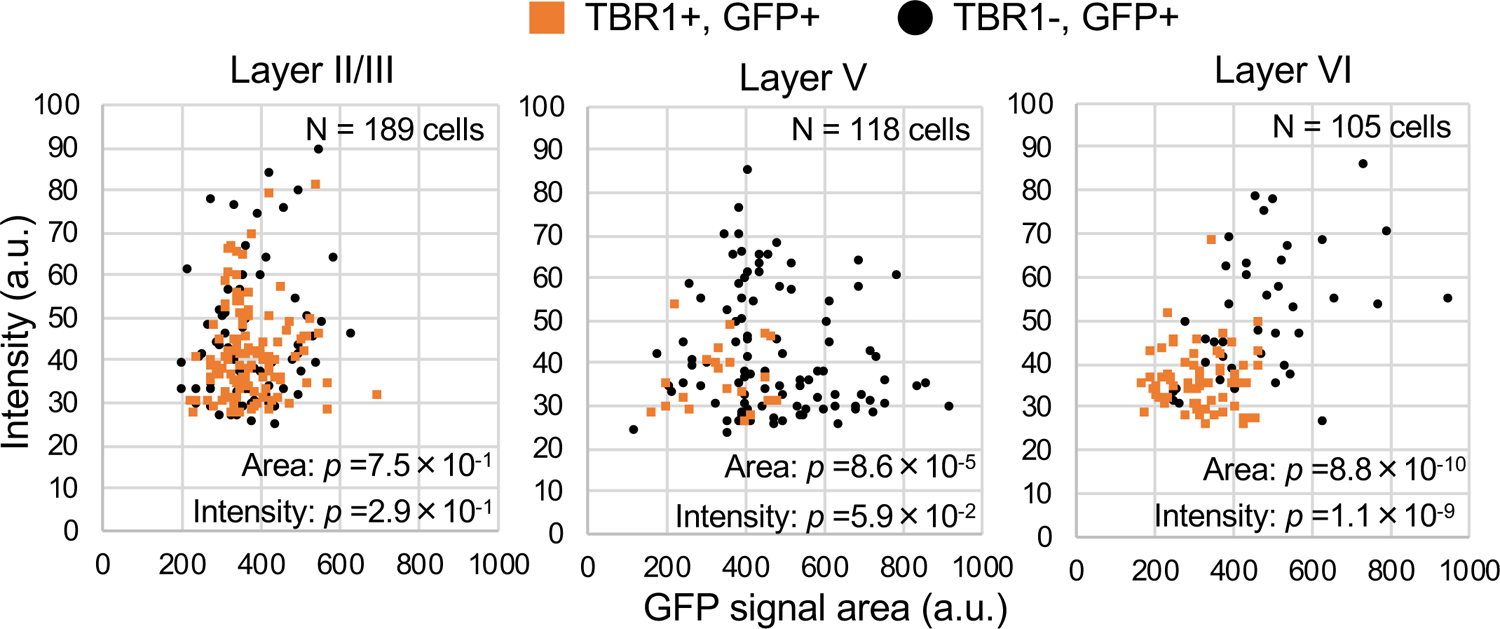
TBR1-positive cells have lower GFP signal intensities in *Scn1a*-GFP mouse neocortex. Scatter plots of intensities and area sizes of GFP immunosignals in GFP-positive cells with TBR1-positive/negative nuclei. Cells at primary motor cortex in parasagittal sections from 4W *Scn1a*-GFP mouse (line #233) were analyzed. TBR1-positive (orange squares) and negative (black circles) cells in neocortical L2/3, L5, and L6 are plotted (see also Supplementary table S36). Note that GFP signal intensities of TBR1/GFP-double positive cells were significantly lower than that of TBR1-negative/GFP-positive cells (L6), and signal area size of TBR1/GFP-double positive cells was significantly smaller than that of TBR1-negative/GFP-positive cells (L5, L6). Statistical significance was assessed using t-test. a.u., arbitrary unit; +, positive; -, negative.

To investigate whether TBR1-positive cells express Nav1.2, we performed triple immunostaining of TBR1, Nav1.2 and ankyrinG on *Scn1a*-GFP mice (Supplementary figure S12 and Supplementary tables S40-S43). In L5, contrary to the low ratios of GFP-positive cells among TBR1-positive cells (11% at P15 and 5% at 4W) (Figure 14B-middle panels and Supplementary table S34), the ratios of Nav1.2-positive cells among TBR1-positeve cells are high (69% (L2/3), 69%(L5) and 69% (L6) at P15) (Supplementary figure S12B-right-upper panel and Supplementary table S42). The ratios of TBR1-positive cells among Nav1.2-positive cells are 29% (L2/3), 53% (L5) and 62% (L6) at P15 (Supplementary figure S12B-middle upper panel and Supplementary table S41).

We further performed triple immunostaining for TBR1, Nav1.2 and GFP on *Scn1a*-GFP mice (Supplementary figure S13 and Supplementary table S44) and found that most (88%) of L6 TBR1-positive cells are GFP-negative. Taken together, these results indicate that most TBR1-positive neurons including L6 CT neurons do not express Nav1.1 but expresses Nav1.2.

As a whole, above results showed that a minor subpopulation of L2/3 CC and L5 PT neurons express Nav1.1 while the majority of L2/3 CC, L5/6 CS and L6 CT neurons express Nav1.2. A breakdown of L5 neuron species containing PT neurons is specifically described (Supplementary figure S14).

## Discussion

In our present study, we developed *Scn1a* promoter-driven GFP mice in which the expression of GFP mimics that of Nav1.1. All PV-INs and most of SST-INs were GFP-positive in the neocortex and hippocampus of the *Scn1a*-GFP mouse, being consistent with the previous reports of Nav1.1 expression in those inhibitory neurons (Ogiwara et al., 2007; Lorincz and Nusser, 2008; Ogiwara et al., 2013; Li et al., 2014; Tai et al., 2014; Tian et al., 2014; Yamagata et al, 2017). All Nav1.1-positive cells were GFP-positive. Reversely all GFP-positive cells were also Nav1.1-positive in the hippocampus, but in the neocortex only a half of GFP-positive cells were Nav1.1-positive. This is largely because in the hippocampus Nav1.1 expression is restricted to inhibitory neurons, but in the neocortex Nav1.1 is expressed not only in inhibitory but also in a subpopulation of excitatory neurons in which Nav1.1 expression is low and not easily detected immunohistochemically by anti-Nav1.1 antibodies. In neocortex, one third of GFP-positive cells were GABAergic cells such as PV-INs and SST-INs, and remained two-third were excitatory neurons. GFP signals were especially intense in PV-positive cells indicating strong Nav1.1 expression in those cells, while GFP-signals in SST-positive cells were similar to those in excitatory neurons. In addition, extensive immunostaining analyses using projection neuron markers FEZF2 and TBR1 together with anti-Nav1.2 antibody also revealed that most L5 PT neurons and a minor subpopulation of L2/3 CC neurons express Nav1.1 while the majority of L6 CT, L5/6 CS and L2/3 CC neurons express Nav1.2.

The above observations should contribute to understanding of neural circuits responsible for diseases such as epilepsy and neurodevelopmental disorders caused by *SCN1A* and *SCN2A* mutations. Dravet syndrome is a sporadic intractable epileptic encephalopathy characterized by early onset (6 months ∼ 1 year after birth) epileptic seizures, intellectual disability, autistic features, ataxia and increased risk of sudden unexpected death in epilepsy (SUDEP). De novo loss-of-function mutations of *SCN1A* are found in more than 80% of the patients (Claes et al., 2001; Sugawara et al., 2002; Fujiwara et al., 2003; Dravet et al., 2005; Depienne et al., 2009; Meng et al., 2015). In mice, loss-of-function *Scn1a* mutations caused clinical features reminiscent of Dravet syndrome, including early-onset epileptic seizures, hyperactivity, learning and memory deficits, reduced sociability and ataxic gaits and premature sudden death (Yu et al., 2006; Ogiwara et al., 2007; Oakley et al., 2009; Cao et al., 2012; Han et al., 2012; Kalume et al., 2013; Ito et al., 2013). As also shown in the present study, Nav1.1 is densely localized at AISs of inhibitory cells such PV-IN (Ogiwara et al., 2007; Ogiwara et al., 2013; Li et al., 2014; Tai et al., 2014) and selective elimination of Nav1.1 in PV-IN in mice leads to epileptic seizures, sudden death and deficits in social behavior and spatial memory (Ogiwara et al., 2013; Tatsukawa et al., 2018). It is thus plausible that Nav1.1 haplo-deficiency in PV-IN plays a pivotal role in the pathophysiology of many clinical aspects of Dravet syndrome. Notably, mice with selective Nav1.1 haplo-elimination in global inhibitory neurons were at a greater risk of lethal seizure than systemic Nav1.1 haplo-deficient mice, and the mortality risk of mice with Nav1.1 haplo-deficiency in inhibitory neurons was significantly decreased or improved with additional Nav1.1 haplo-elimination in dorsal telencephalic excitatory neurons (Ogiwara et al., 2013), which indicates beneficial effects of Nav1.1 deficiency in excitatory neurons for epileptic seizures and sudden death. Because of the absence of Nav1.1 in hippocampal excitatory neurons (Ogiwara et al., 2007; Ogiwara et al., 2013; Yamagata et al., 2017, and the present study), the ameliorating effect was most possibly caused by Nav1.1 haploinsufficiency in neocortical excitatory neurons. Kalume and colleagues (2013) reported that parasympathetic hyperactivity is observed in Nav1.1 haplo-deficient mice and it causes ictal bradycardia and finally result in seizure-associated sudden death.

Our present finding of Nav1.1 expression in L5 pyramidal tract projection neurons which innervate the vagus nerve may possibly elucidate the ameliorating effects of Nav1.1 haploinsufficiency in neocortical excitatory neurons for sudden death of Nav1.1 haplo-deficient mice and may contribute to the understanding of the neural circuit for SUDEP in patients with Dravet syndrome. Further studies including retrograde tracing and electrophysiological analyses are awaited.

We previously proposed that impaired cortico-striatal excitatory neurotransmission causes epilepsies in *Scn2a* haplodeficient mouse (Miyamoto et al., 2019). Our present finding of Nav1.2 expression in CS neurons is consistent and further support the proposal. Because *SCN2A* has been well established as one of top genes which show de novo loss-of-function mutations in patients with ASD (Hoischen et al., 2014; Johnson et al., 2016) and because impaired striatal function was suggested in multiple ASD animal models (Fuccillo et al., 2016), our finding of Nav1.2 expression in CS neurons may also contribute to the understanding of neural circuit for ASD caused by *SCN2A* mutations.

In summary, the present investigations using a newly-developed *Scn1a* promoter-driven GFP mice together with anti-Nav1.1/Nav1.2 antibodies and neocortical neuron markers revealed the expression loci of Nav1.1 and Nav1.2 in more detail. Further developments of transgenic mice for other sodium channel genes’ promoter-driven reporter molecules and combinatorial analyses of those mice are awaited to segregate and redefine their unique functional roles in each of highly diverse neuronal species and complexed neural circuits.

## Materials and Methods

### Animal work statement

All animal experimental protocols were approved by the Animal Experiment Committee of Nagoya City University and RIKEN Center for Brain Science. Mice were handled in accordance with the guidelines of the Animal Experiment Committee.

### Mice

*Scn1a*-GFP BAC transgenic mice were generated as follows. A murine BAC clone RP23-232A20 containing the *Scn1a* locus was obtained from the BACPAC Resource Center (https://bacpacresources.org). A GFP reporter cassette, comprising a red-shifted variant GFP cDNA and a downstream polyadenylation signal derived from pIRES2-EGFP (Takara Bio), was inserted in-frame into the initiation codon of the *Scn1a* coding exon 1 using the Red/ET Recombineering kit (Gene Bridges), according to the manufacturer’s instructions. A correctly modified BAC clone verified using PCR and restriction mapping was digested with *Sac*II, purified using CL-4B sepharose (GE Healthcare), and injected into pronuclei of C57BL/6J zygotes. Mice carrying the BAC transgene were identified using PCR with primers: m*Scn1a*_TG_check_F1, 5’-TGTTCTCCACGTTTCTGGTT-3’, m*Scn1a*_TG_check_R1, 5’-TTAGCCTTCTCTTCTGCAATG-3’ and EGFP_R1, 5’-GCTCCTGGACGTAGCCTTC-3’ that detect the wild-type *Scn1a* allele as an internal control (186 bp) and the inserted transgene (371 bp). Of fifteen independent founder lines that were crossed with C57BL/6J mice, twelve lines successfully transmitted the transgene to their progeny. Of twelve founders, two lines (#184 and 233) that display much stronger green fluorescent intensity compared with other lines were selected, and maintained on a congenic C57BL/6J background. The mouse lines had normal growth and development. The line #233 has been deposited to the RIKEN BioResource Research Center (https://web.brc.riken.jp/en/) for distribution under the registration number RBRC10241. *Vgat*-Cre BAC transgenic mice and loxP flanked transcription terminator cassette CAG promotor driven tdTomato transgeni (*Rosa26*-tdTomato; B6.Cg- Gt(ROSA)26Sortm14(CAG-tdTomato)Hze/J, Stock No: 007914, The Jackson Laboratory, USA) mice were maintained on a C57BL/6J background. To generate triple mutant mice (*Scn1a*-GFP^gfp/-^, *Vgat*-Cre^cre/-^, *Rosa26*-tdTomamto^tomato/-^), heterozygous *Scn1a*-GFP and *Vgat*-Cre mice were mated with homozygous *Rosa26*-tdTomato transgenic mice.

### Western blot analysis

Mouse brains at 5W were isolated and homogenized in homogenization buffer [(0.32 M sucrose, 10 mM HEPES, 2 mM EDTA and 1× complete protease inhibitor cocktail (Roche Diagnostics), pH 7.4)], and centrifuged for 15 min at 1,000 *g*. The supernatants were next centrifuged for 30 min at 30,000 *g*. The resulting supernatants were designated as the cytosol fraction. The pellets were subsequently resuspended in lysis buffer (50 mM HEPES and 2 mM EDTA, pH 7.4) and centrifuged for 30 min at 30,000 *g*. The resulting pellets, designated as the membrane fraction, were dissolved in 2 M Urea, 1× NuPAGE reducing agent (Thermo Fisher Scientific) and 1× NuPAGE LDS sample buffer (Thermo Fisher Scientific). The cytosol and membrane fractions were separated on the NuPAGE Novex Tris-acetate 3–8% gel (Thermo Fisher Scientific) or the PAG mini SuperSep Ace Tris-glycine 5–20% gel (FUJIFILM Wako Pure Chemical), and transferred to nitrocellulose membranes (Bio-Rad). Membranes were probed with the rabbit anti-Nav1.1 (250 ng/ml; IO1, Ogiwara et al., 2007), chicken anti-GFP (1:5,000; ab13970, Abcam) and mouse anti-β tubulin (1:10,000; T0198, Sigma-Aldrich) antibodies, and incubated with the horseradish peroxidase-conjugated goat anti-rabbit IgG (1:2,000; sc-2004, Santa Cruz Biotechnology), rabbit anti-chicken IgY (1:1,000; G1351, Promega) and goat anti-mouse IgG (1:5,000; W4011, Promega) antibodies. Blots were detected using the enhanced chemiluminescence reagent (PerkinElmer). The intensity of the Nav1.1 immunosignals were quantified using the Image Studio Lite software (LI-COR, Lincoln, Nebraska USA) and normalized to the level of β tubulin or Glyceraldehyde 3-phosphate dehydrogenase.

### Histochemistry

Mice were deeply anesthetized, perfused transcardially with 4% paraformaldehyde (PFA) in PBS (10 mM phosphate buffer, 2.7 mM KCl, and 137 mM NaCl, pH 7.4) or periodate-lysine-4% PFA (PLP). Brains were removed from the skull and post-fixed. For fluorescent imaging, PFA-fixed brains were cryoprotected with 30% sucrose in PBS, cut in 30 µm parasagittal sections, and mounted on glass slides. The sections on glass slides were treated with TrueBlack Lipofuscin Autofluorescence Quencher (Biotium) to reduce background fluorescence. For immunostaining, frozen parasagittal sections (30 µm) were blocked with 4% BlockAce (DS Pharma Biomedical) in PBS for 1 hour at room temperature (RT), and incubated with rat anti-GFP (1:500; GF090R, Nacalai Tesque). The sections were then incubated with the secondary antibodies conjugated with biotin (1:200; Vector Laboratories). The antibody-antigen complexes were visualized using the Vectastain Elite ABC kit (Vector Laboratories) with Metal enhanced DAB substrate (34065, PIERCE). For immunofluorescent staining, we prepared 6 µm parasagittal sections from paraffin embedded PLP-fixed brains of mice. The sections were processed as previously described (Yamagata et al., 2017). Following antibodies were used to detect GFP, Nav1.1, Nav1.2, TBR1, FEZF2, ankyrinG, NeuN, parvalbumin and somatostatin; mouse anti-GFP antibodies (1:500; 11814460001, Roche Diagnostics), anti-Nav1.1 antibodies (1:10,000; rabbit IO1, 1:500; goat SC-16031, Santa Cruz Biotechnology), anti-Nav1.2 antibodies (1:1,000; rabbit ASC-002, Alomone Labs; goat SC-31371, Santa Cruz Biotechnology), rabbit anti-TBR1 antibody (1:1,000; ab31940, Abcam or 1:500; SC-376258, Santa Cruz Biotechnology), rabbit anti-FEZF2 antibody (1:500; #18997, IBL), ankyrinG antibodies (1:500; mouse SC-12719, rabbit SC-28561; goat, SC-31778, Santa Cruz Biotechnology), mouse anti-NeuN biotin conjugated antibody (1:2,000; MAB377B, Millipore), rabbit anti-parvalbumin (1:5,000; PC255L, Merck) and rabbit anti-somatostatin (1:5,000; T-4103, Peninsula Laboratories, 1:1,000; SC-7819, Santa Cruz Biotechnology) antibodies. As secondary antibodies, Alexa Fluor Plus 488, 555, 594 and 647 conjugated antibodies (1:1,000; A32723, A32766, A32794, A32754, A32849, A32795, A32787, Thermo Fisher Scientific) were used. To detect NeuN, Alexa-647 conjugated streptavidin (1:1,000; S21374, Thermo Fisher Science) was used. Images were captured using fluorescence microscopes (BZ-8100 and BZ-X710, Keyence), and processed with Adobe Photoshop Elements 10 (Adobe Systems) and BZ-X analyzer (Keyence).

### Fluorescence and Immunofluorescence quantification

For quantification of inhibitory neurons in GFP-positive cells, we used *Scn1a*-GFP/*Vgat*-Cre/*Rosa26*-tdTomamto mice at 4W. We acquired multiple color images of primary motor cortex and hippocampus from three parasagittal sections per animal. Six images per region of interest were manually counted and summarized using Adobe Photoshop Elements 10 and Excel (Microsoft). On immunofluorescence quantification, we used *Scn1a*-GFP mice at P15 and/or 4W for the quantification of immunosignals. For quantification of GFP, NeuN, PV, SST, Nav1.1, Nav1.2, FEZF2 or TBR1-positive cells, we acquired multiple color images of primary motor cortex and hippocampus from three parasagittal sections per animal. Six ∼ nine images per region of interest were manually quantified and summarized. For quantification of PV, SST, FEZF2 or TBR1-positive cells, intensity and area size of GFP fluorescent signals were measured by Fiji software. Statistical analyses were performed by one-way ANOVA followed by Tukey–Kramer post-hoc multiple comparison test using Kyplot 6.0 (KyensLab Inc.). P-value smaller than 0.05 was considered statistically significant. Data are presented as the mean ± standard error of the mean (SEM).

### *In situ* hybridization

Frozen sections (30 µm) of PFA-fixed mouse brains at 4W were incubated in 0.3% H_2_O_2_ in PBS for 30 min at RT to quench endogenous peroxidases, and mounted on glass slides. The sections on slides were UV irradiated with 1,250 mJ/cm^2^ (Bio-Rad), permeabilized with 0.3% TritonX-100 in PBS for 15 min at RT, and digested with 1 µg/ml proteinase K (Nacalai Tesque) in 10mM Tris-HCl and 1mM EDTA, pH 8.0, for 30 min at 37°C, washed twice with 100 mM glycine in PBS for 5 min at RT, fixed with 4% formaldehyde in PBS for 5 min at RT, and acetylated with 0.25% acetic anhydride in 100 mM triethanolamine, pH8.0. After acetylation, the sections were washed twice with 0.1 M phosphate buffer, pH 8.0, incubated in a hybridization buffer [(50% formamide, 5× SSPE, 0.1% SDS, and 1 mg/ml Yeast tRNA (Roche Diagnostics)] containing the Avidin solution (Vector Laboratories) for 2 hr at 60°C, and hybridized with 2 µg/ml digoxigenin (DIG)- and dinitrophenol (DNP)-labeled probes in a hybridization buffer containing the Biotin solution (Vector Laboratories) overnight at 60°C in a humidified chamber. The hybridized sections were washed with 50% formamide in 2× SSC for 15 min at 50°C twice, incubated in TNE (1 mM EDTA, 500 mM NaCl, 10 mM Tris-HCl, pH 8.0) for 10 min at 37°C, treated with 20 µg/ml RNase A (Nacalai Tesque) in TNE for 15 min at 37°C, washed 2× SSC twice for 15 min each at 37°C twice and 0.2× SSC twice for 15 min each at 37°C. After washing twice in a high stringency buffer (10 mM Tris, 10 mM EDTA and 500 mM NaCl, pH 8.0) for 10 min each at RT, the sections were blocked with a blocking buffer [20 mM Tris and 150 mM NaCl, pH 7.5 containing 0.05% Tween-20, 4% BlockAce (DS Pharma Biomedical) and 0.5× Blocking reagent (Roche Diagnostics)] for 1 hr at RT, and incubated with the alkaline phosphatase-conjugated sheep anti-DIG (1:500; 11093274910, Roche Diagnostics) and biotinylated rabbit anti-DNP (1:100; BA-0603, Vector laboratories) antibodies in a blocking buffer overnight at 4°C, followed by incubation with the biotinylated goat anti-rabbit antibody (1:200; BA-1000, Vector laboratories) in a blocking buffer at RT for 1 to 2hr. The probes were visualized using the NBT/BCIP kit (Roche Diagnostics), VECTASTAIN Elite ABC kit (Vector laboratories) and ImmPACT DAB substrate (Vector laboratories).

The DIG-labeled RNA probes for *Scn1a* designed to target the 3’-untranlated region (nucleotides 6,488–7,102 from accession number NM_001313997.1) were described previously (Ogiwara et al., 2007), and synthesized using the MEGAscript transcription kits (Thermo Fisher Scientific) with DIG-11-UTP (Roche Diagnostics). The DNP-labeled RNA probes for GFP were derived from the fragment corresponding to nucleotides 1,256–1,983 in pIRES2-EGFP (Takara Bio), and prepared using the MEGAscript transcription kits (Thermo Fisher Scientific) with DNP-11-UTP (PerkinElmer).

## Acknowledgment

We thank Dr. Yaguchi (Laboratory for Behavioral Genetics) and the staff members at the Research Resources Division of RIKEN Center for Brain Science for technical assistance in generating *Scn1a*-GFP BAC Tg mice and Dr. Kaneda (Nippon Medical School) for his support. This study was supported in part by MEXT/JSPS KAKENHI JP20H03566, JP17H01564, JP16H06276, AMED JP18dm0107092, RIKEN Center for Brain Science (K.Y.); Takeda Science Foundation, Kiyokun Foundation (I.O. and K.Y.); MEXT/JSPS KAKENHI JP19790747, JP21791020, JP16K15564, JP19K08284 (I.O.); and Japan Epilepsy Research Foundation (I.O. and T.T.).

## Supplementary Information

**Supplementary figure S1.**
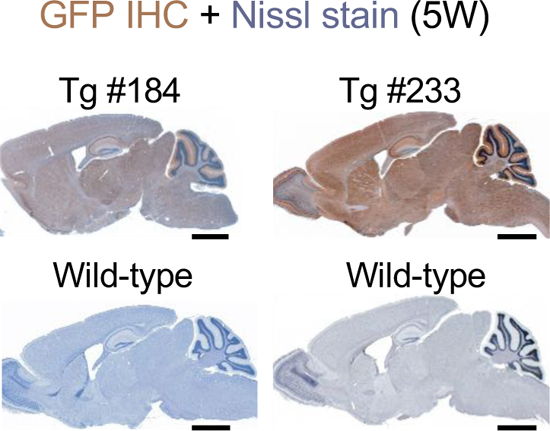
GFP expressions in brains of two *Scn1a*-GFP transgenic mouse lines. Chromogenic immunostaining of GFP in parasagittal sections from brains of 5W *Scn1a*-GFP mouse lines (#184, #233) and wild-type controls by anti-GFP antibody (brown). Sections were counterstained with Nissl for labeling of neurons (violet). IHC, immunohistochemistry. Scale bars; 1 mm.

**Supplementary figure S2.**
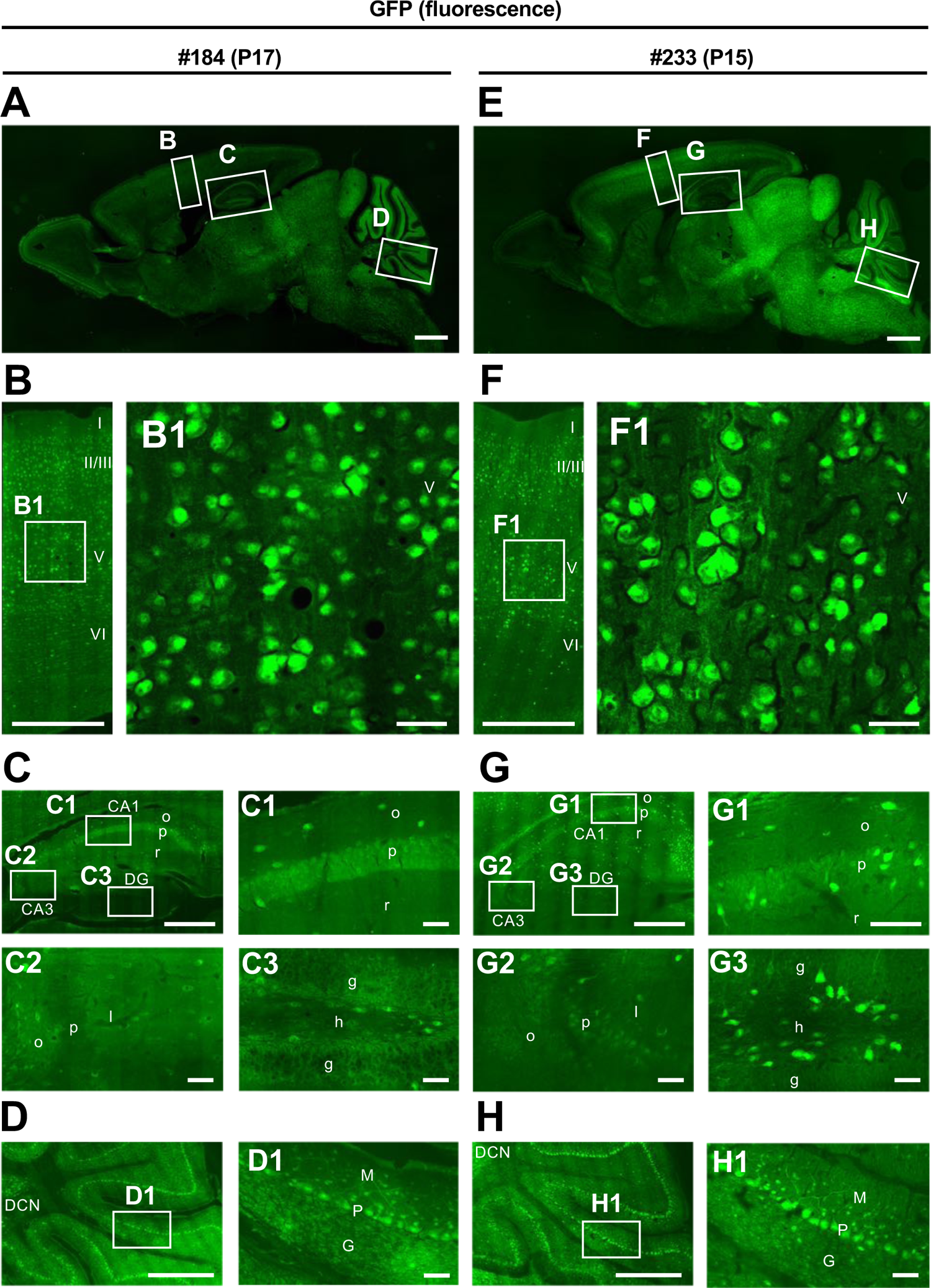
Similar GFP distribution in brains of two *Scn1a*-GFP mouse lines (#184, #233). Fluorescence images of parasagittal sections from brains of P15-17 *Scn1a*-GFP mouse lines #184 (**A-D**) and #233 (**E-H**). Magnified images outlined in (A, E) are shown in (B-D, F-H) and further magnified in (B1, C1∼3, D1, F1, G1∼3, H1). Although both lines #184 and #233 show a similar distribution of GFP-signals across all brain regions, the GFP signals are more intense in #233 than that in #184 in caudal part of the brain such as mid brain and brainstem. CA, cornu ammonis; DG, dentate gyrus; o, stratum oriens; p, stratum pyramidale; r, stratum radiatum; l, stratum lucidum; g, stratum granulosum; h, hilus; DCN, deep cerebellar nuclei; M, molecular layer; P, Purkinje cell layer; G, granular cell layer. Scale bars; 1 mm (A, E), 500 µm (B-D, F-H), 50 µm (B1, C1-3, D1, F1, G1-3, H1).

**Supplementary figure S3.**
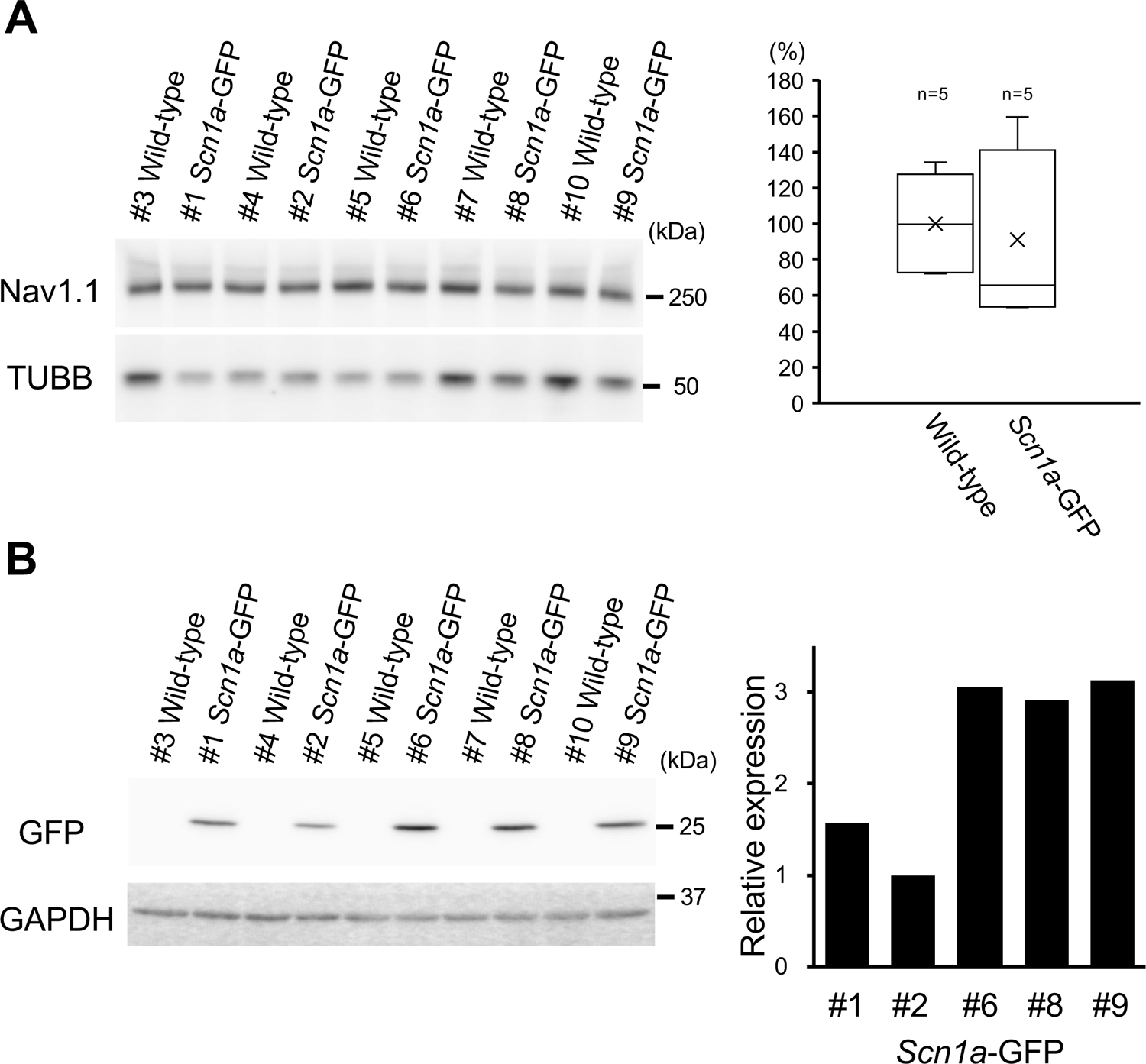
Expression levels of Nav1.1 are stable but that of GFP are rather variable among *Scn1a*-GFP individual mice. (**A**) Western blot analysis for Nav1.1 in *Scn1a*-GFP and wild-type mice. The whole membrane fractions from 5W *Scn1a*-GFP brains (line #233) were probed with anti-Nav1.1 antibody (left panel). Nav1.1 values quantified from the blots were normalized by beta-tubulin (TUBB), and the mean of Nav1.1 amount in wild-type was assigned as a value of 100 (right panel). The amount of Nav1.1 was not significantly changed in the *Scn1a*-GFP mice compared to wild-type mice (t-test; p > 0.05). The boxes show median, 25th and 75th percentiles, and whiskers represent minimum and maximum values. Cross marks indicate mean values for each genotype. (**B**) Western blot analysis for GFP in *Scn1a*-GFP and wild-type mice. The whole cytosolic fractions from the same series of mice used in (A) were probed with anti-GFP antibody (left panel). GFP values were normalized by glyceraldehyde 3-phosphate dehydrogenase (GAPDH), and the lowest GFP amount in the #2 mouse was assigned as a value of 1 (right panel). Statistical significance was assessed using t-test.

**Supplementary figure S4.**
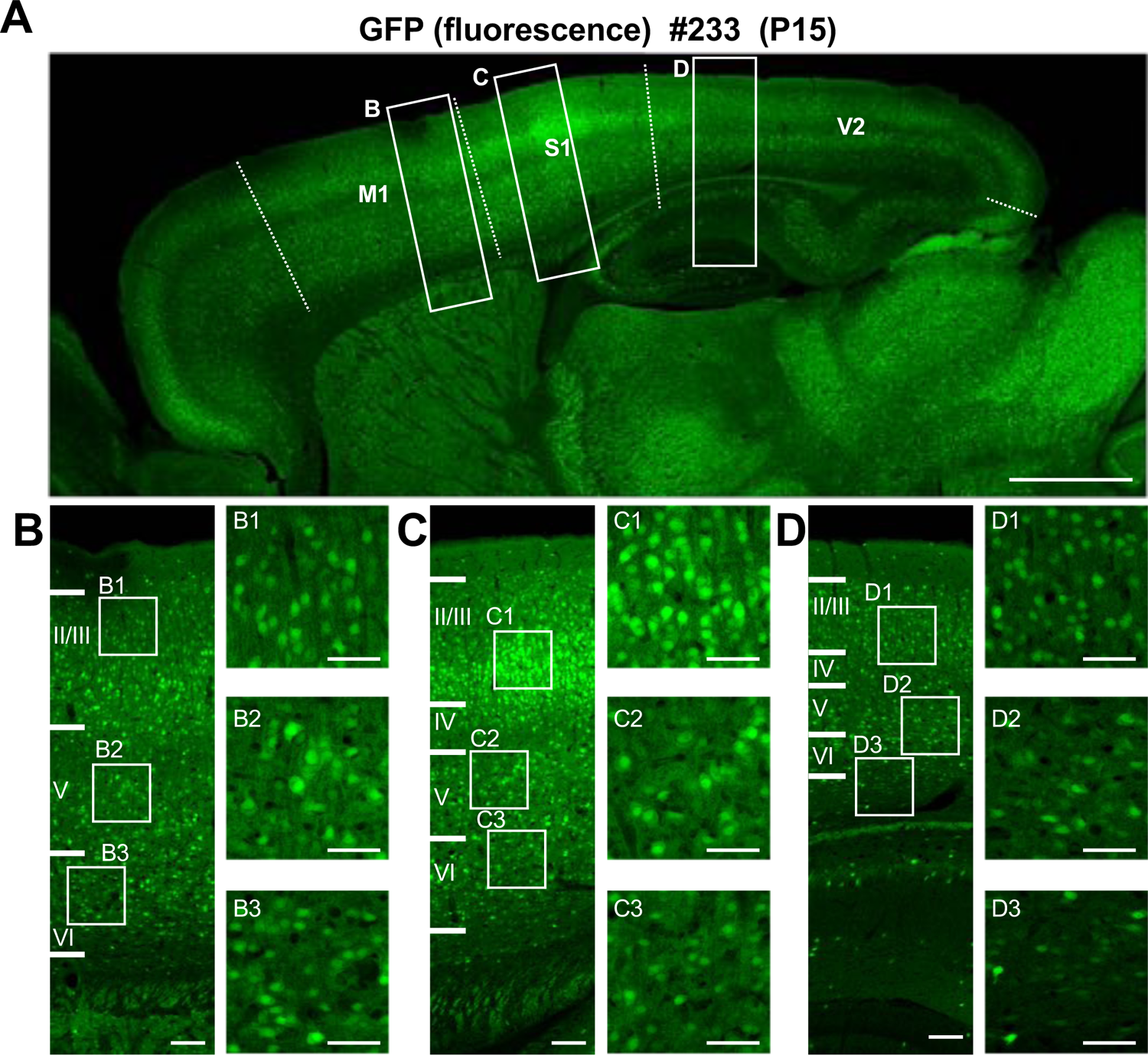
Distribution of GFP fluorescent signals in neocortical layers of *Scn1a*-GFP mouse brain. (**A**) Fluorescent images of parasagittal section spanning whole neocortical area of P15 *Scn1a*-GFP brain (line #233). (**B-D**) Outlined areas in (A) are magnified in (B-D) and further in (B1-B3, C1-C2, D1-D3). GFP signals in L2/3 of primary somatosensory cortex (C) are brighter than other regions such as primary motor cortex (B) and secondary visual cortex (D). Brain regions were defined using mouse brain atlas (Paxinos and Franklin, 2001) as reference. Dashed lines indicate the borders between M1, S1 and V2 areas. M1, primary motor cortex; S1, primary somatosensory cortex; V2, secondary visual cortex. Scale bars; 1 mm (A), 100 µm (B-D), 50 µm (B1-B3, C1-C3, D1-D3).

**Supplementary figure S5.**
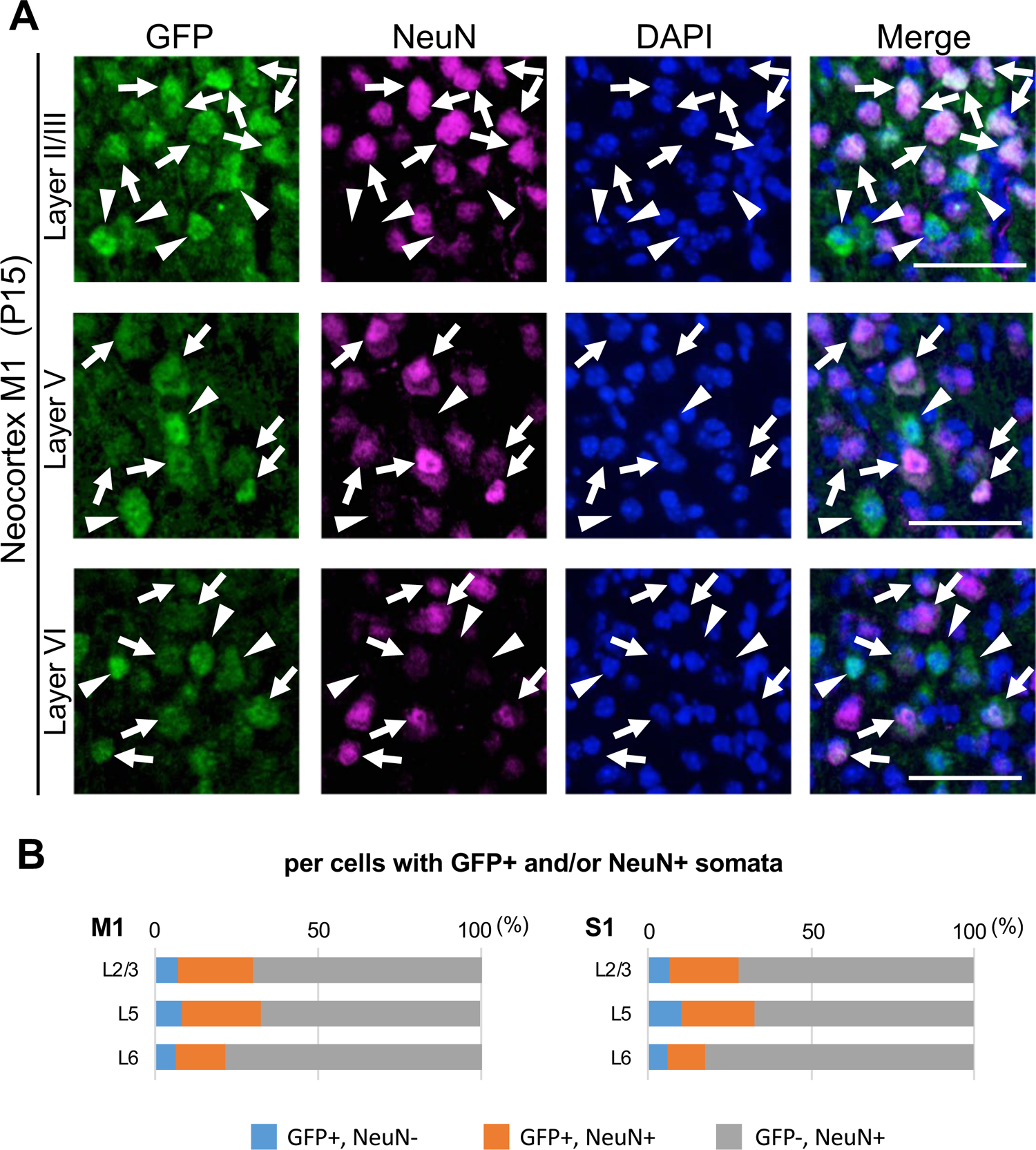
Immunostaining of NeuN and GFP in the neocortex of *Scn1a*-GFP mouse. (**A**) Double immunostaining of parasagittal sections from P15 *Scn1a*-GFP mouse brain (line #233) by mouse anti-GFP (green) and mouse anti-NeuN (magenta) antibodies with DAPI (blue). Merged images are shown in the right panels. Arrows indicate GFP/NeuN-double positive cells. Arrowheads indicate GFP-positive but NeuN-negative cells. Note that cells with intense GFP-signals, which are assumed to be inhibitory neurons (see Figure 8), were NeuN-negative. All images are oriented from pial surface (top) to callosal (bottom). Scale µm. (**B**) Bar graphs indicating the percentage of cells with GFP-positive somata among cells with GFP- and/or NeuN-positive somata in neocortical layers. Cells in primary motor cortex (M1) and primary somatosensory cortex (S1) of P15 *Scn1a*-GFP mice were counted (see also Supplementary table S1). L2/3, L5, L6: neocortical layer II/III, V, VI. +, positive; -, negative.

**Supplementary figure S6.**
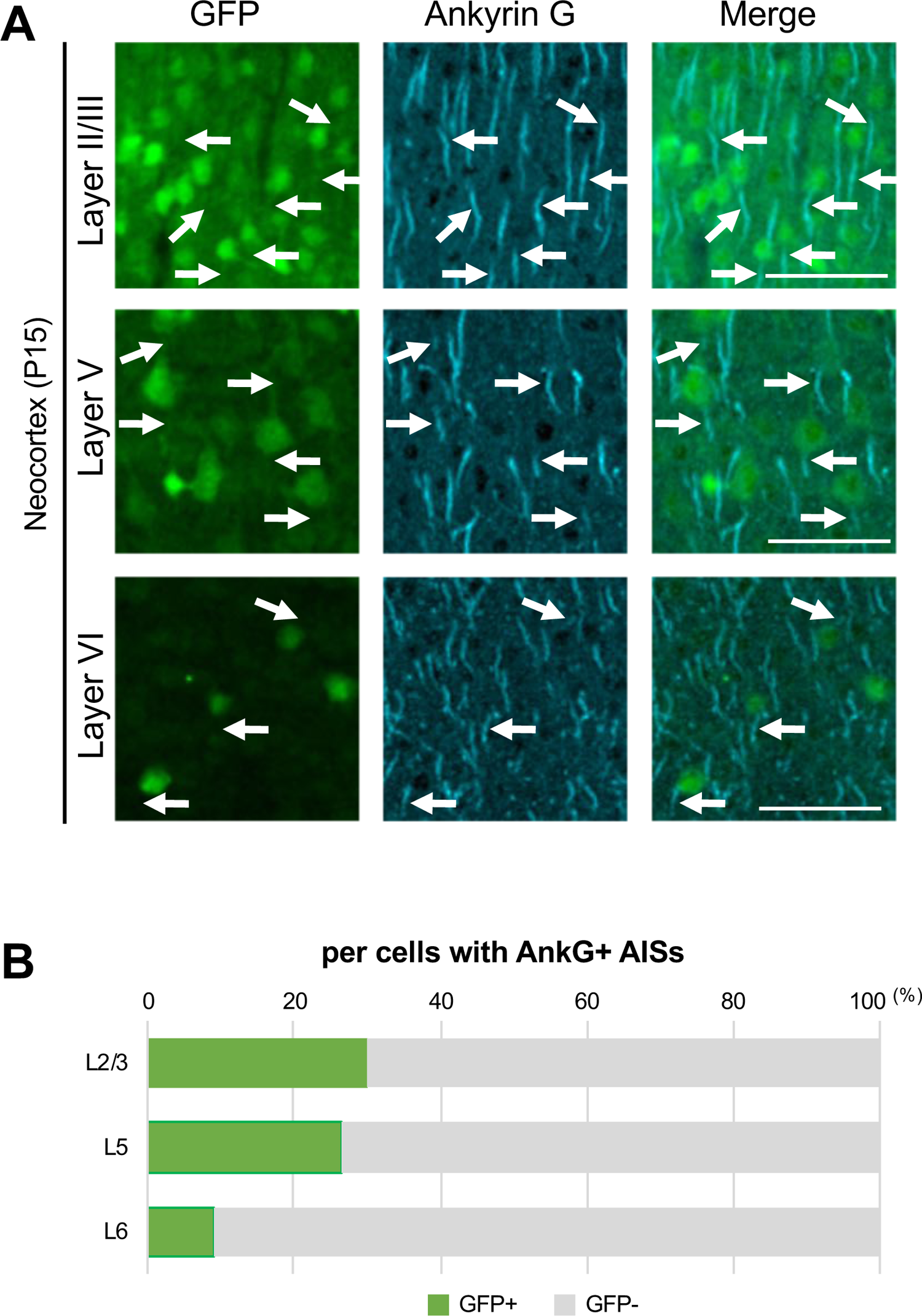
Immunostaining of GFP and ankyrinG in the neocortex of *Scn1a*-GFP mouse. (A) Double immunostaining of parasagittal sections from P15 *Scn1a*-GFP mouse brain (line #233) by mouse anti-GFP (green) and goat anti-ankyrinG (cyan) antibodies. Merged images are shown in the right panels. Arrows indicate ankyrinG-positive AISs with GFP-positive soma. All images are oriented from pial surface (top) to callosal (bottom). Scale bars; 50μ (B) Bar graphs indicating the percentage of cells with GFP-positive/negative somata per cells with ankyrinG-positive AISs in L2/3, L5, and L6 (see also Supplementary table S5). Note that 30% (L2/3), 26% (L5) and 9% (L6) of cells with ankyrinG-positive AISs have GFP-positive somata. L2/3, L5, L6: neocortical layer II/III, V, VI. AnkG, ankyrinG; +, positive; -, negative.

**Supplementary figure S7.**
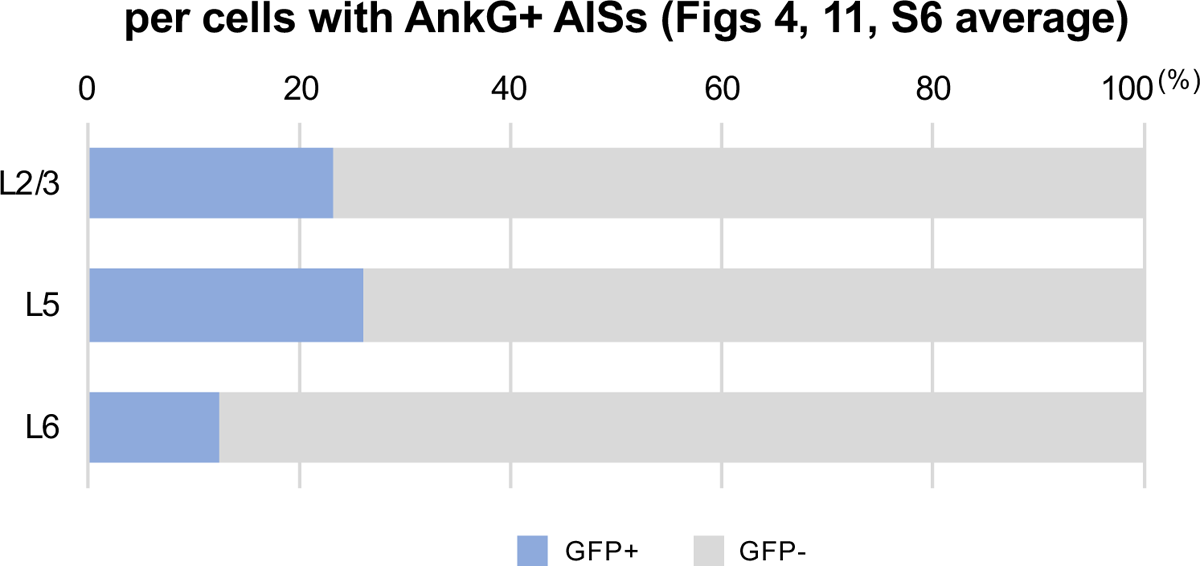
GFP-positive cells are more abundant in layers II/III and V than in layer VI of the neocortex. Bar graphs indicating the average percentage of cells with GFP-positive/negative somata per cells with ankyrinG-positive AISs in three different assessments (see also Supplementary table S6). Note that 23% (L2/3), 26% (L5) and 13% (L6) of cells with ankyrinG positive AISs have GFP-positive somata. L2/3, L5, L6: neocortical layer II/III, V, VI. AnkG, ankyrinG; +, positive; -, negative.

**Supplementary figure S8.**
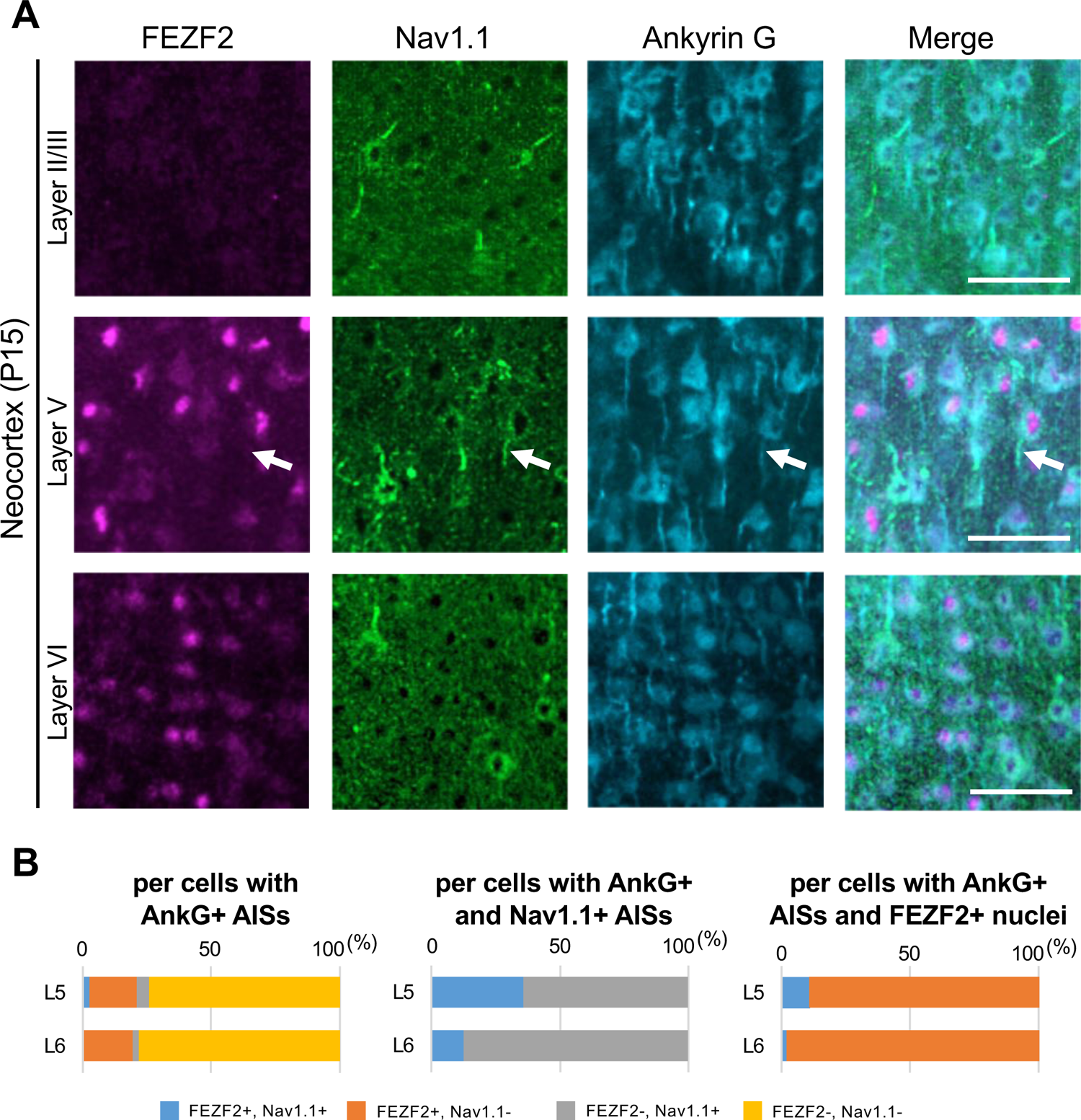
A part of FEZF2-positive cells at L5 have Nav1.1-positive AIS of *Scn1a*-GFP mouse neocortex. (**A**) Triple immunostaining of parasagittal section from P15 *Scn1a*-GFP mouse brain (line #233) by rabbit anti-FEZF2 (magenta), goat anti-Nav1.1 (green), and mouse anti-ankyrinG (cyan) antibodies. Merged images are shown in the right panels. An arrow indicates Nav1.1-positive AIS of FEZF2-positive cell. All images are oriented from pial surface (top) µm. (**B**) Bar graphs indicating the percentage of cells with FEZF2- and Nav1.1-positive/negative nuclei or AISs per cells with ankyrinG-positive AISs (left panel) (see also Supplementary table S25), the percentage of cells with FEZF2-positive/negative nuclei per cells with ankyrinG and Nav1.1-positive AISs (middle panel) (see also Supplementary table S26), and the percentage of cells with Nav1.1-positive/negative AISs per cells with ankyrinG-positive AISs and FEZF2-positive nuclei (right panel) (see also Supplementary table S27). Cells in primary motor cortex of *Scn1a*-GFP mouse at P15 were counted. L5, L6: neocortical layer V, VI. AnkG, ankyrinG; +, positive; -, negative.

**Supplementary figure S9.**
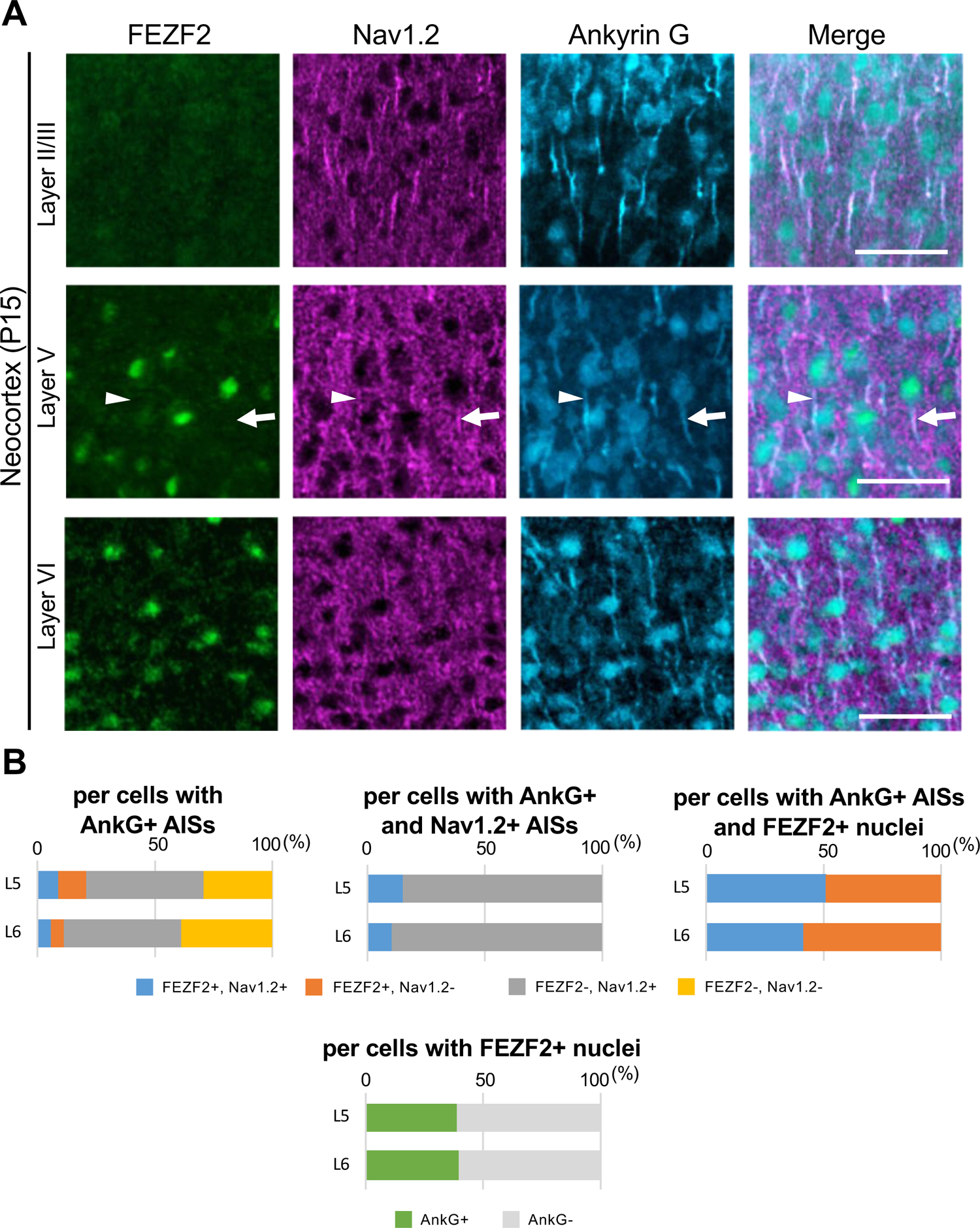
A half of FEZF2-positive cells at L5 have Nav1.2-positive AIS of *Scn1a*-GFP mouse neocortex. (**A**) Triple immunostaining of parasagittal section from P15 *Scn1a*-GFP mouse brain (line #233) by rabbit anti-FEZF2 (green), goat anti-Nav1.2 (magenta), and mouse anti-ankyrinG (cyan) antibodies. Merged images are shown in the right panels. An arrow indicates FEZF2/Nav1.2-double positive cell. An arrowhead indicates Nav1.2-negative AIS of FEZF2-positive cell. All images are oriented from pial surface (top) to callosal (bottom). Scale bars; 50 μm. (**B**) Bar graphs indicating the percentage of cells with FEZF2 and Nav1.2-positive/negative nuclei or AISs per cells with ankyrinG-positive AISs (left-upper panel) (see also Supplementary table S28), the percentage of cells with FEZF2-positive/negative nuclei per cells with ankyrinG and Nav1.2-positive AISs (middle-upper panel) (see also Supplementary table S29), and the percentage of cells with Nav1.2-positive/negative AISs per cells with ankyrinG-positive AISs and FEZF2-positive nuclei (right-upper panel) (see also Supplementary table S30). Cells with ankyrinG-positive/negative AISs per cells with FEZF2-positive nuclei (lower panel) (see also Supplementary table S31). Cells in primary motor cortex of *Scn1a*-GFP mouse at P15 were counted. L5, L6: neocortical layer V, VI. +; positive. -; negative.

**Supplementary figure S10.**
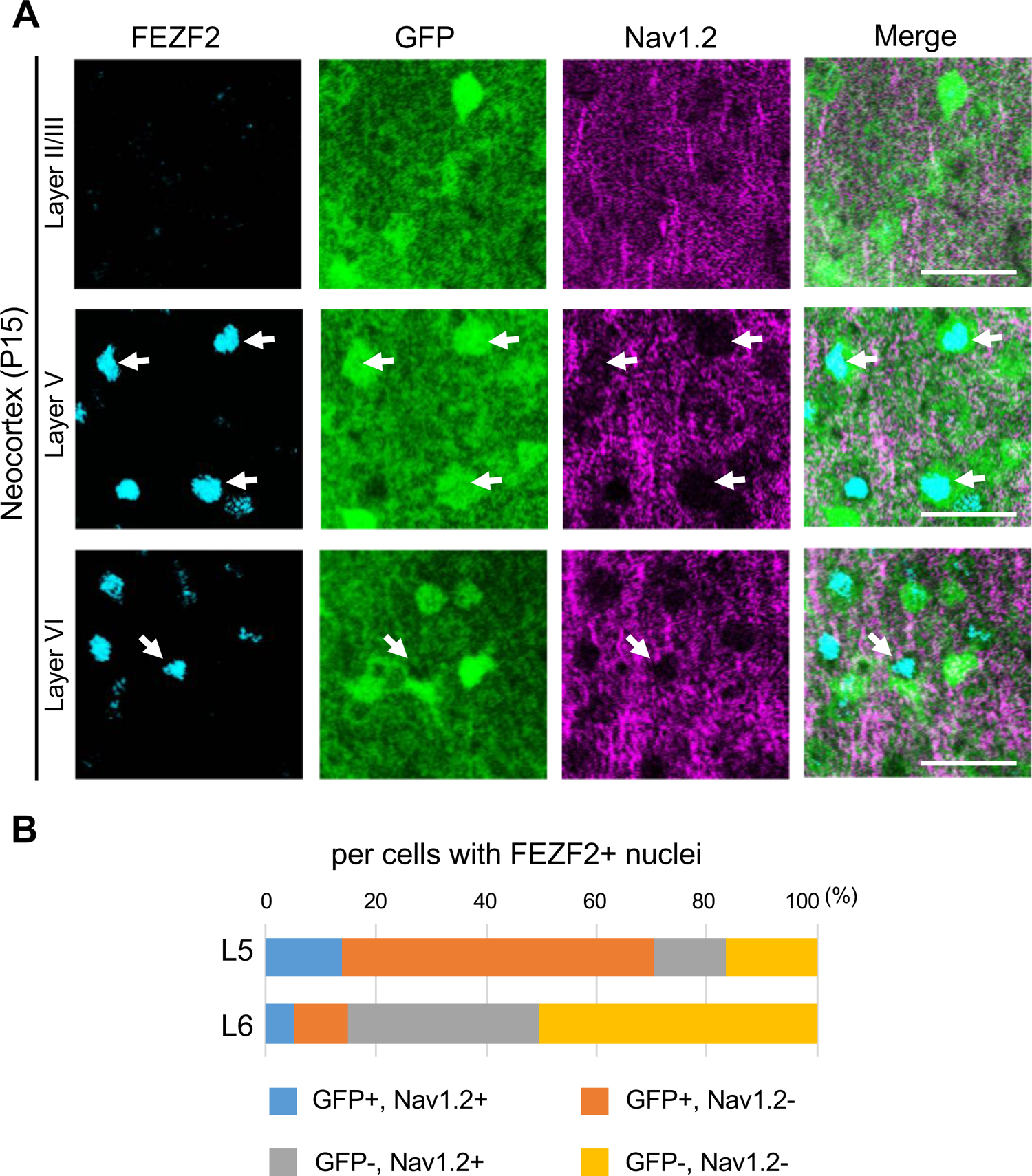
Most of the FEZF2/GFP-double positive cells are Nav1.2-negative at L5 in *Scn1a*-GFP mouse neocortex. (**A**) Triple immunostaining of parasagittal sections from P15 *Scn1a*-GFP mouse brain (line #233) by rabbit anti-FEZF2 (cyan), mouse anti-GFP (green), and goat anti-Nav1.2 (magenta) antibodies. Merged images are shown in the right panels. Arrows indicate FEZF2/GFP-double positive cells (cells with GFP-positive somata and FEZF2-positive nuclei). Note that Nav1.2-positive AISs do not belong to GFP/FEZF2-double positive cells. All images are oriented from pial surface (top) to callosal (bottom). Scale bars; 25 µm. (**B**) Bar graphs indicating the percentage of cells with GFP- and Nav1.2-positive/negative somata and AISs per cells with FEZF2-positive nuclei in L5 and L6 (see also Supplementary table S32). For this graph, to obtain correct cell population for Nav1.2-negative cells, virtual cell numbers were estimated using percentage of ankyrinG/FEZF2-double positive cells in Supplementary figure S9B-lower panel (see also Supplementary table S28). Cells in primary motor cortex of *Scn1a*-GFP mouse at P15 were counted. L5, L6: neocortical layer V, VI. +, positive; -, negative.

**Supplementary figure S11.**
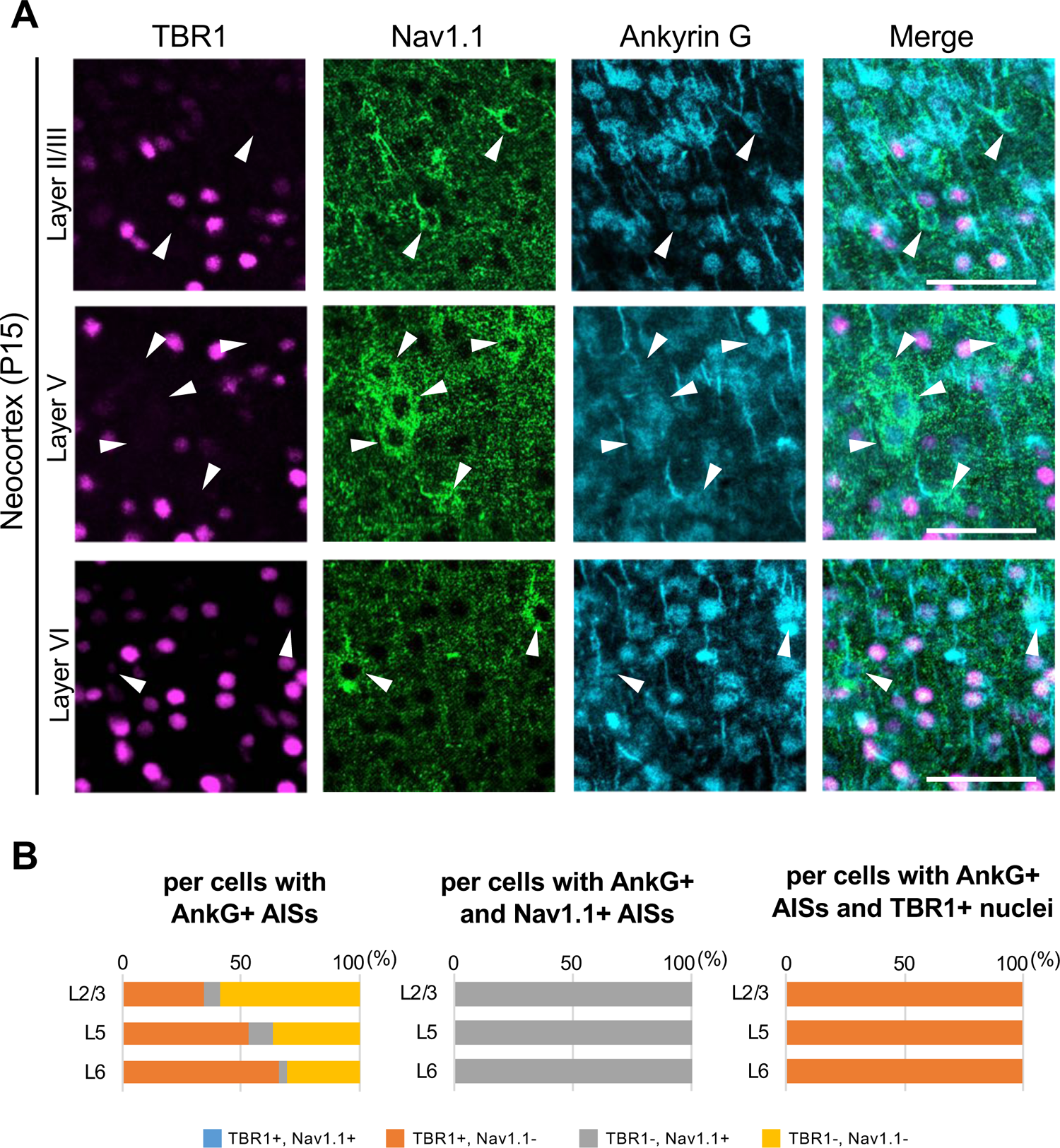
AISs of TBR1-positive cells are Nav1.1-negative in *Scn1a*-GFP mouse neocortex. (**A**) Triple immunostaining of parasagittal section from P15 *Scn1a*-GFP mouse brain (line #233) by rabbit anti-TBR1 (magenta), goat anti-Nav1.1 (green), and mouse anti-ankyrinG (cyan) antibodies. The regions of motor cortex were shown. Merged images are shown in the right panels. Arrowheads indicate Nav1.1-positive cells. All images are oriented from pial surface (top) to callosal (bottom). Scale bars; 50 μm. (**B**) Bar graphs indicating the percentage of cells with TBR1- and Nav1.1-positive/negative nuclei or AISs per cells with ankyrinG-positive AISs (left panel) (see also Supplementary table S37), the percentage of cells with TBR1-positive/negative nuclei per cells with ankyrinG- and Nav1.1-positive AISs (middle panel) (see also Supplementary table S38), and the percentage of cells with Nav1.1-positive/negative AISs per cells with ankyrinG-positive AISs and TBR1-positive nuclei (right panel) (see also Supplementary table S39). Cells in primary motor cortex of *Scn1a*-GFP mouse at P15 were counted. L2/3, L5, L6: neocortical layer II/III, V, VI. AnkG, ankyrinG; +, positive; -, negative.

**Supplementary figure S12.**
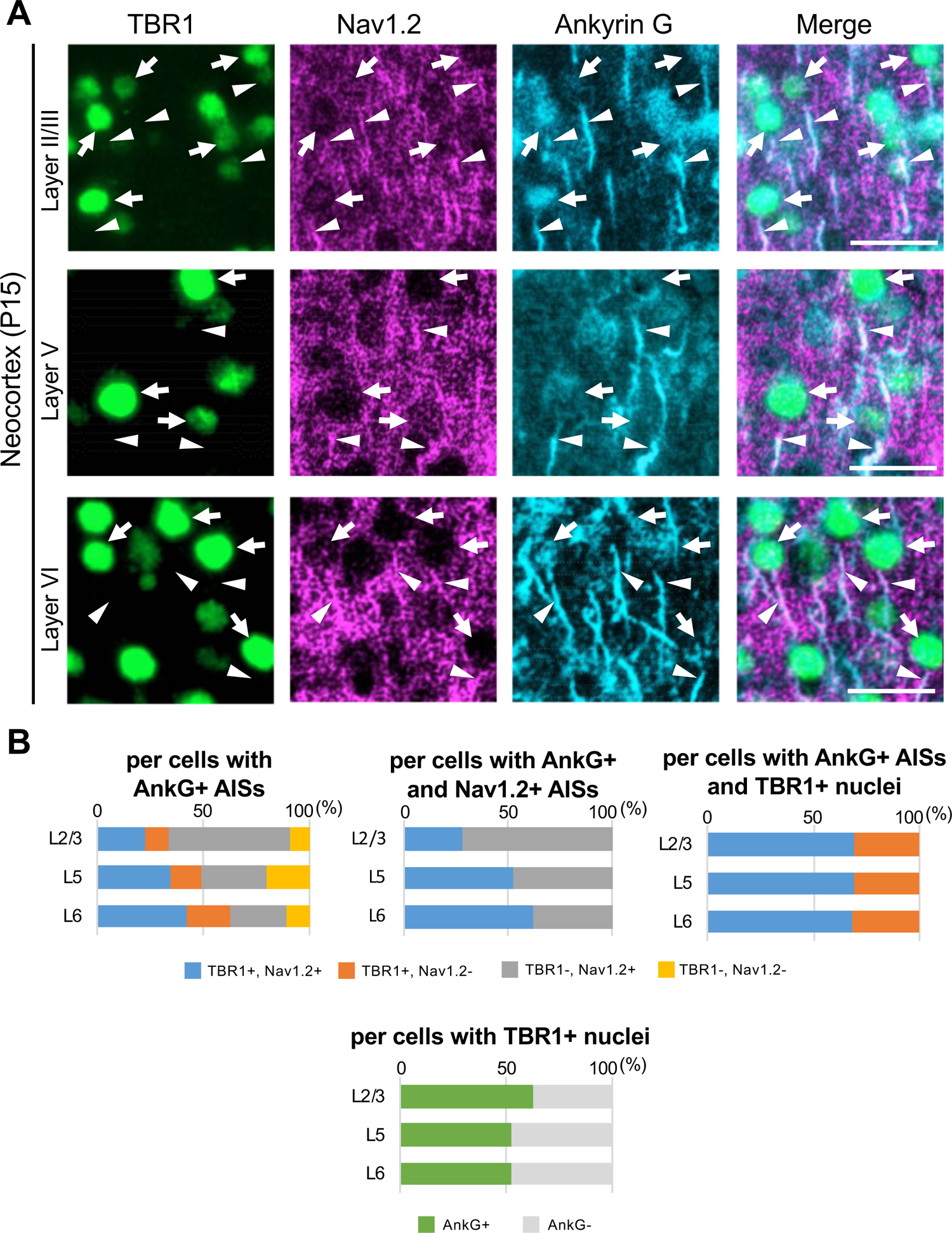
70% of TBR1-positive cells have Nav1.2-positive AIS in *Scn1a*-GFP mouse neocortex. (**A**) Triple immunostaining of parasagittal section from P15 *Scn1a*-GFP mouse brain (line #233) by rabbit anti-TBR1 (green), goat anti-Nav1.2 (magenta), and mouse anti-ankyrinG (cyan) antibodies. Merged images are shown in the right panels. Arrows indicate TBR1/ankyrinG-double positive cells. Arrowheads indicate Nav1.2-positive AIS of TBR1-positive cells. All images are oriented from pial surface (top) to callosal (bottom). Scale bars; 25 μm. (**B**) Bar graphs indicating the percentage of cells with TBR1- and Nav1.2-positive/negative nuclei or AISs per cells with ankyrinG-positive AISs (left-upper panel) (see also Supplementary table S40), the percentage of cells with TBR1-positive/negative nuclei per cells with ankyrinG- and Nav1.2-positive AISs (middle-upper panel) (see also Supplementary table S41), and the percentage of cells with Nav1.2-positive/negative AISs per cells with ankyrinG-positive AISs and TBR1-positive nuclei (right-upper panel) (see also Supplementary table S42). Cells with ankyrinG-positive/negative AISs per cells with TBR1-positive nuclei (lower panel) (see also Supplementary table S43). Cells in primary motor cortex of *Scn1a*-GFP mouse at P15 were counted. L2/3, L5, L6: neocortical layer II/III, V, VI. AnkG, ankyrinG; +, positive; -, negative.

**Supplementary figure S13.**
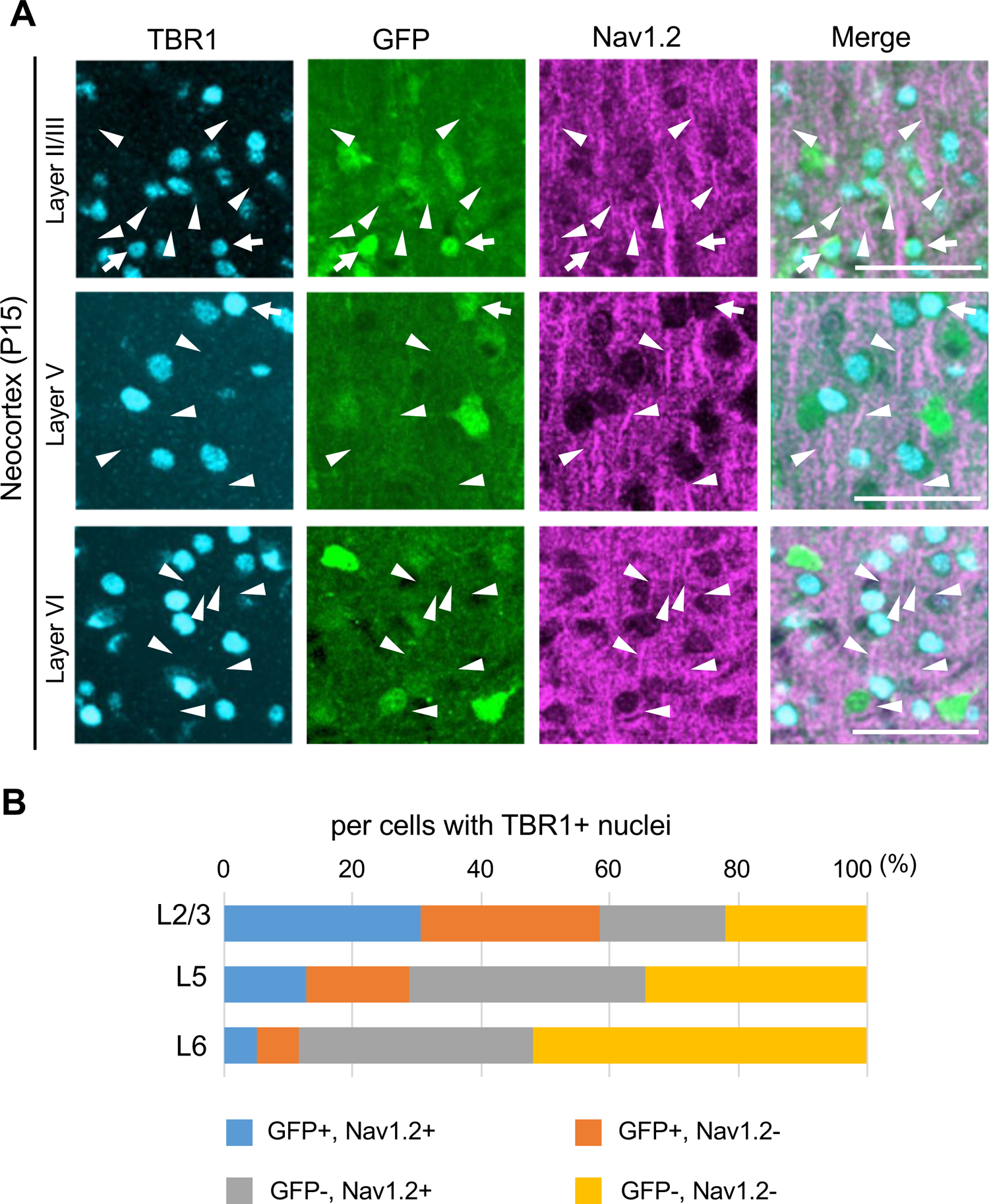
TBR1/Nav1.2-double positive cells are GFP-negative at L5 and L6 in *Scn1a*-GFP mouse neocortex. (**A**) Triple immunostaining of parasagittal section from P15 *Scn1a*-GFP mouse brain (line #233) by rabbit anti-TBR1 (cyan), mouse anti-GFP (green), and goat anti-Nav1.2 (magenta) antibodies. Merged images are shown in the right panels. Arrows indicate TBR1/GFP-double positive cells. Arrowheads indicate Nav1.2-positive AIS of TBR1-positive cells. All images are oriented from pial surface (top) to callosal (bottom). Scale bar; 50 µm. (**B**) Bar graphs indicating the percentage of cells with GFP- and Nav1.2-positive/negative somata and AISs per cells with TBR1-positive nuclei in L2/3, L5, and L6 (see also Supplementary table S44). For this graph, to obtain correct cell population for Nav1.2-negative cells, virtual cell numbers were estimated using percentage of ankyrinG/TBR1-double positive cells in Supplementary figure S12B-lower panel (see also Supplementary table S40). Cells in primary motor cortex of *Scn1a*-GFP mouse at P15 were counted. L2/3, L5, L6: neocortical layer II/III, V, VI. +, positive; -, negative.

**Supplementary figure S14.**
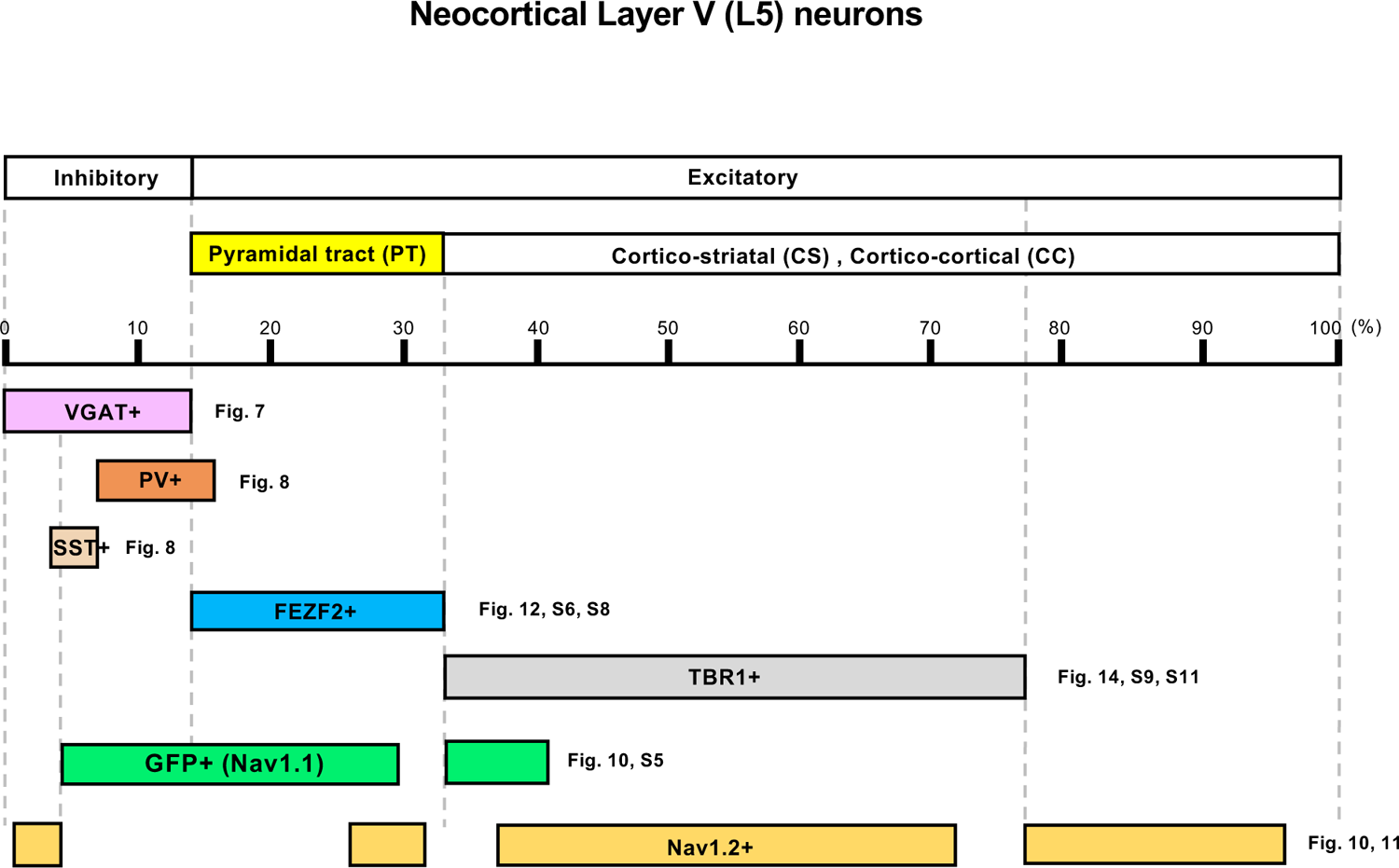
Distributions of Nav1.1 (GFP) and Nav1.2 in neocortical layer V revealed by the analysis of *Scn1a*-GFP mouse.

**Supplementary table S1.** Percentage of cells with GFP- and/or NeuN-positive somata among cells with GFP- and/or NeuN-positive somata in *Scn1a*-GFP mouse neocortex. Values for Supplementary figure S5B. ^a^M1, S1, L2/3, L5, L6: primary motor cortex, primary somatosensory cortex, neocortical layer II/III, V, VI. Cells in three *Scn1a*-GFP mice (line #233) at P15 were counted; ^b^L2/3 (M1; N = 2599 cells, S1; N=2932), L5 (M1; N = 1734 cells, S1; N=1827) and L6 (M1; N = 2465 cells, S1; N=2335). Values are presented as mean ± SEM. +, positive; -, negative.

| per cells with GFP+ and/or NeuN+ somata |  |  |  |  |
| --- | --- | --- | --- | --- |
| Region <sup>a</sup> |  | % GFP+, NeuN- <sup>b</sup> | % GFP+, NeuN+ <sup>b</sup> | % GFP-, NeuN+ <sup>b</sup> |
| M1 | L2/3 | 6.8 ± 0.5 | 23.3 ± 0.5 | 70.2 ± 0.5 |
|  | L5 | 8.4 ± 0.6 | 23.8 ± 0.1 | 67.7 ± 1.3 |
|  | L6 | 6.5 ± 0.6 | 15.2 ± 2.2 | 78.3 ± 2.3 |
| S1 | L2/3 | 6.7 ± 0.1 | 21.3 ± 2.0 | 72.0 ± 2.1 |
|  | L5 | 10.2 ± 0.9 | 22.6 ± 2.1 | 67.1 ± 1.3 |
|  | L6 | 5.9 ± 0.7 | 11.7 ± 1.8 | 82.4 ± 1.8 |

**Supplementary table S2.**
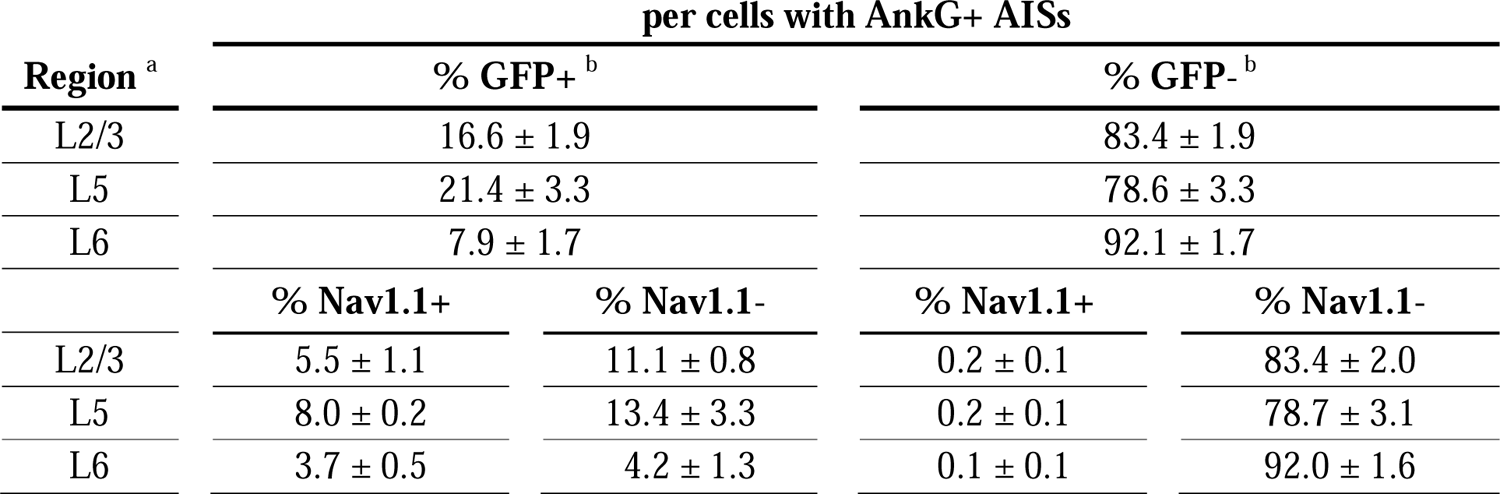
Percentage of cells with GFP- and Nav1.1-positive/negative somata and AISs per cells with ankyrinG-positive AISs in *Scn1a*-GFP mouse neocortex. Values for Figure 4D-left panel. ^a^L2/3, L5, L6: neocortical layer II/III, V, VI. Cells in three *Scn1a*-GFP mice (line #233) at P15 were counted; ^b^L2/3 (N = 1412 cells), L5 (N = 1054 cells), and L6 (N = 1481 cells). Values are presented as mean ± SEM. AnkG, ankyrinG; +, positive; -, negative.

**Supplementary table S3.** Percentage of cells with GFP-positive/negative somata per cells with ankyrinG-positive AISs and Nav1.1-positive somata and/or AISs in *Scn1a*-GFP mouse neocortex. Values for Figure 4D-middle panel. ^a^L2/3, L5, L6: neocortical layer II/III, V, VI. Cells in three *Scn1a*-GFP mice (line #233) at P15 were counted; ^b^L2/3 (N = 108 cells), L5 (N = 124 cells), and L6 (N = 82 cells). Values are presented as mean ± SEM. AnkG, ankyrinG; +, positive; -, negative.

| Region <sup>a</sup> | per cells with AnkG+ AISs and Nav1.1+ somata and/or AISs |  |
| --- | --- | --- |
|  | % GFP+ <sup>b</sup> | % GFP- <sup>b</sup> |
| L2/3 | 98.5 ± 0.8 | 1.55 ± 0.8 |
| L5 | 98.7 ± 0.6 | 1.27 ± 0.6 |
| L6 | 97.2 ± 2.8 | 2.82 ± 2.8 |

**Supplementary table S4.** Percentage of cells with GFP-positive/negative somata per cells with ankyrinG-positive AISs and Nav1.1+ somata and/or AISs in *Scn1a*-GFP mouse neocortex. Values for Figure 4D-right panel. ^a^L2/3, L5, L6: neocortical layer II/III, V, VI. Cells in three *Scn1a*-GFP mice (line #233) at P15 were counted; ^b^L2/3 (N = 229 cells), L5 (N = 220 cells), and L6 (N = 111 cells). Values are presented as mean ± SEM. AnkG, ankyrinG; +, positive; -, negative.

| Region <sup>a</sup> | per cells with AnkG+ AISs and GFP+ somata |  |
| --- | --- | --- |
|  | % Nav1.1+ <sup>b</sup> | % Nav1.1- <sup>b</sup> |
| L2/3 | 32.8 ± 3.5 | 67.2 ± 3.5 |
| L5 | 39.9 ± 6.4 | 60.1 ± 6.4 |
| L6 | 49.5 ± 8.4 | 50.5 ± 8.4 |

**Supplementary table S5.** Percentage of cells with GFP-positive/negative somata per cells with ankyrinG-positive AISs in *Scn1a*-GFP mouse neocortex. Values for Supplementary figure S6B. ^a^L2/3, L5, L6: neocortical layer II/III, V, VI. Cells in three *Scn1a*-GFP mice (line #233) at P15 were counted; ^b^L2/3 (N = 1787 cells), L5 (N = 1450 cells), and L6 (N = 2330 cells). Values are presented as mean ± SEM. AnkG, ankyrinG; +, positive; -, negative.

| Region <sup>a</sup> | per cells with AnkG+ AISs |  |
| --- | --- | --- |
|  | % GFP+ <sup>b</sup> | % GFP- <sup>b</sup> |
| L2/3 | 30.0 ± 1.1 | 70.0 ± 1.1 |
| L5 | 26.4 ± 1.6 | 73.6 ± 1.6 |
| L6 | 9.0 ± 1.0 | 91.0 ± 1.0 |

**Supplementary table S6.** Percentage of cells with GFP-positive somata per cells with ankyrinG-positive AISs in the three different assessments. Values for Supplementary figure S7. ^a^L2/3, L5, L6: neocortical layer II/III, V, VI. ^b^Values indicate that percentage of GFP positive cells in Supplementary figure S6B, Figure 4D-left panel and Figure 11B-left panel, respectively. AnkG, ankyrinG; +, positive; -, negative.

| Region <sup>a</sup> | % GFP+/AnkG+ AIS <sup>b</sup> | Nav1.1/GFP/AnkG | Nav1.2/GFP/AnkG | Average | SEM |
| --- | --- | --- | --- | --- | --- |
|  |  | % GFP+/AnkG+ AIS <sup>b</sup> | % GFP+/AnkG+ AIS <sup>b</sup> |  |  |
| L2/3 | 30.0 | 16.6 | 22.6 | 23.1 | 3.9 |
| L5 | 26.4 | 21.4 | 30.2 | 26.0 | 2.5 |
| L6 | 9.0 | 7.9 | 20.7 | 12.5 | 4.1 |

**Supplementary table S7.** Percentage of cells with Nav1.1-positive/negative AISs per cells with GFP-positive somata and ankyrinG-positive AISs in *Scn1a*-GFP mouse hippocampus. Values for Figure 5F-left panel. ^a^CA1: Cornu ammonis 1. ^b^Cells in three *Scn1a*-GFP mice (line #233) at P15 were counted (N = 117 cells). Values are presented as mean ± SEM. AnkG, ankyrinG; +, positive; -, negative.

| Region <sup>a</sup> | per cells with GFP+ somata and AnkG+ AISs |  |
| --- | --- | --- |
|  | % Nav1.1+ <sup>b</sup> | % Nav1.1- <sup>b</sup> |
| CA1 | 98.3 ± 0.9 | 1.7 ± 0.9 |

**Supplementary table S8.** Percentage of cells with GFP-positive/negative somata per cells with Nav1.1/ankyrinG-double positive AISs in *Scn1a*-GFP mouse hippocampus. Values for Figure 5F-right panel. ^a^CA1: Cornu ammonis 1. ^b^Cells in three *Scn1a*-GFP mice (line #233) at P15 were counted (N = 114 cells). Values are presented as mean ± SEM. AnkG, ankyrinG; +, positive; -, negative.

| Region <sup>a</sup> | per cells with Nav1.1+ and AnkG+ AISs |  |
| --- | --- | --- |
|  | % GFP+ <sup>b</sup> | % GFP- <sup>b</sup> |
| CA1 | 100.0 ± 0.0 | 0.0 ± 0.0 |

**Supplementary table S9.** Percentage of cells with Tomato-positive/negative somata per cells with GFP-positive somata in *Scn1a*-GFP/*Vgat*-Cre/*Rosa26*-tdTomato mouse neocortex and hippocampus. Values for Figure 7C-left panels. ^a^L2/3, L5, L6, CA1, DG: neocortical layer II/III, V, VI, cornu ammonis 1, dentate gyrus. Cells in two *Scn1a*-GFP/*Vgat*-Cre/*Rosa26*-tdTomato mice (line #233) at 4W were counted; ^b^L2/3 (N = 882 cells), L5 (N = 693 cells), L6 (N = 590 cells), CA1 (N = 270 cells) and DG (N = 191 cells). Values are presented as mean ± SEM. +, positive; -, negative.

| Region <sup>a</sup> | per cells with GFP+ somata |  |
| --- | --- | --- |
|  | %Tomato+ <sup>b</sup> | % Tomato- <sup>b</sup> |
| L2/3 | 23.2 ± 2.4 | 76.8 ± 2.4 |
| L5 | 28.1 ± 0.8 | 71.9 ± 0.8 |
| L6 | 27.4 ± 3.3 | 72.6 ± 3.3 |
| CA1 | 97.9 ± 0.1 | 2.1 ± 0.1 |
| DG | 93.5 ± 4.5 | 6.5 ± 4.5 |

**Supplementary table S10.** Percentage of cells with GFP-positive/negative somata per cells with Tomato-positive somata in *Scn1a*-GFP/*Vgat*-Cre/*Rosa26*-tdTomato mouse neocortex and hippocampus. Values for Figure 7C-right panels. ^a^L2/3, L5, L6, CA1, DG: neocortical layer II/III, V, VI, cornu ammonis 1, dentate gyrus. Cells in two *Scn1a*-GFP/*Vgat*-Cre/*Rosa26*-tdTomato mice (line #233) at 4W were counted; ^b^L2/3 (N = 263 cells), L5 (N = 224 cells), L6 (N = 300 cells), CA1 (N = 276 cells) and DG (N = 364 cells). Values are presented as mean ± SEM. +, positive; -, negative.

| Region <sup>a</sup> | per cells with Tomato+ somata |  |
| --- | --- | --- |
|  | %GFP+ <sup>b</sup> | % GFP- <sup>b</sup> |
| L2/3 | 72.7 ± 5.0 | 27.3 ± 5.0 |
| L5 | 76.9 ± 0.0 | 23.1 ± 0.0 |
| L6 | 83.0 ± 5.5 | 17.0 ± 5.5 |
| CA1 | 93.2 ± 0.2 | 6.8 ± 0.2 |
| DG | 77.2 ± 5.2 | 22.8 ± 5.2 |

**Supplementary table S11.** Percentage of cells with PV-positive/negative somata per cells with GFP-positive somata in *Scn1a*-GFP mouse neocortex and hippocampus. Values for Figure 8B-left panels. ^a^L2/3, L5, L6, CA1, CA2/3, DG: neocortical layer II/III, V, VI, cornu ammonis 1, 2 plus 3, dentate gyrus. Cells in three *Scn1a*-GFP mice (line #233) at 4W were counted; ^b^L2/3 (N = 631 cells), L5 (N = 533 cells), L6 (N = 390 cells), CA1 (N = 231 cells), CA2/3 (N = 311 cells) and DG (N = 258 cells). Values are presented as mean ± SEM. +, positive; -, negative.

| Region <sup>a</sup> | per cells with GFP+ somata |  |  |  |
| --- | --- | --- | --- | --- |
|  | % PV+, SST+ <sup>b</sup> | % PV+ <sup>b</sup> | % SST+ <sup>b</sup> | % PV-, SST- <sup>b</sup> |
| L2/3 | 0.1 ± 0.1 | 14.7 ± 1.4 | 6.4 ± 2.5 | 78.8 ± 3.8 |
| L5 | 0.1 ± 0.1 | 26.3 ± 1.5 | 11.1 ± 2.1 | 62.4 ± 3.7 |
| L6 | 0.4 ± 0.4 | 21.8 ± 2.2 | 14.8 ± 1.6 | 63.0 ± 3.7 |
| CA1 | 2.0 ± 0.3 | 33.6 ± 3.2 | 22.7 ± 3.5 | 41.7 ± 7.4 |
| CA2/3 | 1.4 ± 0.3 | 24.6 ± 5.1 | 16.0 ± 1.5 | 58.1 ± 6.5 |
| DG | 0.0 ± 0.0 | 19.3 ± 4.5 | 21.4 ± 1.4 | 59.3 ± 3.4 |

**Supplementary table S12.** Percentage of cells with GFP-positive/negative somata per cells with PV-positive somata. Values for Figure 8B-middle panels. ^a^L2/3, L5, L6, CA1, CA2/3, DG: neocortical layer II/III, V, VI, cornu ammonis 1, 2 plus 3, dentate gyrus. Cells in three *Scn1a*-GFP mice (line #233) at 4W were counted; ^b^L2/3 (N = 95 cells), L5 (N = 144 cells), L6 (N = 84 cells), CA1 (N = 83 cells), CA2/3 (N = 78 cells) and DG (N = 37 cells). Values are presented as mean ± SEM. +, positive; -, negative.

| Region <sup>a</sup> | per cells with PV+ somata |  |
| --- | --- | --- |
|  | %GFP+ <sup>b</sup> | % GFP- <sup>b</sup> |
| L2/3 | 100.0 ± 0.0 | 0.0 ± 0.0 |
| L5 | 98.4 ± 1.5 | 1.6 ± 1.5 |
| L6 | 100.0 ± 0.0 | 0.0 ± 0.0 |
| CA1 | 100.0 ± 0.0 | 0.0 ± 0.0 |
| CA2/3 | 100.0 ± 0.0 | 0.0 ± 0.0 |
| DG | 100.0 ± 0.0 | 0.0 ± 0.0 |

**Supplementary table S13.** Percentage of cells with GFP-positive/negative somata per cells with SST-positive somata. Values for Figure 8B-right panels. ^a^L2/3, L5, L6, CA1, CA2/3, DG: neocortical layer II/III, V, VI, cornu ammonis 1, 2 plus 3, dentate gyrus. Cells in three *Scn1a*-GFP mice (line #233) at 4W were counted; ^b^L2/3 (N = 46 cells), L5 (N = 76 cells), L6 (N = 58 cells), CA1 (N = 58 cells), CA2/3 (N = 56 cells) and DG (N = 55 cells). Values are presented as mean ± SEM. +, positive; -, negative.

| Region <sup>a</sup> | per cells with SST+ somata |  |
| --- | --- | --- |
|  | %GFP+ <sup>b</sup> | % GFP- <sup>b</sup> |
| L2/3 | 89.8 ± 7.6 | 10.2 ± 7.6 |
| L5 | 83.3 ± 4.1 | 16.7 ± 4.1 |
| L6 | 95.4 ± 2.8 | 4.6 ± 2.8 |
| CA1 | 100.0 ± 0.0 | 0.0 ± 0.0 |
| CA2/3 | 96.4 ± 1.9 | 3.4 ± 1.9 |
| DG | 99.3 ± 0.7 | 0.7 ± 0.7 |

**Supplementary table S14.**
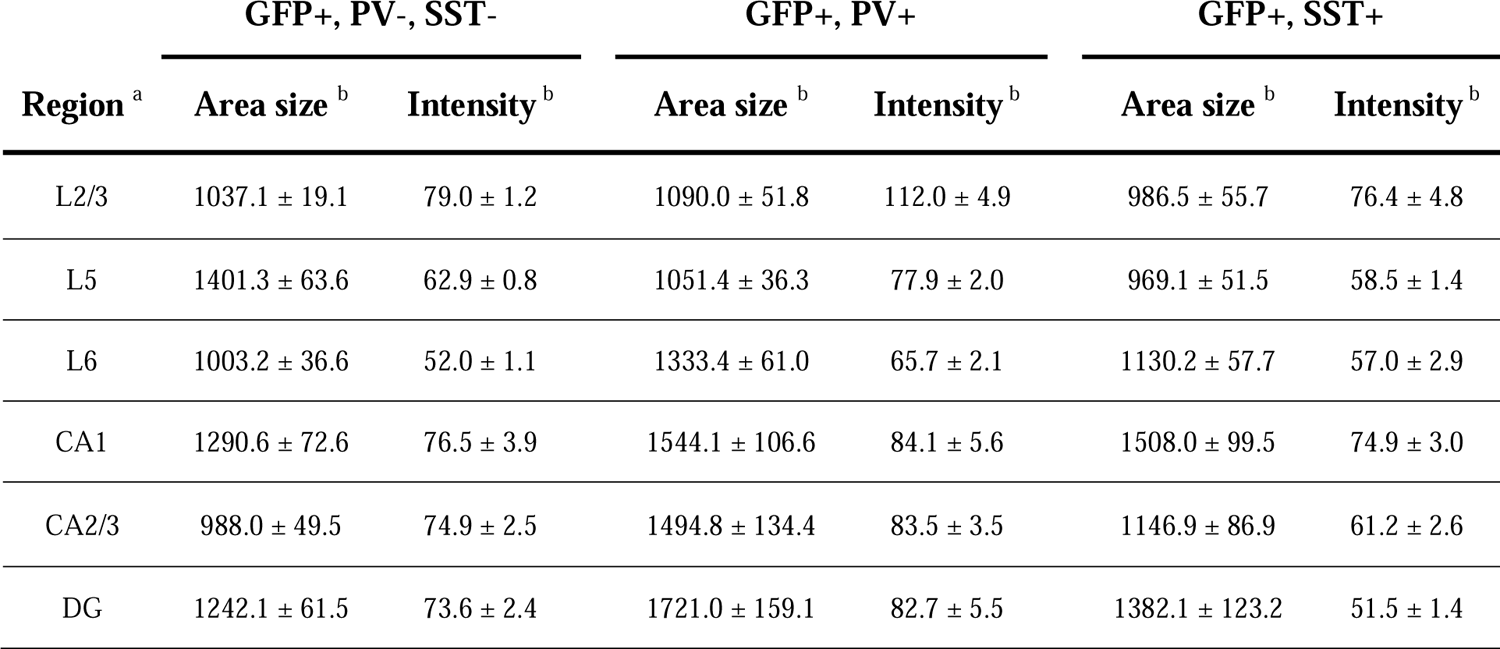
Area size and intensity of GFP immunosignals in GFP-positive cells with PV/SST-positive or negative somata in *Scn1a*-GFP mouse neocortex and hippocampus. Values for Figure 9A. ^a^L2/3, L5, L6, CA1, CA2/3, DG: neocortical layer II/III, V, VI, cornu ammonis 1, 2 plus 3, dentate gyrus. Cells in two *Scn1a*-GFP mice (line #233) at 4W were counted; ^b^L2/3 (N = 157 cells), L5 (N = 122 cells) and L6 (N = 109 cells), CA1 (N = 52 cells), CA2/3 (N = 85 cells) and DG (N = 60 cells). Values are presented as mean ± SEM. +, positive; -, negative.

| Region <sup>a</sup> | GFP+, PV-, SST- |  | GFP+, PV+ |  | GFP+, SST+ |  |
| --- | --- | --- | --- | --- | --- | --- |
|  | Area size <sup>b</sup> | Intensity <sup>b</sup> | Area size <sup>b</sup> | Intensity <sup>b</sup> | Area size <sup>b</sup> | Intensity <sup>b</sup> |
| L2/3 | 1037.1 ± 19.1 | 79.0 ± 1.2 | 1090.0 ± 51.8 | 112.0 ± 4.9 | 986.5 ± 55.7 | 76.4 ± 4.8 |
| L5 | 1401.3 ± 63.6 | 62.9 ± 0.8 | 1051.4 ± 36.3 | 77.9 ± 2.0 | 969.1 ± 51.5 | 58.5 ± 1.4 |
| L6 | 1003.2 ± 36.6 | 52.0 ± 1.1 | 1333.4 ± 61.0 | 65.7 ± 2.1 | 1130.2 ± 57.7 | 57.0 ± 2.9 |
| CA1 | 1290.6 ± 72.6 | 76.5 ± 3.9 | 1544.1 ± 106.6 | 84.1 ± 5.6 | 1508.0 ± 99.5 | 74.9 ± 3.0 |
| CA2/3 | 988.0 ± 49.5 | 74.9 ± 2.5 | 1494.8 ± 134.4 | 83.5 ± 3.5 | 1146.9 ± 86.9 | 61.2 ± 2.6 |
| DG | 1242.1 ± 61.5 | 73.6 ± 2.4 | 1721.0 ± 159.1 | 82.7 ± 5.5 | 1382.1 ± 123.2 | 51.5 ± 1.4 |

**Supplementary table S15.**
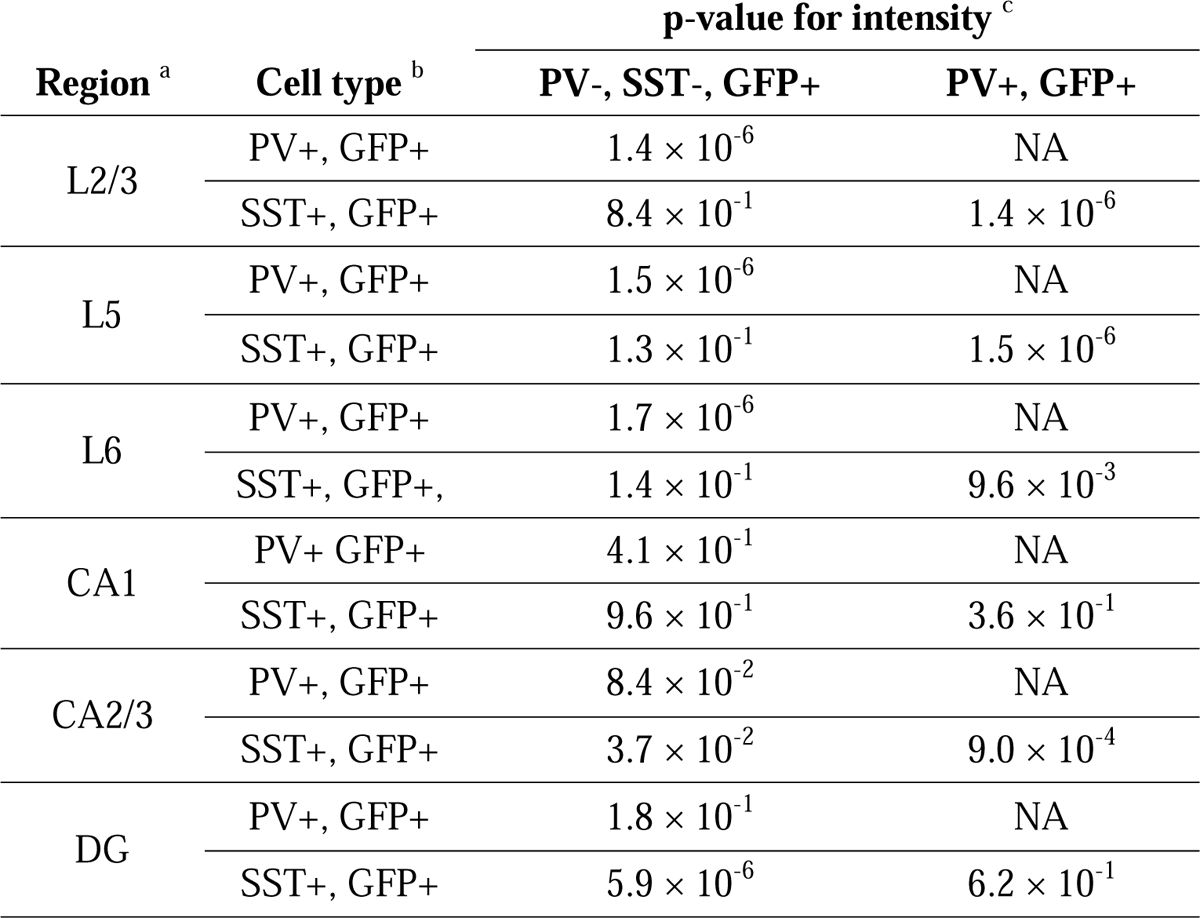
Statistical significance for intensity of GFP immunosignals in GFP-positive cells with PV/SST-positive or negative somata. Values for Figure 9B-left panels. ^a^L2/3, L5, L6, CA1, CA2/3, DG: neocortical layer II/III, V, VI, cornu ammonis 1, 2 plus 3, dentate gyrus. Cells in two *Scn1a*-GFP mice (line #233) at 4W were counted; ^b^L2/3 (N = 157 cells), L5 (N = 122 cells) and L6 (N = 109 cells), CA1 (N = 52 cells), CA2/3 (N = 85 cells) and DG (N = 60 cells). ^c^Statistical significance was assessed using one-way ANOVA followed by Tukey–Kramer post-hoc multiple comparison test. Values are presented as mean ± SEM. +, positive; -, negative; NA, not available.

**Supplementary table S16.** Statistical significance for area size of GFP immunosignals in GFP-positive cells with PV/SST-positive or negative somata. Values for Figure 9B-right panels. ^a^L2/3, L5, L6, CA1, CA2/3, DG: neocortical layer II/III, V, VI, cornu ammonis 1, 2 plus 3, dentate gyrus. Cells in two *Scn1a*-GFP mice (line #233) at 4W were counted; ^b^L2/3 (N = 157 cells), L5 (N = 122 cells) and L6 (N = 109 cells), CA1 (N = 52 cells), CA2/3 (N = 85 cells) and DG (N = 60 cells). ^c^Statistical significance was assessed using one-way ANOVA followed by Tukey–Kramer post-hoc multiple comparison test. Values are presented as mean ± SEM. +, positive; -, negative; NA, not available.

| Region <sup>a</sup> | Cell type <sup>b</sup> | p-value for area size <sup>c</sup> |  |
| --- | --- | --- | --- |
|  |  | PV-, SST-, GFP+ | PV+, GFP+ |
| L2/3 | PV+, GFP+ | $4.9 \times 10^{-1}$ | NA |
| | SST+, GFP+ | $7.2 \times 10^{-1}$ | $3.6 \times 10^{-1}$ |
| L5 | PV+, GFP+ | $1.2 \times 10^{-4}$ | NA |
| | SST+, GFP+ | $1.5 \times 10^{-4}$ | $7.3 \times 10^{-1}$ |
| L6 | PV+, GFP+ | $8.0 \times 10^{-6}$ | NA |
| | SST+, GFP+ | $1.9 \times 10^{-1}$ | $3.8 \times 10^{-2}$ |
| CA1 | PV+, GFP+ | $1.2 \times 10^{-1}$ | NA |
| | SST+, GFP+ | $2.0 \times 10^{-1}$ | $9.6 \times 10^{-1}$ |
| CA2/3 | PV+, GFP+ | $8.2 \times 10^{-5}$ | NA |
| | SST+, GFP+ | $5.6 \times 10^{-1}$ | $9.8 \times 10^{-2}$ |
| DG | PV+, GFP+ | $9.7 \times 10^{-3}$ | NA |
| | SST+, GFP+ | $5.1 \times 10^{-1}$ | $1.5 \times 10^{-1}$ |

**Supplementary table S17.** Percentage of cells with Nav1.1- and Nav1.2-positive/negative cells per cells with ankyrinG-positive AISs in *Scn1a*-GFP mouse neocortex. Values for Figure 10B. ^a^L2/3, L5, L6: neocortical layer II/III, V, VI. Cells in three *Scn1a*-GFP mice (line #233) at P15 were counted; ^b^L2/3 (N = 1300 cells), L5 (N = 895 cells), and L6 (N = 1295 cells). Values are presented as mean ± SEM. AnkG, ankyrinG; +, positive; -, negative.

| Region <sup>a</sup> | per cells with AnkG+ AISs |  |  |  |
| --- | --- | --- | --- | --- |
|  | % Nav1.1+ <sup>b</sup> |  | % Nav1.1- <sup>b</sup> |  |
| L2/3 | 4.7 ± 0.1 |  | 95.3 ± 0.1 |  |
| L5 | 6.0 ± 1.5 |  | 94.0 ± 1.5 |  |
| L6 | 2.5 ± 0.3 |  | 97.5 ± 0.3 |  |
|  | % Nav1.2+ | % Nav1.2- | % Nav1.2+ | % Nav1.2- |
| L2/3 | 0.3 ± 0.2 | 4.4 ± 0.1 | 78.1 ± 3.6 | 17.2 ± 3.7 |
| L5 | 0.1 ± 0.1 | 5.9 ± 1.4 | 69.3 ± 4.5 | 24.7 ± 5.7 |
| L6 | 0.1 ± 0.1 | 2.3 ± 0.1 | 69.3 ± 3.4 | 28.3 ± 3.2 |

**Supplementary table S18.**
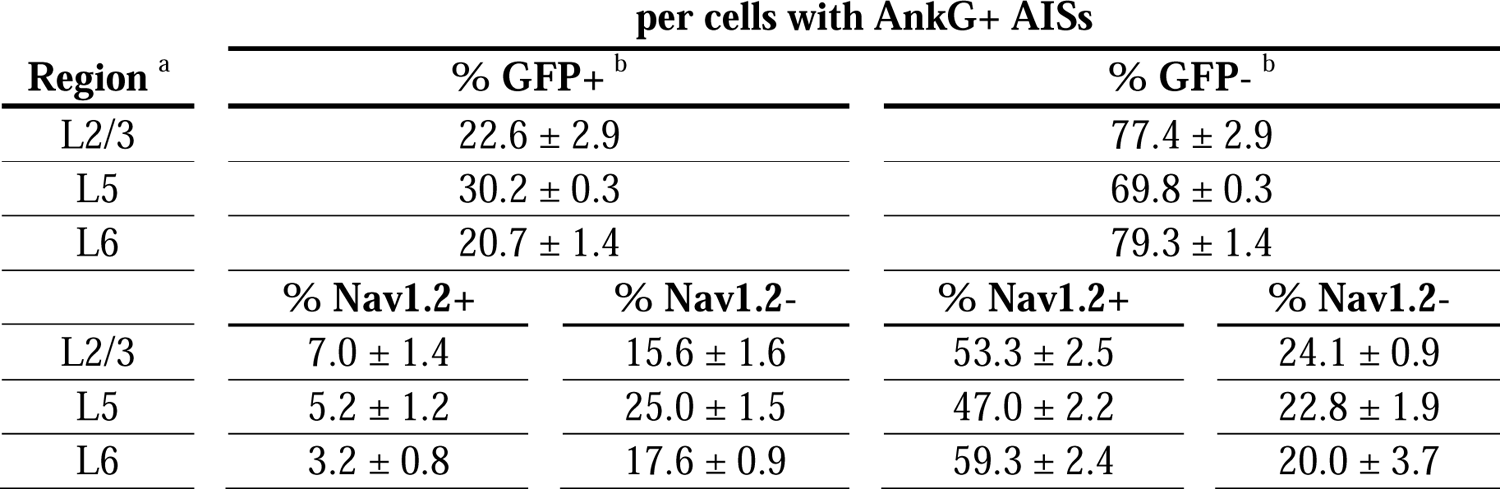
Percentage of cells with GFP- and Nav1.2-positive/negative somata and AISs per cells with ankyrinG-positive AISs in *Scn1a*-GFP mouse neocortex. Values for Figure 11B-left panel. ^a^L2/3, L5, L6: neocortical layer II/III, V, VI. Cells in three *Scn1a*-GFP mice (line #233) at P15 were counted; ^b^L2/3 (N = 877 cells), L5 (N = 724 cells) and L6 (N = 682 cells). Values are presented as mean ± SEM. AnkG, ankyrinG; +, positive; -, negative.

| Region <sup>a</sup> | per cells with AnkG+ AISs |  |  |  |
| --- | --- | --- | --- | --- |
|  | % GFP+ <sup>b</sup> |  | % GFP- <sup>b</sup> |  |
| L2/3 | 22.6 ± 2.9 |  | 77.4 ± 2.9 |  |
| L5 | 30.2 ± 0.3 |  | 69.8 ± 0.3 |  |
| L6 | 20.7 ± 1.4 |  | 79.3 ± 1.4 |  |
|  | % Nav1.2+ | % Nav1.2- | % Nav1.2+ | % Nav1.2- |
| L2/3 | 7.0 ± 1.4 | 15.6 ± 1.6 | 53.3 ± 2.5 | 24.1 ± 0.9 |
| L5 | 5.2 ± 1.2 | 25.0 ± 1.5 | 47.0 ± 2.2 | 22.8 ± 1.9 |
| L6 | 3.2 ± 0.8 | 17.6 ± 0.9 | 59.3 ± 2.4 | 20.0 ± 3.7 |

**Supplementary table S19.**
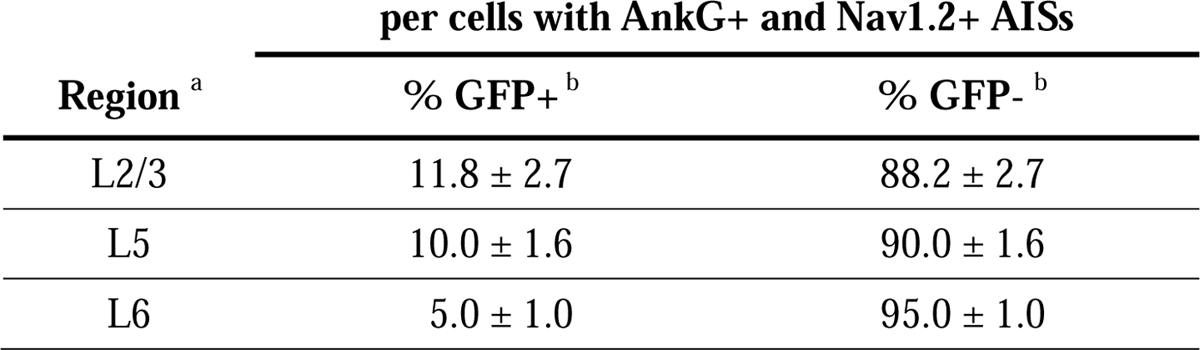
Percentage of cells with GFP-positive/negative somata per cells with Nav1.2/ankyrinG-double positive AISs in *Scn1a*-GFP mouse neocortex. Values for Figure 11B-middle panel. ^a^L2/3, L5, L6: neocortical layer II/III, V, VI. Cells in three *Scn1a*-GFP mice (line #233) at P15 were counted; ^b^L2/3 (N = 527 cells), L5 (N = 378 cells) and L6 (N = 422 cells). Values are presented as mean ± SEM. AnkG, ankyrinG; +, positive; -, negative.

| Region <sup>a</sup> | per cells with AnkG+ and Nav1.2+ AISs |  |
| --- | --- | --- |
|  | % GFP+ <sup>b</sup> | % GFP- <sup>b</sup> |
| L2/3 | 11.8 ± 2.7 | 88.2 ± 2.7 |
| L5 | 10.0 ± 1.6 | 90.0 ± 1.6 |
| L6 | 5.0 ± 1.0 | 95.0 ± 1.0 |

**Supplementary table S20.** Percentage of cells with Nav1.2-positive/negative AISs per cells with ankyrinG-positive AISs and GFP+ somata in *Scn1a*-GFP mouse neocortex. Values for Figure 11B-right panel. ^a^L2/3, L5, L6: neocortical layer II/III, V, VI. Cells in three *Scn1a*-GFP mice (line #233) at P15 were counted; ^b^L2/3 (N = 197 cells), L5 (N = 215 cells) and L6 (N = 139 cells). Values are presented as mean ± SEM. AnkG, ankyrinG; +, positive; -, negative.

| Region <sup>a</sup> | per cells with AnkG+ AISs and GFP+ somata |  |
| --- | --- | --- |
|  | % Nav1.2+ <sup>b</sup> | % Nav1.2- <sup>b</sup> |
| L2/3 | 31.5 ± 2.3 | 68.5 ± 2.3 |
| L5 | 17.1 ± 4.4 | 82.9 ± 4.4 |
| L6 | 14.4 ± 2.6 | 85.6 ± 2.6 |

**Supplementary table S21.**
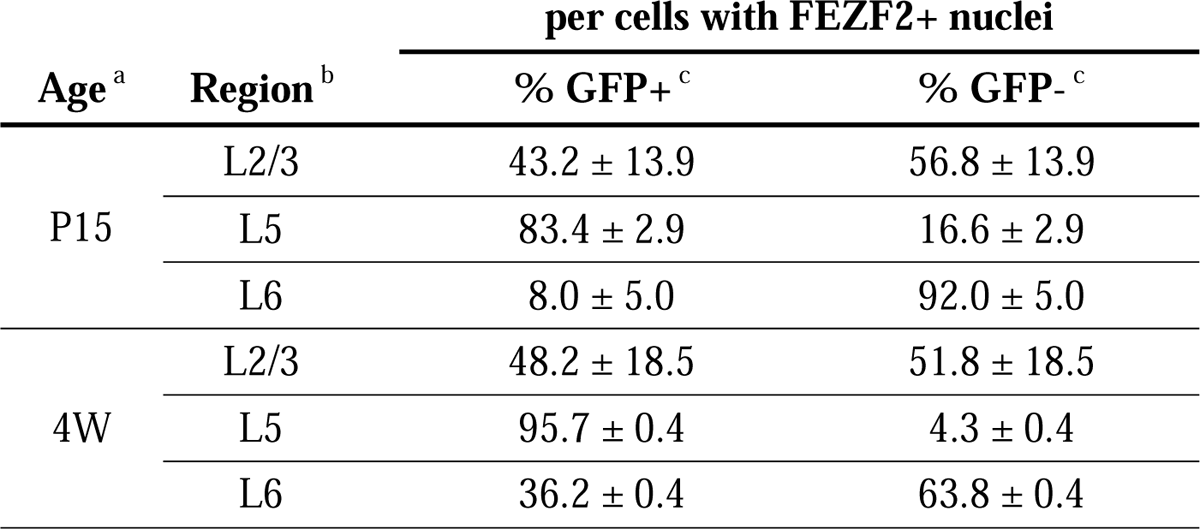
Percentage of cells with FEZF2- and GFP-positive/negative nuclei and somata per cells with GFP-positive somata and/or FEZF2-positive nuclei in *Scn1a*-GFP mouse neocortex. Values for Figure 12B-left panels. ^a^Two animals were counted for each age. ^b^L2/3, L5, L6: neocortical layer II/III, V, VI. Cells in *Scn1a*-GFP mice (line #233) were counted; ^c^L2/3 (P15, N = 670 cells; 4W, N = 325 cells), L5 (P15, N = 733 cells; 4W, N = 431 cells) and L6 (P15, N = 466 cells; 4W, N = 386 cells). Values are presented as mean ± SEM. +, positive; -, negative.

**Supplementary table S22.**
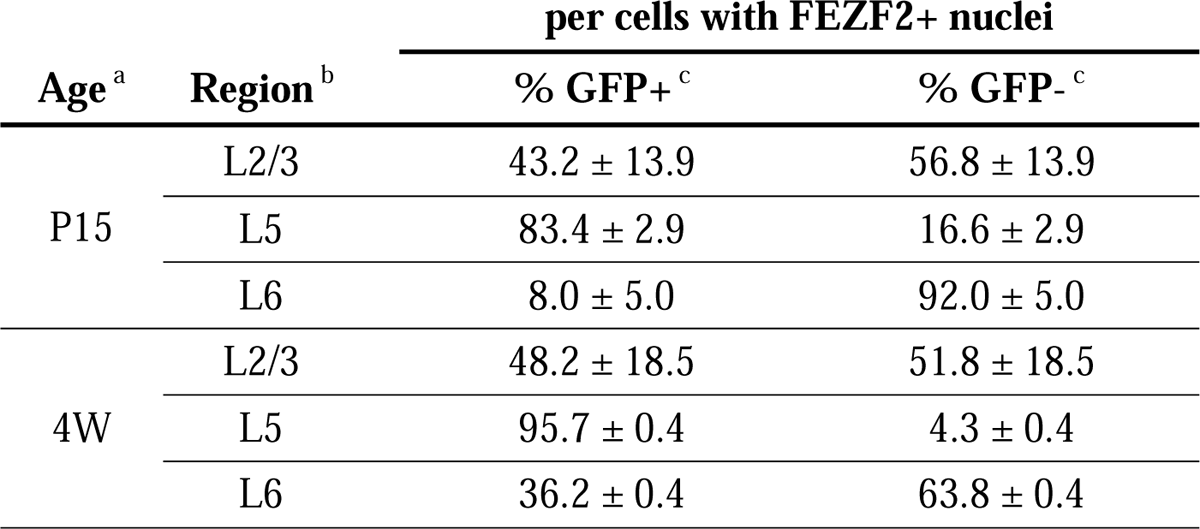
Percentage of cells with GFP-positive/negative somata per cells with FEZF2-positive nuclei in *Scn1a*-GFP mouse neocortex. Values for Figure 12B-middle panels. ^a^Two animals were counted for each age. ^b^L2/3, L5, L6: neocortical layer II/III, V, VI. Cells in *Scn1a*-GFP mice (line #233) were counted; ^c^L2/3 (P15, N = 198 cells; 4W, N = 25 cells), L5 (P15, N = 431 cells; 4W, N = 214 cells) and L6 (P15, N = 283 cells; 4W, N = 202 cells). Values are presented as mean ± SEM. +, positive; -, negative.

**Supplementary table S23.**
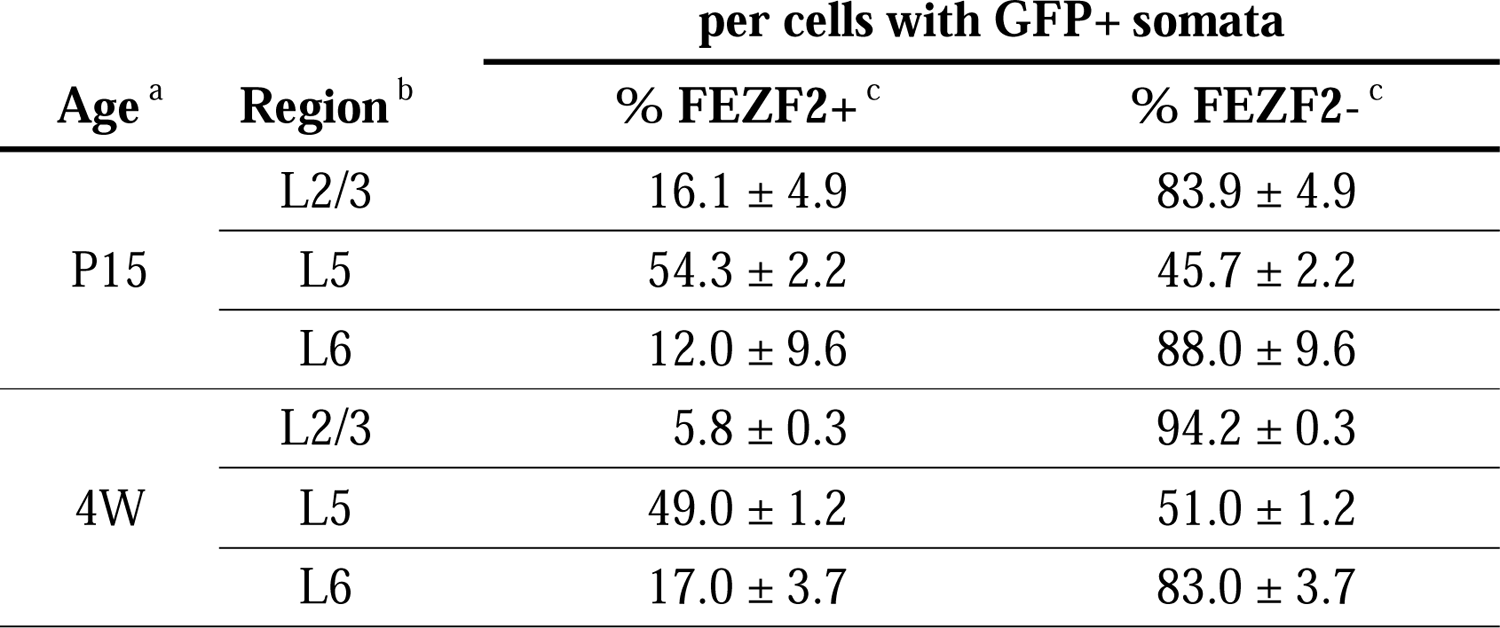
Percentage of cells with FEZF2-positive/negative nuclei per cells with GFP-positive somata in *Scn1a*-GFP mouse neocortex. Values for Figure 12B-right panels. ^a^Two animals were counted for each age. ^b^L2/3, L5, L6: neocortical layer II/III, V, VI. Cells in *Scn1a*-GFP mice (line #233) at P15 were counted; ^c^L2/3 (P15, N = 567 cells; 4W, N = 318 cells), L5 (P15, N = 479 cells; 4W, N = 363 cells) and L6 (P15, N = 202 cells; 4W, N = 228 cells). Values are presented as mean ± SEM. +, positive; -, negative.

**Supplementary table S24.** Area size and intensity of GFP immunosignals in GFP-positive cells with FEZF2-positive or negative nuclei in *Scn1a*-GFP mouse neocortex. Values for Figure 13. ^a^L2/3, L5, L6: neocortical layer II/III, V, VI. Cells in one *Scn1a*-GFP mouse (line #233) at 4W were counted; ^b^L2/3 (N = 159 cells), L5 (N = 147 cells) and L6 (N = 80 cells). ^c^Statistical significance was assessed using t-test. Values are presented as mean ± SEM. +, positive; -, negative.

| Region<sup>a</sup> | FEZF2+, GFP+ |  | FEZF2-, GFP+ |  | p-value |  |
| --- | --- | --- | --- | --- | --- | --- |
|  | Area size<sup>b</sup> | Intensity<sup>b</sup> | Area size<sup>b</sup> | Intensity<sup>b</sup> | Area size<sup>c</sup> | Intensity<sup>c</sup> |
| L2/3 | 510.8 ± 9.0 | 50.4 ± 0.8 | 377.3 ± 9.0 | 48.0 ± 0.8 | 1.0 × 10<sup>-2</sup> | 5.8 × 10<sup>-1</sup> |
| L5 | 577.4 ± 21.4 | 47.6 ± 1.6 | 461.6 ± 18.1 | 25.9 ± 3.1 | 7.3 × 10<sup>-5</sup> | 3.8 × 10<sup>-8</sup> |
| L6 | 389.3 ± 27.9 | 43.0 ± 3.2 | 372.1 ± 19.0 | 52.4 ± 2.3 | 6.4 × 10<sup>-1</sup> | 3.8 × 10<sup>-2</sup> |

**Supplementary table S25.** Percentage of cells with FEZF2- and Nav1.1-positive/negative nuclei or AISs per cells with ankyrinG-positive AISs in *Scn1a*-GFP mouse neocortex. Values for Supplementary figure S8B-left panel. ^a^L5, L6: neocortical layer II/III, V, VI. Cells in three *Scn1a*-GFP mice (line #233) at P15 were counted; ^b^L5 (N = 588 cells) and L6 (N = 580 cells). Values are presented as mean ± SEM. AnkG, ankyrinG; +, positive; -, negative.

| Region <sup>a</sup> | per cells with AnkG+ AISs |  |  |  |
| --- | --- | --- | --- | --- |
|  | % FEZF2+ <sup>b</sup> |  | % FEZF2- <sup>b</sup> |  |
| L5 | 52.8 ± 0.7 |  | 47.2 ± 0.7 |  |
| L6 | 66.3 ± 5.5 |  | 33.7 ± 5.5 |  |
|  | % Nav1.1+ | % Nav1.1- | % Nav1.1+ | % Nav1.1- |
| L5 | 2.2 ± 0.4 | 18.5 ± 0.8 | 4.9 ± 1.6 | 74.4 ± 1.3 |
| L6 | 0.4 ± 0.3 | 18.7 ± 1.4 | 2.3 ± 0.3 | 78.6 ± 1.7 |

**Supplementary table S26.** Percentage of cells with FEZF2-positive/negative nuclei per cells with ankyrinG/Nav1.1-double positive AISs in *Scn1a*-GFP mouse neocortex. Values for Supplementary figure S8B-middle panel. ^a^L5, L6: neocortical layer V, VI. Cells in three *Scn1a*-GFP mice (line #233) at P15 were counted; ^b^L5 (N = 115 cells) and L6 (N = 45 cells). Values are presented as mean ± SEM. AnkG, ankyrinG; +, positive; -, negative.

| Region <sup>a</sup> | per cells with AnkG+ and Nav1.1+ AISs |  |
| --- | --- | --- |
|  | % FEZF2+ <sup>b</sup> | % FEZF2- <sup>b</sup> |
| L5 | 35.9 ± 9.7 | 64.1 ± 9.7 |
| L6 | 12.4 ± 8.6 | 87.6 ± 8.6 |

**Supplementary table S27.** Percentage of cells with Nav1.1-positive/negative AISs per cells with ankyrinG-positive AISs and FEZF2-positive nuclei in *Scn1a*-GFP mouse neocortex. Values for Supplementary figure S8B-right panel. ^a^L5, L6: neocortical layer V, VI. Cells in three *Scn1a*-GFP mice (line #233) at P15 were counted; ^b^L5 (N = 326 cells) and L6 (N = 324 cells). Values are presented as mean ± SEM. AnkG, ankyrinG; +, positive; -, negative.

| Region <sup>a</sup> | % Nav1.1+ <sup>b</sup> | % Nav1.1- <sup>b</sup> |
| --- | --- | --- |
| L5 | 11.1 ± 1.9 | 88.9 ± 1.9 |
| L6 | 1.7 ± 1.3 | 98.3 ± 1.3 |

**Supplementary table S28.**
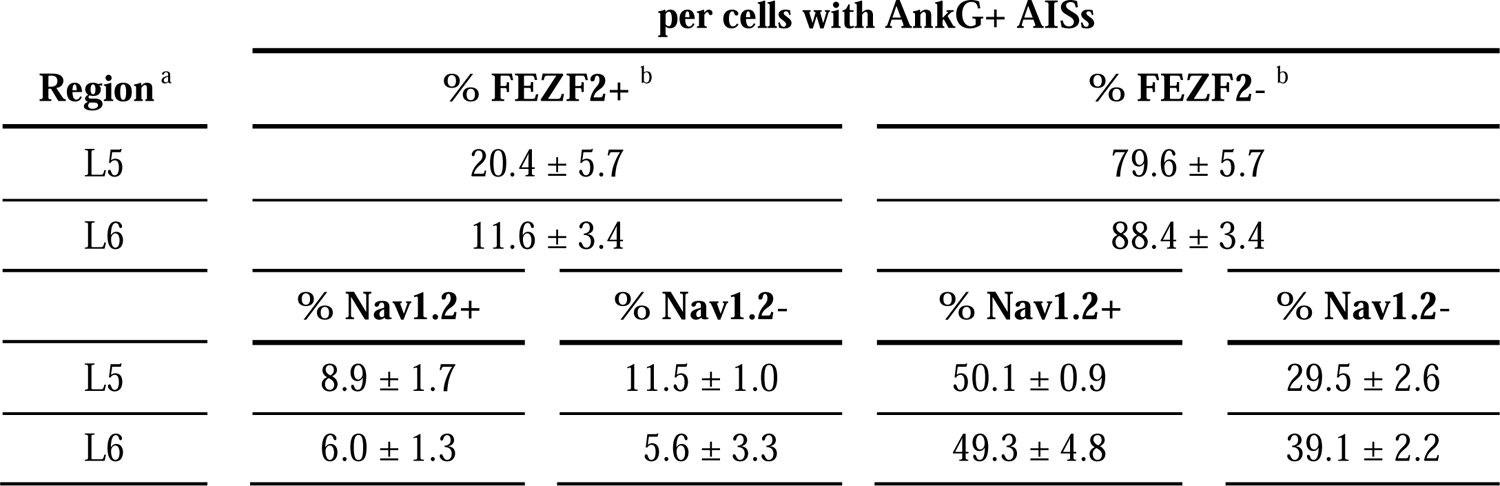
Percentage of cells with FEZF2- and Nav1.2-positive/negative nuclei or AISs per cells with ankyrinG-positive AISs in *Scn1a*-GFP mouse neocortex. Values for Supplementary figure S9B-left-upper panel. ^a^L5, L6: neocortical layer V, VI. Cells in three *Scn1a*-GFP mice (line #233) at P15 were counted; ^b^L5 (N = 644 cells) and L6 (N = 758 cells). Values are presented as mean ± SEM. AnkG, ankyrinG; +, positive; -, negative.

**Supplementary table S29.** Percentage of cells with FEZF2-positive/negative nuclei per cells with ankyrinG/Nav1.2-double positive AISs in *Scn1a*-GFP mouse neocortex. Values for Supplementary figure S9B-middle-upper panel. ^a^L5, L6: neocortical layer V, VI. Cells in three *Scn1a*-GFP mice (line #233) at P15 were counted; ^b^L5 (N = 376 cells) and L6 (N = 415 cells). Values are presented as mean ± SEM. AnkG, ankyrinG; +, positive; -, negative.

| Region <sup>a</sup> | per cells with AnkG+ and Nav1.2+ AISs |  |
| --- | --- | --- |
|  | % FEZF2+ <sup>b</sup> | % FEZF2- <sup>b</sup> |
| L5 | 15.1 ± 2.9 | 84.9 ± 2.9 |
| L6 | 10.5 ± 4.3 | 89.5 ± 4.3 |

**Supplementary table S30.** Percentage of cells with Nav1.2-positive/negative AISs per cells with ankyrinG-positive AISs and FEZF2-positive nuclei in *Scn1a*-GFP mouse neocortex. Values for Supplementary figure S9B-right-upper panel. ^a^L5, L6: neocortical layer V, VI. Cells in three *Scn1a*-GFP mice (line #233) at P15 were counted; ^b^L5 (N = 135 cells) and L6 (N = 94 cells). Values are presented as mean ± SEM. AnkG, ankyrinG; +, positive; -, negative.

| Region <sup>a</sup> | per cells with AnkG+ AISs and FEZF2+ nuclei |  |
| --- | --- | --- |
|  | % Nav1.2+ <sup>b</sup> | % Nav1.2- <sup>b</sup> |
| L5 | 51.2 ± 3.7 | 48.8 ± 3.7 |
| L6 | 41.6 ± 9.5 | 58.4 ± 9.5 |

**Supplementary table S31.** Percentage of cells with ankyrinG-positive/negative AISs per cells with FEZF2-positive nuclei in *Scn1a*-GFP mouse neocortex. Values for Supplementary figure S9B-lower panel. ^a^L5, L6: neocortical layer V, VI. Cells in three *Scn1a*-GFP mice (line #233) at P15 were counted; ^b^L5 (N = 301 cells), and L6 (N = 257 cells). Values are presented as mean ± SEM. AnkG, ankyrinG; +, positive; -, negative.

| Region <sup>a</sup> | per cells with FEZF2+ nuclei |  |
| --- | --- | --- |
|  | % AnkG+ <sup>b</sup> | % AnkG- <sup>b</sup> |
| L5 | 38.6 ± 7.1 | 61.4 ± 7.1 |
| L6 | 39.4 ± 6.3 | 60.6 ± 6.3 |

**Supplementary table S32.** Percentage of cells with GFP- and Nav1.2-positive/negative somata and AISs per cells with FEZF2-positive nuclei in *Scn1a*-GFP mouse neocortex. Values for Supplementary figure S10B. ^a^L5, L6: neocortical layer V, VI. Cells in three *Scn1a*-GFP mice (line #233) at P15 were counted; ^b^L5 (N = 267 cells) and L6 (N = 188 cells). ^c^In these values, because they were including ankyrinG-negative cells, to obtain correct cell population for Nav1.2-negative/ankyrinG-positive cells, virtual cell number were estimated using ratio of ankyrinG/FEZF2 double positive cells in Supplementary figure S9B.Values are presented as mean ± SEM. +, positive; -, negative.

| Region <sup>a</sup> | per cells with FEZF2+ nuclei |  |  |  |
| --- | --- | --- | --- | --- |
|  | % GFP+ <sup>b</sup> |  | % GFP- <sup>b</sup> |  |
| L5 | 74.4 ± 3.4 |  | 25.6 ± 3.4 |  |
| L6 | 16.5 ± 4.7 |  | 83.5 ± 4.7 |  |
|  | % Nav1.2+ <sup>c</sup> | % Nav1.2- <sup>c</sup> | % Nav1.2+ <sup>c</sup> | % Nav1.2- <sup>c</sup> |
| L5 | 13.9 ± 2.7 | 56.4 ± 6.8 | 13.2 ± 5.7 | 16.5 ± 3.2 |
| L6 | 5.3 ± 3.7 | 9.6 ± 3.3 | 34.7 ± 3.0 | 50.3 ± 8.7 |

**Supplementary table S33.** Percentage of cells with TBR1- and GFP-positive/negative nuclei and somata per GFP-positive cells and/or TBR1-positive nuclei in *Scn1a*-GFP mouse neocortex. Values for Figure 14B-left panels. ^a^Two animals were counted for each age. ^b^L2/3, L5, L6: neocortical layer II/III, V, VI. Cells in *Scn1a*-GFP mice (line #233) were counted; ^c^L2/3 (P15, N = 704 cells; 4W, N = 549 cells), L5 (P15, N = 639 cells; 4W, N = 400 cells) and L6 (P15, N = 990 cells; 4W, N = 598 cells). Values are presented as mean ± SEM. +, positive; -, negative.

| Age<sup>a</sup> | Region<sup>b</sup> | per cells with GFP+ somata and/or TBR1+ nuclei |  |  |
| --- | --- | --- | --- | --- |
|  |  | % GFP+, TBR1-<sup>c</sup> | % GFP+, TBR1+<sup>c</sup> | % GFP-, TBR1+<sup>c</sup> |
| P15 | L2/3 | 46.5 ± 0.8 | 38.7 ± 0.4 | 14.8 ± 0.3 |
|  | L5 | 49.3 ± 1.2 | 5.5 ± 2.0 | 45.1 ± 0.8 |
|  | L6 | 14.8 ± 2.3 | 12.9 ± 7.4 | 72.3 ± 9.8 |
| 4W | L2/3 | 35.3 ± 0.2 | 42.1 ± 1.5 | 22.6 ± 1.3 |
|  | L5 | 54.7 ± 0.1 | 2.4 ± 1.8 | 42.9 ± 0.1 |
|  | L6 | 13.3 ± 1.8 | 22.1 ± 4.5 | 64.6 ± 6.3 |

**Supplementary table S34.** Percentage of cells with GFP-positive/negative somata per cells with TBR1-positive nuclei in *Scn1a*-GFP mouse neocortex. Values for Figure 14B-middle panels. ^a^Two animals were counted for each age. ^b^L2/3, L5, L6: neocortical layer II/III, V, VI. Cells in *Scn1a*-GFP mice (line #233) were counted; ^c^L2/3 (P15, N = 367 cells; 4W, N = 358 cells), L5 (P15, N = 322 cells; 4W, N = 180 cells) and L6 (P15, N = 848 cells; 4W, N = 518 cells). Values are presented as mean ± SEM. +, positive; -, negative.

| Age<sup>a</sup> | Region<sup>b</sup> | per cells with TBR1+ nuclei |  |
| --- | --- | --- | --- |
|  |  | % GFP+<sup>c</sup> | % GFP-<sup>c</sup> |
| P15 | L2/3 | 71.9 ± 0.4 | 28.1 ± 0.4 |
|  | L5 | 10.7 ± 3.9 | 89.3 ± 3.9 |
|  | L6 | 15.4 ± 9.1 | 84.6 ± 9.1 |
| 4W | L2/3 | 64.7 ± 1.6 | 35.3 ± 1.6 |
|  | L5 | 5.1 ± 3.9 | 94.9 ± 3.9 |
|  | L6 | 25.6 ± 5.7 | 74.4 ± 5.7 |

**Supplementary table S35.** Percentage of cells with TBR1-positive/negative nuclei per cells with GFP-positive somata in *Scn1a*-GFP mouse neocortex. Values for Figure 14B-right panels. ^a^Two animals were counted for each age. ^b^L2/3, L5, L6: neocortical layer II/III, V, VI. Cells in *Scn1a*-GFP mice (line #233) were counted; ^c^L2/3 (P15, N = 601 cells; 4W, N = 422 cells), L5 (P15, N = 350 cells; 4W, N = 228 cells) and L6 (P15, N = 235 cells; 4W, N = 222 cells). Values are presented as mean ± SEM. +, positive; -, negative.

| Age<sup>a</sup> | Region<sup>b</sup> | per cells with GFP+ somata |  |
| --- | --- | --- | --- |
|  |  | % TBR1+<sup>c</sup> | % TBR1-<sup>c</sup> |
| P15 | L2/3 | 45.1 ± 0.7 | 54.9 ± 0.7 |
|  | L5 | 10.0 ± 3.5 | 90.0 ± 3.5 |
|  | L6 | 41.5 ± 12.9 | 58.5 ± 12.9 |
| 4W | L2/3 | 54.4 ± 0.7 | 45.6 ± 0.7 |
|  | L5 | 4.1 ± 3.2 | 95.9 ± 3.2 |
|  | L6 | 61.2 ± 2.9 | 38.8 ± 2.9 |

**Supplementary table S36.** Area size and intensity of GFP immunosignals in GFP-positive cells with TBR1-positive or negative nuclei in *Scn1a*-GFP mouse neocortex. Values for Figure 15. ^a^L2/3, L5, L6: neocortical layer II/III, V, VI. Cells in one *Scn1a*-GFP mouse (line #233) at 4W were counted; ^b^L2/3 (N = 189 cells), L5 (N = 118 cells) and L6 (N = 105 cells). ^c^Statistical significance was assessed using t-test. Values are presented as mean ± SEM. +, positive; -, negative.

| Region<sup>a</sup> | TBR1+, GFP+ |  | TBR1-, GFP+ |  | p-value |  |
| --- | --- | --- | --- | --- | --- | --- |
|  | Area size<sup>b</sup> | Intensity<sup>b</sup> | Area size<sup>b</sup> | Intensity<sup>b</sup> | Area size<sup>c</sup> | Intensity<sup>c</sup> |
| L2/3 | 372.9 ± 7.5 | 41.8 ± 1.1 | 377.0 ± 11.0 | 43.8 ± 1.8 | 7.5 × 10<sup>-1</sup> | 2.9 × 10<sup>-1</sup> |
| L5 | 337.5 ± 21.4 | 36.0 ± 1.7 | 481.6 ± 16.2 | 42.0 ± 1.5 | 8.6 × 10<sup>-5</sup> | 5.9 × 10<sup>-2</sup> |
| L6 | 318.2 ± 10.3 | 35.9 ± 0.9 | 475.8 ± 24.3 | 50.6 ± 2.3 | 8.8 × 10<sup>-10</sup> | 1.1 × 10<sup>-9</sup> |

**Supplementary table S37.**
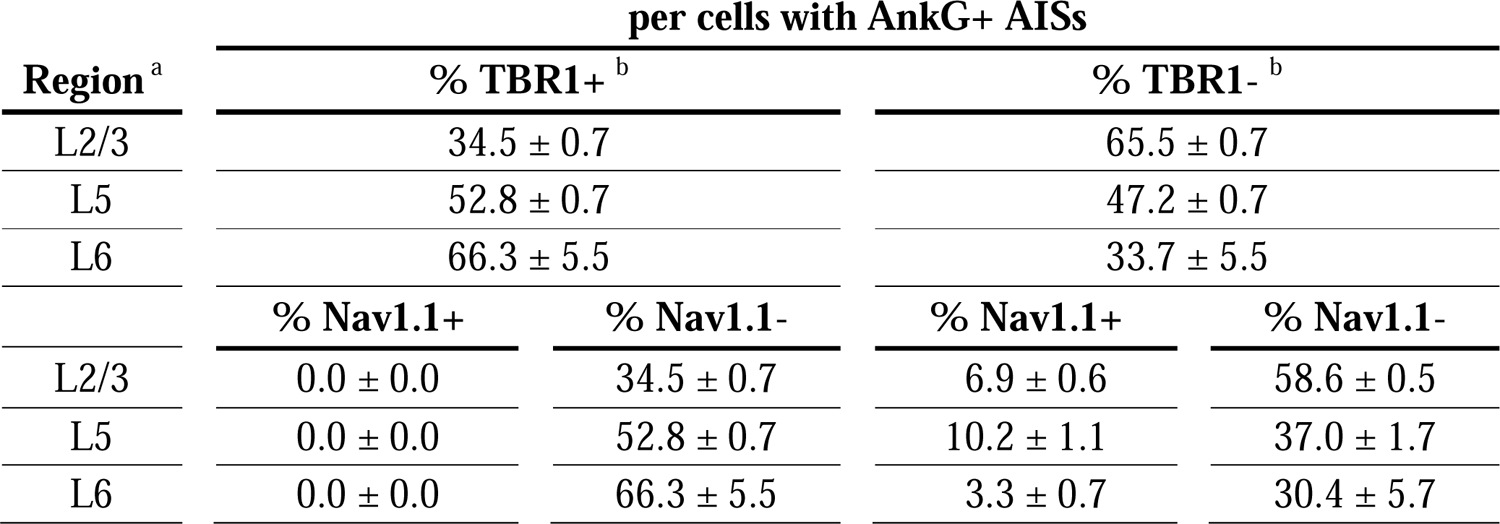
Percentage of cells with TBR1- and Nav1.1-positive/negative nuclei or AISs per cells with ankyrinG-positive AISs in *Scn1a*-GFP mouse neocortex. Values for Supplementary figure S11B-left panel. ^a^L2/3, L5, L6: neocortical layer II/III, V, VI. Cells in three *Scn1a*-GFP mice (line #233) at P15 were counted; ^b^L2/3 (N = 879 cells), L5 (N = 614 cells) and L6 (N = 816 cells). Values are presented as mean ± SEM. AnkG, ankyrinG; +, positive; -, negative.

**Supplementary table S38.** Percentage of cells with TBR1-positive/negative nuclei per cells with ankyrinG/Nav1.1-double positive AISs in *Scn1a*-GFP mouse neocortex. Values for Supplementary figure S11B-middle panel. ^a^L2/3, L5, L6: neocortical layer II/III, V, VI. Cells in three *Scn1a*-GFP mice (line #233) at P15 were counted; ^b^L2/3 (N = 59 cells), L5 (N = 63 cells) and L6 (N = 28 cells). Values are presented as mean ± SEM. AnkG, ankyrinG; +, positive; -, negative.

| Region<sup>a</sup> | per cells with AnkG+ and Nav1.1+ AISs |  |
| --- | --- | --- |
|  | % TBR1+<sup>b</sup> | % TBR1-<sup>b</sup> |
| L2/3 | 0.0 ± 0.0 | 100.0 ± 0.0 |
| L5 | 0.0 ± 0.0 | 100.0 ± 0.0 |
| L6 | 0.0 ± 0.0 | 100.0 ± 0.0 |

**Supplementary table S39.** Percentage of cells with Nav1.1-positive/negative AISs per cells with ankyrinG-positive AISs and TBR1-positive nuclei in *Scn1a*-GFP mouse neocortex. Values for Supplementary figure S11B-right panel. ^a^L2/3, L5, L6: neocortical layer II/III, V, VI. Cells in three *Scn1a*-GFP mice (line #233) at P15 were counted; ^b^L2/3 (N = 296 cells), L5 (N = 319 cells) and L6 (N = 552 cells). Values are presented as mean ± SEM. AnkG, ankyrinG; +, positive; -, negative.

| Region <sup>a</sup> | per cells with AnkG+ AISs and TBR1+ nuclei |  |
| --- | --- | --- |
|  | % Nav1.1+ <sup>b</sup> | % Nav1.1- <sup>b</sup> |
| L2/3 | 0.0 ± 0.0 | 100.0 ± 0.0 |
| L5 | 0.0 ± 0.0 | 100.0 ± 0.0 |
| L6 | 0.0 ± 0.0 | 100.0 ± 0.0 |

**Supplementary table S40.** Percentage of cells with TBR1- and Nav1.2-positive/negative nuclei or AISs per cells with ankyrinG-positive AISs in *Scn1a*-GFP mouse neocortex. Values for Supplementary figure S12B-left-upper panel. ^a^L2/3, L5, L6: neocortical layer II/III, V, VI. Cells in three *Scn1a*-GFP mice (line #233) at P15 were counted; ^b^L2/3 (N = 967 cells), L5 (N = 667 cells), and L6 (N = 833 cells). Values are presented as mean ± SEM. AnkG, ankyrinG; +, positive; -, negative.

| Region <sup>a</sup> | per cells with AnkG+ AISs |  |  |  |
| --- | --- | --- | --- | --- |
|  | % TBR1+ <sup>b</sup> |  | % TBR1- <sup>b</sup> |  |
| L2/3 | 33.3 ± 5.0 |  | 66.7 ± 5.0 |  |
| L5 | 48.9 ± 2.5 |  | 51.1 ± 2.5 |  |
| L6 | 62.7 ± 3.5 |  | 37.3 ± 3.5 |  |
|  | % Nav1.2+ | % Nav1.2- | % Nav1.2+ | % Nav1.2- |
| L2/3 | 22.6 ± 2.2 | 10.7 ± 2.9 | 57.4 ± 3.5 | 9.3 ± 1.6 |
| L5 | 34.6 ± 1.6 | 14.3 ± 1.0 | 31.3 ± 0.9 | 19.8 ± 2.6 |
| L6 | 42.4 ± 1.2 | 20.3 ± 3.3 | 26.5 ± 4.8 | 10.8 ± 2.2 |

**Supplementary table S41.** Percentage of cells with TBR1-positive/negative nuclei per cells with ankyrinG/Nav1.2-double positive AISs in *Scn1a*-GFP mouse neocortex. Values for Supplementary figure S12B-middle-upper panel. ^a^L2/3, L5, L6: neocortical layer II/III, V, VI. Cells in three *Scn1a*-GFP mice (line #233) at P15 were counted; ^b^L2/3 (N = 995 cells), L5 (N = 667 cells) and L6 (N = 923 cells). Values are presented as mean ± SEM. AnkG, ankyrinG; +, positive; -, negative.

| Region<sup>a</sup> | per cells with AnkG+ and Nav1.2+ AISs |  |
| --- | --- | --- |
|  | % TBR1+<sup>b</sup> | % TBR1-<sup>b</sup> |
| L2/3 | 28.5 ± 3.2 | 71.5 ± 3.2 |
| L5 | 52.8 ± 1.4 | 47.2 ± 1.4 |
| L6 | 62.4 ± 4.5 | 37.6 ± 4.5 |

**Supplementary table S42.** Percentage of cells with Nav1.2-positive/negative AISs per cells with ankyrinG-positive AISs and TBR1-positive nuclei in *Scn1a*-GFP mouse neocortex. Values for Supplementary figure S12B-right-upper panel. ^a^L2/3, L5, L6: neocortical layer II/III, V, VI. Cells in three *Scn1a*-GFP mice (line #233) at P15 were counted; ^b^L2/3 (N = 317 cells), L5 (N = 309 cells) and L6 (N = 523 cells). Values are presented as mean ± SEM. AnkG, ankyrinG; +, positive; -, negative.

| Region<sup>a</sup> | per cells with AnkG+ AISs and TBR1+ nuclei |  |
| --- | --- | --- |
|  | % Nav1.2+<sup>b</sup> | % Nav1.2-<sup>b</sup> |
| L2/3 | 69.3 ± 3.5 | 30.8 ± 3.5 |
| L5 | 68.9 ± 2.7 | 31.1 ± 2.7 |
| L6 | 68.7 ± 3.8 | 31.3 ± 3.8 |

**Supplementary table S43.** Percentage of cells with AnkG-positive/negative AISs per cells with TBR1-positive nuclei in *Scn1a*-GFP mouse neocortex. Values for Supplementary figure S12B-lower panel. ^a^L2/3, L5, L6: neocortical layer II/III, V, VI. Cells in three *Scn1a*-GFP mice (line #233) at P15 were counted; ^b^L2/3 (N = 509 cells), L5 (N = 582 cells) and L6 (N = 998 cells). Values are presented as mean ± SEM. AnkG, ankyrinG; +, positive; -, negative.

| Region <sup>a</sup> | per cells with TBR1+ nuclei |  |
| --- | --- | --- |
|  | % AnkG+ <sup>b</sup> | % AnkG- <sup>b</sup> |
| L2/3 | 62.9 ± 4.9 | 37.1 ± 4.9 |
| L5 | 52.2 ± 5.5 | 47.8 ± 5.5 |
| L6 | 52.3 ± 3.4 | 47.7 ± 3.4 |

**Supplementary table S44.** Percentage of cells with GFP- and Nav1.2-positive/negative somata and AISs per cells with TBR1-positive nuclei in *Scn1a*-GFP mouse neocortex. Values for Supplementary figure S13B. ^a^L2/3, L5, L6: neocortical layer II/III, V, VI. Cells in three *Scn1a*-GFP mice (line #233) at P15 were counted; ^b^L2/3 (N = 461 cells), L5 (N = 464 cells) and L6 (N = 913 cells). ^c^In these values, because they were including AnkG-negative cells, to obtain correct cell population for Nav1.2-/AnkG+ cells, virtual cell numbers were estimated using ratio of AnkG+/TBR1+ cells in Supplementary figure S12B. Values are presented as mean ± SEM. AnkG, ankyrinG; +, positive; -, negative.

| Region <sup>a</sup> | per cells with TBR1+ nuclei |  |  |  |
| --- | --- | --- | --- | --- |
|  | % GFP+ <sup>b</sup> |  | % GFP- <sup>b</sup> |  |
| L2/3 | 57.7 ± 5.0 |  | 42.3 ± 5.0 |  |
| L5 | 29.7 ± 6.0 |  | 70.3 ± 6.0 |  |
| L6 | 11.6 ± 2.5 |  | 88.4 ± 2.5 |  |
|  | % Nav1.2+ <sup>c</sup> | % Nav1.2- <sup>c</sup> | % Nav1.2+ <sup>c</sup> | % Nav1.2- <sup>c</sup> |
| L2/3 | 30.2 ± 6.8 | 27.4 ± 2.5 | 20.4 ± 3.6 | 22.0 ± 2.7 |
| L5 | 12.8 ± 5.6 | 16.3 ± 2.6 | 36.8 ± 5.4 | 34.1 ± 1.8 |
| L6 | 5.0 ± 1.3 | 7.0 ± 2.3 | 36.6 ± 5.2 | 51.5 ± 2.8 |

